# Mechanistic insights into intramembrane proteolysis by *E. coli* site-2 protease homolog RseP

**DOI:** 10.1101/2022.01.31.478169

**Authors:** Yuki Imaizumi, Kazunori Takanuki, Takuya Miyake, Mizuki Takemoto, Kunio Hirata, Mika Hirose, Rika Oi, Tatsuya Kobayashi, Kenichi Miyoshi, Rie Aruga, Tatsuhiko Yokoyama, Shizuka Katagiri, Hiroaki Matsuura, Kenji Iwasaki, Takayuki Kato, Mika K. Kaneko, Yukinari Kato, Michiko Tajiri, Satoko Akashi, Osamu Nureki, Yohei Hizukuri, Yoshinori Akiyama, Terukazu Nogi

## Abstract

Site-2 proteases are a conserved family of intramembrane proteases that cleave transmembrane substrates to regulate signal transduction and maintain proteostasis. Here, we elucidated crystal structures of inhibitor-bound forms of bacterial site-2 proteases including E. coli RseP. Our observations are consistent with a rearrangement of the RseP domains surrounding the active center to expose the substrate-binding site where a conserved electrostatic linkage between the transmembrane and membrane-associated domains mediates the conformational changes, suggesting that RseP has a gating mechanism to regulate substrate entry. Mutational analysis also supports that the substrate transmembrane helix is unwound by strand addition to the intramembrane β sheet and is clamped at the active center for efficient cleavage. Furthermore, this substrate accommodation mechanism appears to be common across distinct intramembrane proteases.

## Introduction

Intramembrane proteolysis – hydrolysis of a peptide bond within the lipid bilayer – is implicated in a variety of cellular processes in all three domains of life, including signal transduction and membrane protein homeostasis (*1-3*). In humans, deregulation of this cleavage leads to diseases such as Alzheimer’s disease (*4, 5*) while the signal transduction through the intramembrane proteolysis is associated with pathogenic infections (*6, 7*). The cleavage of substrate transmembrane (TM) segments is catalyzed by three distinct families of intramembrane proteases, each classified based on catalytic mechanism: the zinc metalloprotease site-2 protease (S2P), the aspartic protease presenilin/signal peptide peptidase (SPP), and the serine protease Rhomboid (*8, 9*). Eukaryotic S2Ps, including human S2P, have been identified in signal transduction for lipid metabolism (*10-12*) and endoplasmic reticulum stress responses (*13*), in which they perform the intramembrane proteolysis of transcription factor precursors after the extracytoplasmic cleavage of the substrates by site-1 protease. RseP from *E. coli* is an S2P homolog classified as the same subfamily (Group I) as human S2P (*1, 14, 15*) (Fig. S1). RseP is also involved in the second step of sequential cleavage of type II membrane proteins for signal transduction. RseP cleaves the TM segment of anti-sigma factor RseA in the extracytoplasmic stress response after the periplasmic region of RseA is cleaved off by the membrane-anchored protease DegS (*16-18*) (Fig. S2). In RseP, two tandemly arranged periplasmic PDZ domains (PDZ tandem) were proposed to serve as a size-exclusion filter to sterically hinder active site entry by any substrate possessing a bulky periplasmic domain (*19-21*). This substrate discrimination by size-exclusion was also proposed for the human S2P, which possesses an extracytoplasmic PDZ domain (*22*). Besides, S2Ps are presumed to possess a β-sheet in the proximity of the active center in the TM domain commonly (*23, 24*). RseP is also predicted to possess two intramembrane β hairpins (Fig. S3), which were shown to bind the substrate near the bond that is cleaved and to contribute to substrate discrimination (*24, 25*). However, the absence of structural data hampers understanding of how site-1-cleaved substrates pass through the size-exclusion filter to access the active center and of how the substrate TM segments exactly bind with the intramembrane β-sheet of S2Ps. Because the active center of S2Ps is predicted to be located within the hydrophobic milieu of the lipid bilayer, it must both form a hydrophilic compartment around the catalytic zinc for efficient hydrolysis and also accommodate hydrophobic segments of the substrate TM domain. Concerning the S2P family, a crystal structure including the active center is available for the TM domain of the archaeon *Methanocaldococcus jannaschii* S2P homolog (*Mj*S2P) (*26*). However, *Mj*S2P belongs to the Group III subfamily of S2P and does not possess PDZ domains in the extracytoplasmic region (*15*) (Fig. S1). Furthermore, *Mj*S2P was proposed to regulate substrate entry by using the TM helices flanking the catalytic core TM domain as a gate, but those helices are not present in Group I S2Ps such as *Ec*RseP and human S2P. Therefore, further structural analysis, especially on the Group I S2Ps possessing extracytoplasmic PDZ domains, is essential for understanding the mechanism of sequential cleavage and substrate accommodation in the intramembrane proteolysis by S2Ps.

Here, we began by performing X-ray crystallographic analysis on *Ec*RseP and its ortholog to produce the first atomic models for the Group I subfamily of S2P. Specifically, we have elucidated their 3D structures in complex with a peptide-mimetic inhibitor, which aided the design of mutational analyses to examine the binding mode of substrate TM segments. Furthermore, the observed structural differences between *Ec*RseP and the ortholog prompted us to examine the possibility of domain rearrangement in *Ec*RseP during substrate accommodation and cleavage. The results provide crucial insights into the mechanism of the intramembrane proteolysis not only for S2Ps, but also for the other intramembrane protease families.

## Results

### Crystallographic analysis of *Ec*RseP and *Kk*RseP

For structural analysis, we purified RseP from *E. coli* (*Ec*RseP) and its ortholog from marine bacterium *Kangiella koreensis* (*Kk*RseP) (Fig. S3). *Kk*RseP restored the growth deficiency of *E. coli rseP* mutant cells (Fig. S4a). In *E. coli*, *Kk*RseP cleaved an analogue of its native substrate containing the TM segment from the *K. koreensis* RseA ortholog and substrate analogues for *Ec*RseP. Mutations to the putative *Kk*RseP active site impaired this cleavage activity (Fig. S4b, c). Furthermore, detergent-solubilized *Kk*RseP can cleave *E. coli* RseA, although with reduced activity as compared with *Ec*RseP (Fig. 1j). In this study, crystal structures of *Ec*RseP and SeMet-substituted *Kk*RseP were determined in complex with batimastat, an inhibitor for *Ec*RseP (*27*) (Fig. 1, Fig. S5).

**Fig. 1.**
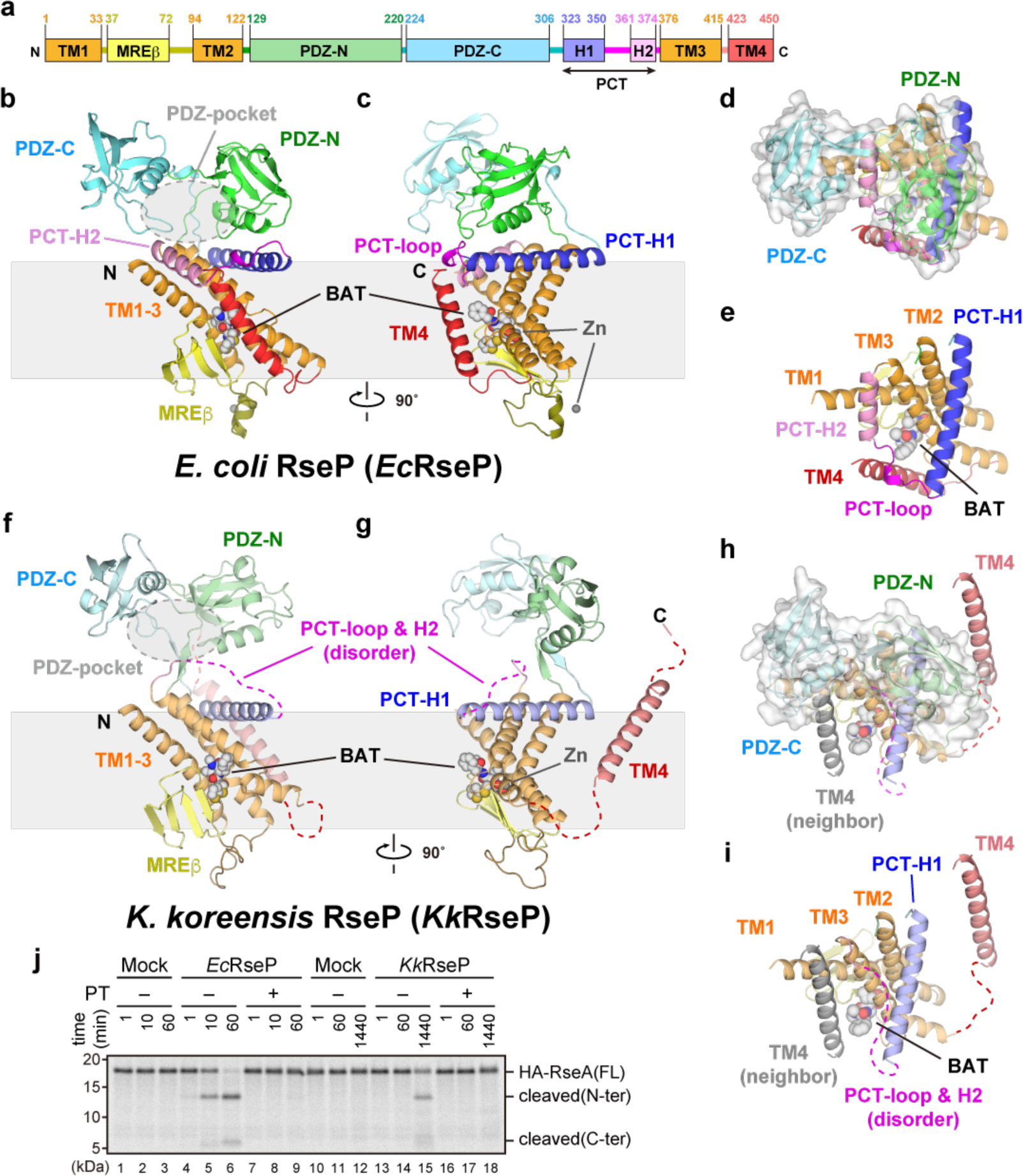
Crystal structures of *Ec*RseP and *Kk*RseP. **(a)** Domain organization of *Ec*RseP. The yellow square labeled with MREβ indicates the MREβ-sheet. The squares labeled with H1 and H2 indicate PCT-H1 and -H2, respectively. **(b, c)** Two views of the structure of full-length *Ec*RseP. Polypeptide chains are shown as ribbon models. Batimastat (BAT) and zinc ions (Zn) are shown as sphere models. Each domain or motif is colored as in **(a)**. **(d, e)** View of the structure of full-length *Ec*RseP from an outside-in perspective relative to the membrane. The PDZ tandem is highlighted with a transparent surface in **(d)** while the PDZ tandem is removed to visualize the PCT region and the helix bundle of the TM domain in **(e)**. **(f-i)** Structure of full-length *Kk*RseP in the same views as that of *Ec*RseP. In **(h)** and **(i)**, the neighboring TM4 segment accommodated into the TM1-TM3 cleft is shown in gray ribbon model. **(j)** *In vitro* substrate cleavage assay with purified RseP proteins. ^35^S-Met-labeled model substrate HA-RseA148 from cell-free synthesis was incubated at 37°C for the indicated time with 2.5 ng/μL (50 nM) *Ec*RseP, 100 ng/μL (2.0 μM) *Kk*RseP, or enzyme buffer only (mock) plus 5 mM of zinc chelator *1,10*-phenanthroline (PT+) or 5% DMSO (PT-) as indicated. Model substrate and cleavage products (labels, right) on a Bis-Tris SDS-PAGE gel were visualized using a phosphor imager. Data are representative of three technical replicates.

### Overall structure of *Ec*RseP

The final model of *Ec*RseP (M1-F447) is full length except for the three C-terminal residues and recombinant tag (Fig. 1b-e). TM1 (M1-C33) contains two zinc-coordinating His residues, H22 and H26 (Fig. S6a). TM3 is divided by a loop-like bulge into two segments, TM3-N and TM3-C. TM3-C contains the third zinc-coordinating residue, D402 (Fig. S6a). Besides the zinc ion in the active center, a second zinc ion was bound to H86 and H87 in the cytoplasmic region. However, its physiological role is currently unknown as H86A and/or H87A mutations did not affect the proteolytic activity (Fig. S6b-d). *Ec*RseP was predicted to possess two intramembrane β-hairpins, the C1N loop (*25*) and the membrane-reentrant β (MREβ) loop (*24*), between TM1 and TM2 (Fig. S3). The corresponding regions are integrated into a four-stranded β-sheet (hereafter referred to as the MREβ sheet) (Fig. S7a, b). Strand 4 corresponds to the edge strand and forms one side of the substrate-binding site where its backbone makes direct contacts with batimastat (Fig. 2). These observations are consistent with our previous findings that proteolytic activity is reduced by introducing Pro mutations into the two strands closest to the active site as observed here, either into the N-terminal region of C1N (R39-F44 corresponding to strand 1) or into the C-terminal region of the MREβ loop (G67-V70 corresponding to strand 4) (*25*). Similarly, substrate cleavage was strongly impaired in the G43A/I61G double mutant but not by either single mutation. In the crystal structure, G43 and I61 are proximal (G43 on the C1N loop and I61G on the MREβ loop). *Mj*S2P also possesses a membrane-embedded β-sheet (*26*) while its topology differs in strand order. However, strand 4 is still proximal to the active center (Fig. S7c, d), and the arrangement of the β-sheet relative to the active center is the same as for the MREβ sheet of *Ec*RseP. Taken together, TM1-3 and the MREβ sheet structurally align with their equivalents in *Mj*S2P. On the cytoplasmic side of the *Ec*RseP MREβ sheet, the presence of several basic residues likely excludes lipid molecules from the region around the MREβ sheet on the cell membrane and permits the entry of water molecules into the active center of *Ec*RseP (Fig. S7e, f).

**Fig. 2.**
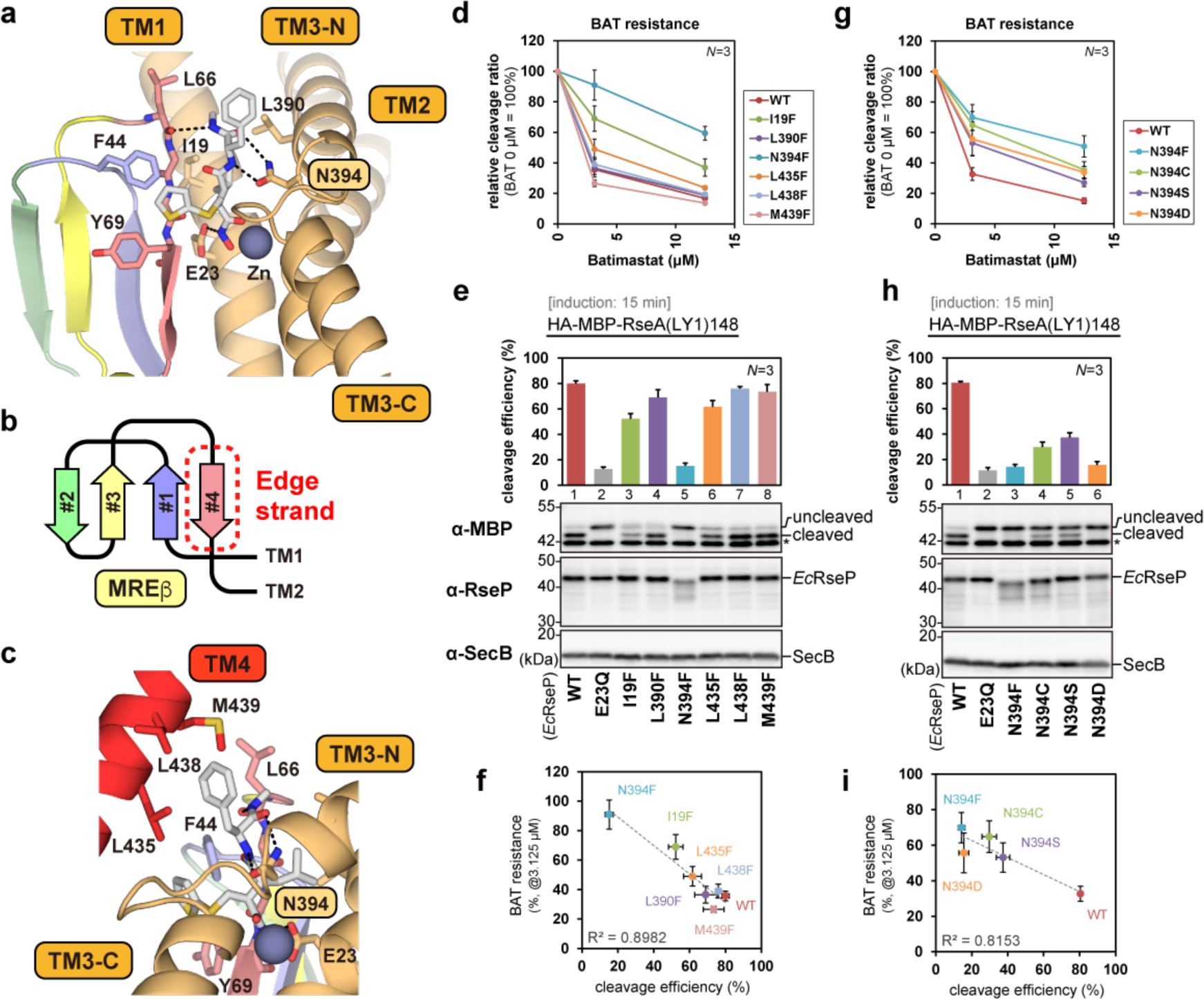
Binding mode of batimastat to *Ec*RseP and dependence on residue N394. **(a)** Close-up view of the batimastat-binding site. Batimastat and the residues in direct contact with batimastat are shown as stick models. The label is highlighted for functionally important residue N394. The four strands constituting the MREβ sheet are shown in different colors. The zinc ion in the active site is shown as a sphere model. TM4 is omitted to visualize the binding site. **(b)** Topology diagram of the MREβ sheet. **(c)** Interaction between TM4 and batimastat. The model is shown similarly to that in **(a)**. **(d, g)** *In vivo* batimastat sensitivity assay. *E. coli* YH2902 (Δ*rseA* Δ*rseP* Δ*acrA*) cells harboring one plasmid for the HA-MBP-RseA(LY1)148 model substrate (pYH124) and one for tag-less *Ec*RseP (pYH825) or its variants (labels, right), were pre-treated with 0, 12.5 or 3.125 μM of batimastat at 30°C for 10 min before inducing expression. After 1 hour induction with 1 mM IPTG, the relative cleavage ratio was determined by quantitating the signal from SDS-PAGE and immunoblots of TCA-precipitated proteins and comparing the value with that for the 0 mM BAT condition. Points and error bars represent the means ± S.D. from three biological replicates. (See Fig. S8 for the immunoblotting results and the quantified cleavage ratio in each condition.) **(e, h)** *In vivo* substrate cleavage assay with short induction. YH2902 cells carrying two plasmids as in **(d, g)** were prewarmed at 30°C for 10 min and incubated for 15 min with 1 mM IPTG to induce an *Ec*RseP variant and the model substrate. The cleavage efficiency was determined by quantitating the signal from SDS-PAGE and immunoblots of TCA-precipitated proteins (images). The cytoplasmic protein SecB serves as a loading control (α-SecB), and an asterisk indicates endogenous MBP protein. Bar plots and error bars represent the means ± S.D. from three biological replicates. **(f, i)** Scatter plots of the BAT resistances of the RseP mutants (relative cleavage ratio under 3.125 μM of BAT concentration shown in **(d)** or **(g)**) against the cleavage efficiencies of the model substrate (shown in **(e)** or **(h)**). The coefficient of determination (R^2^) is indicated on each plot.

The PDZ tandem protrudes into the periplasmic space (Fig. 1b, c). The two PDZ domains form a pocket-like space (PDZ pocket) oriented toward the TM domain containing the active center (Fig. 1b-d). As predicted in the previous study (*21*), the N-terminal residues of the PDZ C-terminal (PCT) region (P323-T350) forms an amphiphilic helix (PCT-H1) at the membrane surface (Fig. 1c). PCT-H1 leads to a loop region (G351-G360) containing a short 310 helix (PCT-SH) (Fig. 1e, Fig. S3). PCT-H1 and this loop region make direct contacts with PDZ-N. The present structural analysis also showed that the C-terminal residues of PCT (P361-L374) also form a helix (PCT-H2), which makes a sharp turn at the C-terminal end (G375) and leads to TM3-N (Fig. 1e). PCT-H2 is accommodated into a cleft formed between TM1 and TM3-N and is located just above the batimastat bound to the active center. In *Ec*RseP, TM4 (V423-F447) interacts with PCT-SH and PCT-H2 at the periplasmic side and covers the batimastat.

### Binding mode of batimastat

The peptide-mimetic batimastat adopts an extended conformation and is flanked by the edge strand of the MREβ sheet and TM3-N in *Ec*RseP (Fig. 2, Fig. S8a). The main chain of batimastat forms hydrogen bonds with the main chain of L66 on the edge strand and with the side chain of N394 on TM3. For the side chains of batimastat, the isobutyl group is oriented toward I19 in TM1 and L390 in TM3-N while the thienyl group is close to both of the residues that were reported to interact with substrates, F44 of the conserved GFG motif (*25*) and Y69 on the edge strand of the MREβ-sheet (*24*). In addition, the phenyl group of batimastat forms van der Waals interactions with L435, L438, and M439 on TM4 (Fig. 2c). Thus, TM1, 3, and 4, together with the edge strand of the MREβ sheet, form a compartment separated from the lipid bilayer that accommodates batimastat and the catalytic zinc. The interior of the compartment is hydrophilic due to the presence of several charged or polar residues (N108, S387, N394, Y428, S432, and the zinc-coordinating residues) and the exposed backbone of the edge strand. (Fig. S7e, f).

Based on the structural data, we examined if side chains could also impact inhibitor (and thereby substrate) accommodation in addition to the backbone interactions with the edge strand. I19N and I19F mutations to TM1 have been reported to reduce the sensitivity of *Ec*RseP to batimastat (*27*). We further introduced Phe mutations to the residues interacting with batimastat on TM3 and TM4 (L390 and N394 on TM3, and L435, L438, and M439 on TM4), and examined their effects on batimastat sensitivity. To eliminate the influence of the C-terminal tags on this analysis, we introduced these mutations into a tag-less *Ec*RseP construct (Fig. 2d, Fig. S8b, c). In the wild type, addition of 3.125 μM batimastat reduced the relative cleavage ratio of the substrate to 40%. At the same batimastat concentration with N394F, the relative cleavage ratio was only reduced to 90%. I19F also showed increased resistance to batimastat as reported (*27*). Furthermore, batimastat resistance seems to correlate negatively with the intrinsic proteolytic activity of the mutants in the absence of batimastat, particularly with a shorter induction of *Ec*RseP (Fig. 2e, f). The N394F mutation with the strongest impact on batimastat resistance also reduced the proteolytic activity to the same degree as the active-site mutation E23Q under the shorter induction condition. Next, we introduced N394C, N394S, and N394D mutations to evaluate the possibility that the reduced activity in N394F is ascribed to lower accumulation resulting from fold destabilization. We observed that all three mutants accumulated in *E. coli*, but still showed significant increase both in batimastat resistance and in reduction of proteolytic activity (Fig. 2g-i, Fig. S8d, e). In particular, the reduction of proteolytic activity for the isosteric mutation N394D indicates that hydrogen bonds via the amide group of the N394 side chain is critical for the cleavage. N394 may clamp a bound substrate at the active center as it interacted with the backbone of batimastat at the opposite side of the MREβ sheet. In total, it is likely that the binding mode of batimastat partly reflects that of native RseP substrates. The substrate segments to be cleaved are thought to be extended by the strand addition and shielded from the hydrophobic milieu of the lipid bilayer by the surrounding TM helices of RseP.

### Overall structure of *Kk*RseP

*Kk*RseP produced two crystal forms with similar crystal packing (Fig. S9a-d). The two crystal structures of *Kk*RseP are almost identical with an RMSD of 1.34 Å for 387 Cα atoms (Fig. S9e, f), excluding some disordered loops. The structures of individual domains in *Kk*RseP are similar to those in *Ec*RseP, including the binding mode of batimastat via the MREβ sheet and sidechain-backbone interactions with N387 (corresponding to N394 in *Ec*RseP) (Fig. 1f-i, Fig. S10). However, the *Kk*RseP structures showed significant differences in the domain arrangement relative to those in *Ec*RseP. For instance, the PDZ tandem is positioned further from the PCT region, and the C-terminal part of PCT (corresponds to PCT-SH and H2 in *Ec*RseP) is disordered (Fig. 1f, g, i). Thus, *Kk*RseP adopts a PDZ-open conformation while *Ec*RseP is in a PDZ-closed conformation. In addition, TM4 moves away from the domains forming the active center to interact with the cleft between TM1 and TM3-N in the crystal packing neighbor (Fig. 1h, i, Fig. S11a). TM4 also interacts with the batimastat bound to the active center of the neighbor. The phenyl group of batimastat makes close contacts with V428 and L429 on TM4 (Fig. S10b). Additionally, residual electron density was observed close to PDZ-C. This electron density is most likely derived from the C-terminal residues of the TEV protease consensus sequence in the *Kk*RseP construct (Fig. S11a, b). It is probable that the accommodation of TM4 into the neighbor is an artifact of the crystal packing, but the conformational difference between *Ec*RseP and *Kk*RseP raised the possibility that the PDZ tandem, the C-terminal part of PCT, and TM4 can rearrange without disrupting the TM core region. It is highly possible that such structural changes, if they occur, would impact the regulation of substrate accommodation.

### Arrangement of the PDZ tandem in *Ec*RseP and *Kk*RseP on the cell membrane

To explore the possibility of domain rearrangement, we first examined the arrangement of the PDZ tandem in *Ec*RseP and *Kk*RseP on the cell membrane using a mal-PEG accessibility assay. For *Ec*RseP, we again used the tag-less construct to eliminate the impact of the C-terminal tags on the conformation of the PDZ tandem. We introduced four single Cys mutations to the PDZ tandem and observed that the Cys in A136C (located outside the PDZ pocket) was efficiently modified with mal-PEG (∼5 kDa). In contrast, residues close to the PCT region (D163C and L167C) and inside the pocket (I304C) were modified only upon the addition of detergent (Fig. S12a-c). These results indicate that *Ec*RseP on the membrane adopts the PDZ-closed conformation as in the crystal structure where the PDZ pocket is sterically hindered by the bilayer and the PCT region. Next, we also examined the conformation of the PDZ tandem in *Kk*RseP on the cell membrane. To conduct the accessibility assay on *Kk*RseP without a C-terminal tag, we inserted an exogenous epitope, PA14 tag (*28*), into a β-turn between K54 and H55 in the MREβ sheet for antibody-labeling. We confirmed that the resulting *Kk*RseP(54-PA14-55) and its four single Cys derivatives (P136C, E163C, F167C, and I302C) accumulated in *E. coli* and maintained proteolytic activity (Fig. S13). If *Kk*RseP on the membrane adopts the PDZ-open conformation as in the crystal structure, all of the four Cys mutants should be modified to some extent in the spheroplast. Nevertheless, although E163C (proximal to PCT) underwent significant but rather low modification, F167C (proximal to PCT) and I302C (inside the PDZ pocket) were unmodified in the absence of detergent (Fig. S12d-f). These results suggest that the two modification sites (F167 and I302) were inaccessible due to closer proximity between the PDZ tandem and the PCT region in *Kk*RseP on the membrane, similar to what is observed in the crystal structure of *Ec*RseP (PDZ-closed) rather than that of *Kk*RseP (PDZ-open) (Fig. S12g). Thus, the PDZ-open conformation of *Kk*RseP in the crystal may be induced by the rearrangement of the PCT region or by the accommodation of TM4 from the crystal packing neighbor.

### Importance of D446 on TM4 for substrate cleavage

We next explored the role of TM4 in substrate cleavage. Although TM4 is less conserved compared to the other three TM regions within the S2P family (*15*) (Fig. 3a), the binding mode of batimastat in *Ec*RseP suggests that TM4 contributes to the formation of the hydrophilic compartment around the active site. Hence, we prepared two *Ec*RseP mutants, ΔTM4 lacking the entire TM4 and C-terminal tail region (F426 to the C-terminus) and ΔCTail lacking only the C-terminal tail region (D446 to the C-terminus), and assessed the effect of the mutations on the cleavage. We observed that both ΔTM4 and ΔCTail did not complement the *Ec*RseP-deficiency (Fig. 3b). Furthermore, both mutants virtually lost proteolytic activity where only ΔCTail exhibited residual activity (Fig. 3c). We next mutated conserved residues on TM4 to alanine (for G431, M439, D446, and R449) or serine (for A442) (Fig. 3a, d, Fig. S14) to examine their contributions to the cleavage. We found that only the mutation to the highly conserved D446 (Fig. S3) impaired complementation and proteolytic activity. As the carboxyl side chain of D446 is positioned for interaction with the electropositive N-terminal end of the helix dipole on PCT-H2 in the crystal structure, the interaction between D446 and PCT-H2 may be critical for the maintenance of the proteolytic activity (Fig. 3e). In fact, analysis of additional D446 mutants showed that only D446E retained proteolytic activity (Fig. 3c, Fig. S15), although its complementation activity was much lower than that of the wild-type (Fig. 3b). Notably, despite the absence of D446 in ΔCTail, this mutant exhibited a significant proteolytic activity. This may be due to the presence of the terminal carboxyl group of N445 in the proximity of the N-terminal end of helix dipole. In total, the above mutational analysis revealed a previously unrecognized importance to substrate cleavage for TM4 and that an Asp at position 446 is most ideal for this function in the native context. One possibility is that the electrostatic interaction between PCT-H2 and D446 is structurally important for the formation of the substrate-binding compartment. Another possibility is that this interaction is important because substrate accommodation requires the coordinated structural rearrangement of the PCT-H2 and TM4 observed between the *Ec*RseP and *Kk*RseP structures.

**Fig. 3.**
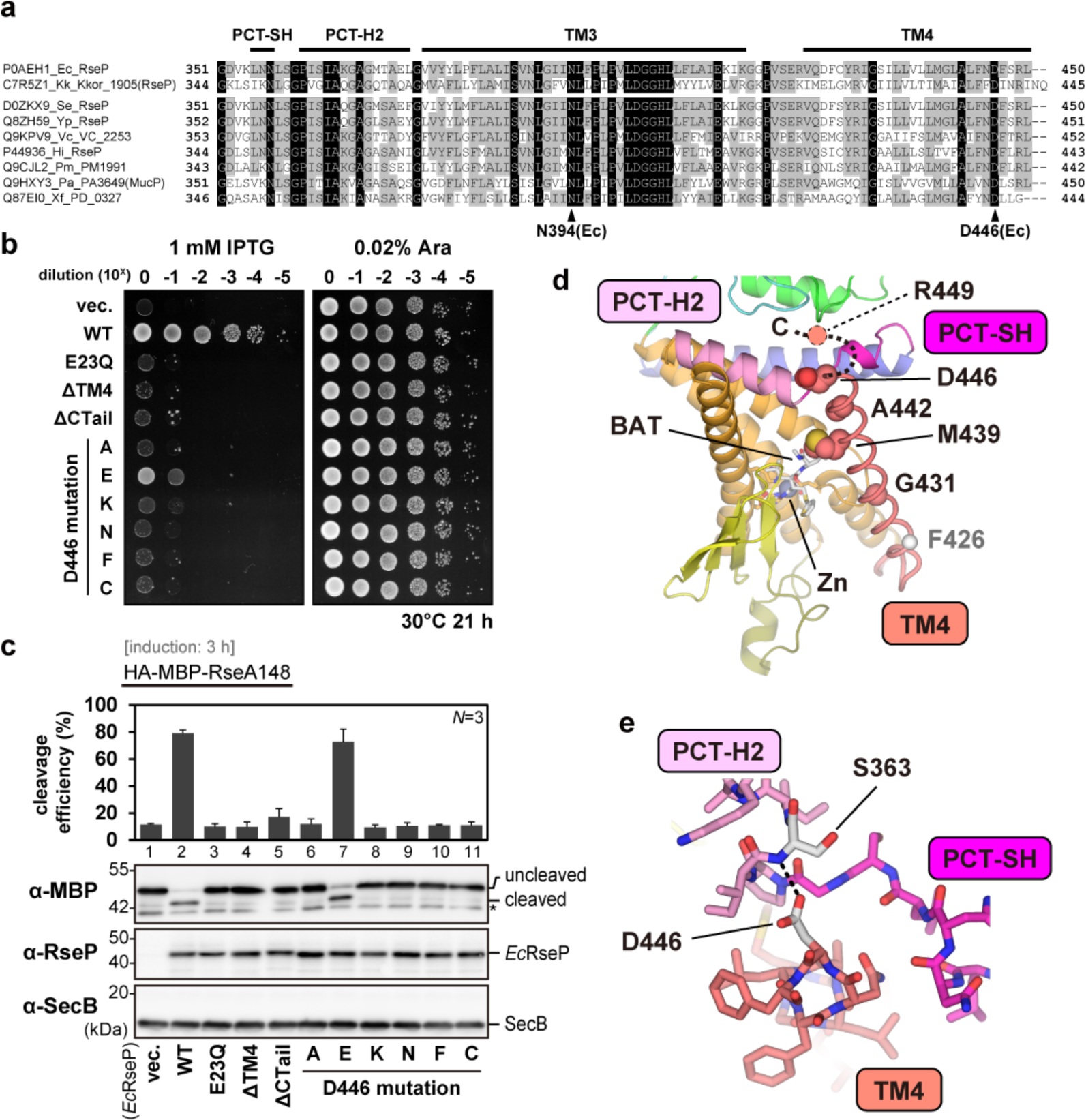
Involvement of TM4 in the substrate cleavage and dependence on residue D446. **(a)** Amino acid sequence alignment of the C-terminal regions of nine RseP orthologues classified as *γ*-proteobacteria. The UniProtKB accession number of each orthologue is shown (Ec, *Escherichia coli*; Kk, *Kangiella koreensis*; Se, *Salmonella enterica* serovar Typhimurium; Yp, *Yersinia pestis*; Vc, *Vibrio cholerae*; Hi, *Haemophilus influenzae*; Pm, *Pasteurella multocida*; Pa, *Pseudomonas aeruginosa*; Xf, *Xylella fastidiosa*). Conserved and similar residues are boxed in black and gray, respectively. The positions of TM segments, structural elements and two featured residues are shown based on those of *Ec*RseP. **(b)** Complementation assay. Cultures of *E. coli* KK31 [Δ*rseP*/pKK6 (PBAD-*rseP*)] cells carrying plasmids for *Ec*RseP (pYH825) or its variants were serially diluted and spotted on L agar plates containing IPTG (left) to test complementation or containing L-arabinose (right) as a control for total-cell count. A representative result from three biological replicates is shown. ΔTM4 and ΔCTail indicate the F426amber and D446amber mutations, respectively. **(c)** *In vivo* cleavage assay with long induction. *E. coli* KA306 (Δ*rseA* Δ*rseP* Δ*clpP*) cells harboring one plasmid for HA-MBP-RseA148 (pKA65) and one for *Ec*RseP (pYH825) or its variants were grown at 30°C for 3 h with 1 mM IPTG and 1 mM cAMP in M9-based medium. The cleavage efficiency was determined from immunoblots (images) as in Fig **2e**. Bar plots and error bars represent the means ± S.D. from three biological replicates. **(d)** Conserved residues on TM4. The residues shown as sphere models were mutated to alanine (A442 was mutated to serine) to examine their roles in the proteolytic activity, as shown in Fig. S14. Another mutated residue, R449, was disordered in the electron density map as indicated by dotted lines. The ΔTM4 mutant in **(b)** lacks the C-terminal residues downstream of F426 (white sphere) while ΔCTail lacks the periplasmic tail region (D446 to the C-terminus). Batimastat and active site residues are shown as stick models. The zinc ion is shown as a sphere. **(e)** Specific interaction between D446 and PCT-H2. TM4 and the PCT-loop, including PCT-SH and PCT-H2, are shown as stick models in magenta, pink, and salmon, respectively. D446 interacts with the electropositive N-terminal end of the PCT-H2 helix dipole where the side chain of D446 and the main chain N-H group of S363 (each in white) form a hydrogen bond.

### Involvement of PCT-H2 in the substrate cleavage

To examine if the conformational rearrangement of PCT-H2 is involved in the substrate cleavage, we first performed Cys-scanning mutagenesis on PCT-H2 and the adjacent region (T350-P381) to examine the mobility of PCT-H2 by a modified version of our accessibility assay with 4-acetamido-4′-maleimidylstilbene-2,2′-disulfonate (AMS) as the probe. AMS, a soluble thiol-alkylating reagent, reacts with Cys residues exposed to the aqueous milieu. We observed that all the Cys residues on PCT-H2 were modified at comparable levels in the spheroplast (Fig. S16), although some residues, such as P361, I364, A365, A368, G369, and A372, are expected to be modified less frequently if PCT-H2 is fixed in the cleft between TM1 and 3 as in the crystal structure. Hence, PCT-H2 is thought to be mobile in *Ec*RseP on the cell membrane. We also examined the proteolytic activity of the Cys mutants and found that only G360C and G375C reduced the proteolytic activity (Fig. 3a, 4a, c, Fig. S17). Considering the high conservation of the two Gly residues flanking both ends of PCT-H2, these results suggest that the proper positioning of PCT-H2 is important for the substrate cleavage.

**Figure 4.**
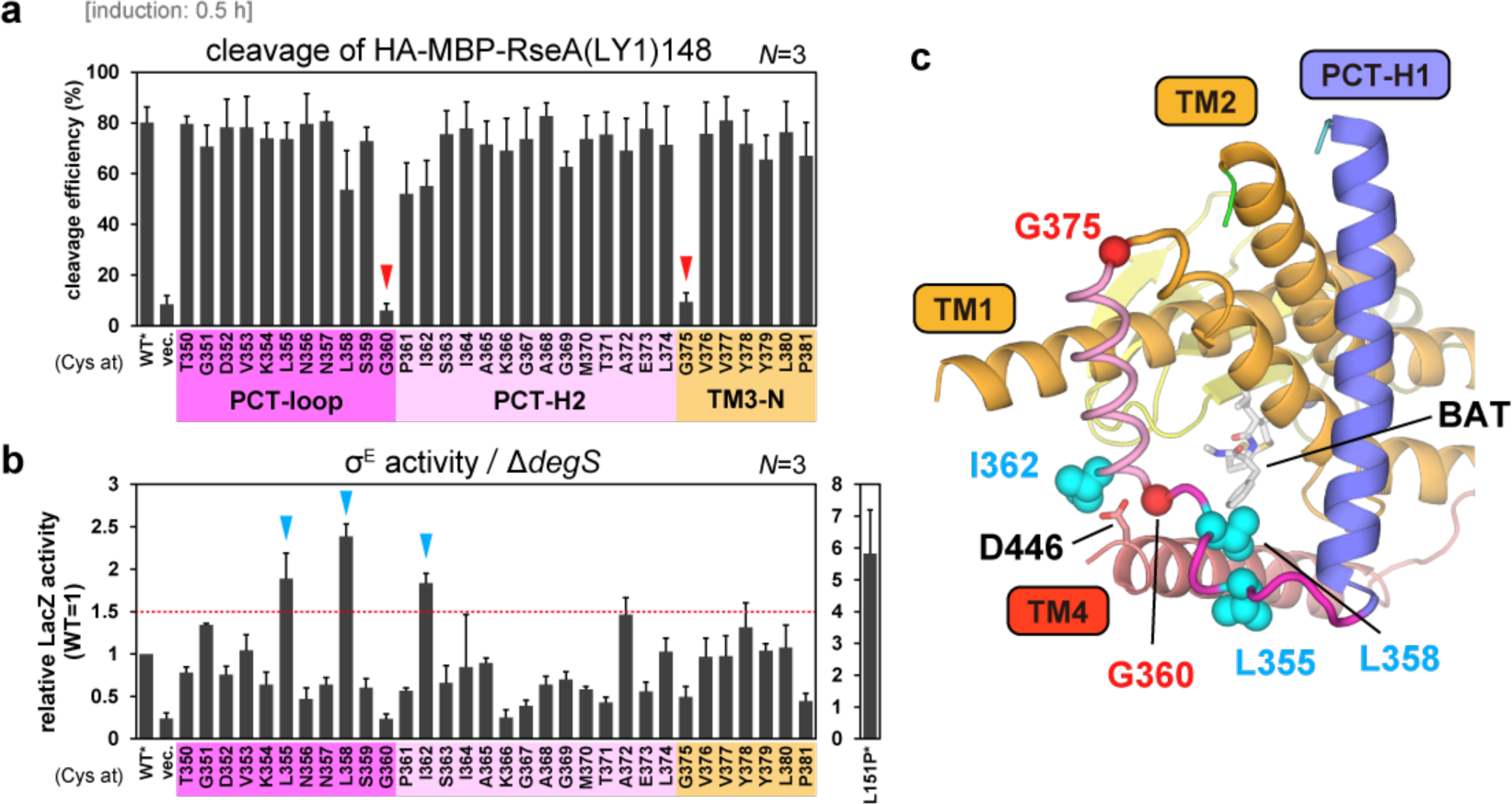
Cysteine-scanning mutagenesis analysis of the PCT-H2 and the adjacent regions. **(a)** Model substrate cleavage with short induction. *E. coli* KK211 (Δ*rseA* Δ*rseP*) cells harboring one plasmid for the HA-MBP-RseA(LY1)148 model substrate (pYH20) and one for a variant of *Ec*RseP(Cys-less)-His6-Myc (pTM132) were grown at 30°C in M9-based medium for 2.5 h and further incubated for 0.5 h with 1 mM IPTG and 5 mM cAMP. The cleavage efficiency was determined from immunoblots as in Fig **2e**. (See Fig. S17 for the immunoblotting results.) Bar plots and error bars represent the means ± S.D. from three biological replicates. The region of each mutation is indicated by the color (magenta, pink, gold) and a label. **(b)** DegS-independent σ^E^ activity of cells expressing RseP Cys mutants. Cells of *rpoH*P3*-lacZ* reporter strain AD2473 (Δ*degS* Δ*rseP*) harboring pSTD343 (*lacI*) and a plasmid for a variant of *Ec*RseP(Cys-less)-His6-Myc (pTM101) were grown at 30°C for 5 h in L medium containing 0.1 mM IPTG and 1 mM cAMP. The measured LacZ activities are normalized as the ratio to the activity for the reporter strain expressing wild type RseP (WT). The bar plot shows the means ± S.D. from three biological replicates. Red dashed line indicates the threshold for deregulation. The previously isolated L151P mutant (right) shows high LacZ activity characteristic of deregulation. WT* and L151P* indicate Cys-less derivatives of WT and L151P RseP, respectively. **(c)** Mapping mutations on the *Ec*RseP model. Residues where the Cys mutation impaired the proteolytic activity are indicated with red sphere models. Residues where the Cys mutations caused deregulation in the sequential cleavage of RseA are indicated with cyan sphere models.

Subsequently, we also tested if PCT-H2 is involved in the regulation of the sequential cleavage. We performed a LacZ reporter assay on the Cys mutants to monitor σ^E^ activation resulting from deregulated cleavage of intact RseA without the prior site-1 cleavage by DegS. We found that L355C, L358C, and I362C deregulate *Ec*RseP although the extent of deregulation was smaller than that from L151P, a known deregulated mutant on PDZ-N (*29*) (Fig. 4b, c). We also examined trypsin susceptibility of the three mutants on the cell membrane as a proxy for stability. As reported previously, L151P showed increased trypsin susceptibility because this mutation disrupts the folding of the PDZ-N. In contrast, almost no degradation was observed for the other mutants or for WT RseP (Fig. S18). We also observed increased conformational flexibility in the PDZ-C domain of L358C using negative-stain electron microscopy (EM) (Figs. S19, S20), suggesting that the L358C mutation can deregulate *Ec*RseP by altering the arrangement of the PDZ tandem without destabilizing the individual PDZ domains. These results indicate that the proper positioning and mobility of PCT-H2 are also important for the regulated sequential cleavage of substrates and likely impact the positioning of TM4.

### Conformational changes of the PCT region and PDZ tandem during substrate cleavage

Several lines of evidence from this study raised the possibility that the PDZ tandem and PCT region, probably in conjunction with TM4, undergo structural changes to accommodate the substrate into the active center for cleavage. Our previous *in vivo* photo-cross-linking experiment using *p*-benzoyl-L-phenylalanine (*p*BPA) indicated that buried residue T341 on PCT-H1 is accessible to RseA (*21*). We therefore performed an intramolecular cross-linking experiment to test if the proteolytic activity of *Ec*RseP is affected by immobilizing the PCT region and/or the PDZ tandem. In this assay, *Ec*RseP mutants possessing two Cys mutations were first expressed under IPTG induction, then the cells were washed and disulfide cross-linking was induced with an oxidizing agent, diamide. As a control, we performed the same procedure but replaced the diamide with a reducing agent, DTT. Finally, a model substrate was induced with arabinose, and cleavage was monitored with an immunoblot assay. The formation of intramolecular cross-links was confirmed by chemically modifying the unlinked Cys residues (Fig. S21). We observed that cross-linking PCT-H2 with TM1 (D7-K366 and S10-G369) reduced the proteolytic activity (Fig. 5a-c), supporting the model that PCT-H2 moves out of the cleft between TM1 and TM3-N to accommodate the substrate. Cross-linking PDZ-N with PCT-H1 (D171-S343 and I173-K347) or with the PCT-loop (E203-K354 and D205-S359) (Fig. 5a-c) also reduced the proteolytic activity. These results suggest that the PDZ tandem in *Ec*RseP also moves away from the membrane as observed in the crystal structure of *Kk*RseP. In *Kk*RseP, the cleft between TM1 and TM3-N accommodates the TM4 from the crystal packing neighbor. We inferred that the bound TM4 might mimic the substrate and tested this prediction in an intermolecular photo-cross-linking experiment with *Ec*RseP and RseA. As anticipated, *p*BPA introduced at residue 378 on the cleft is accessible to RseA *in vivo* (Fig. 5a, d, Fig. S22). Collectively, these results support a model in which the PDZ tandem and the PCT region rearrange to accommodate substrate for cleavage.

**Fig. 5.**
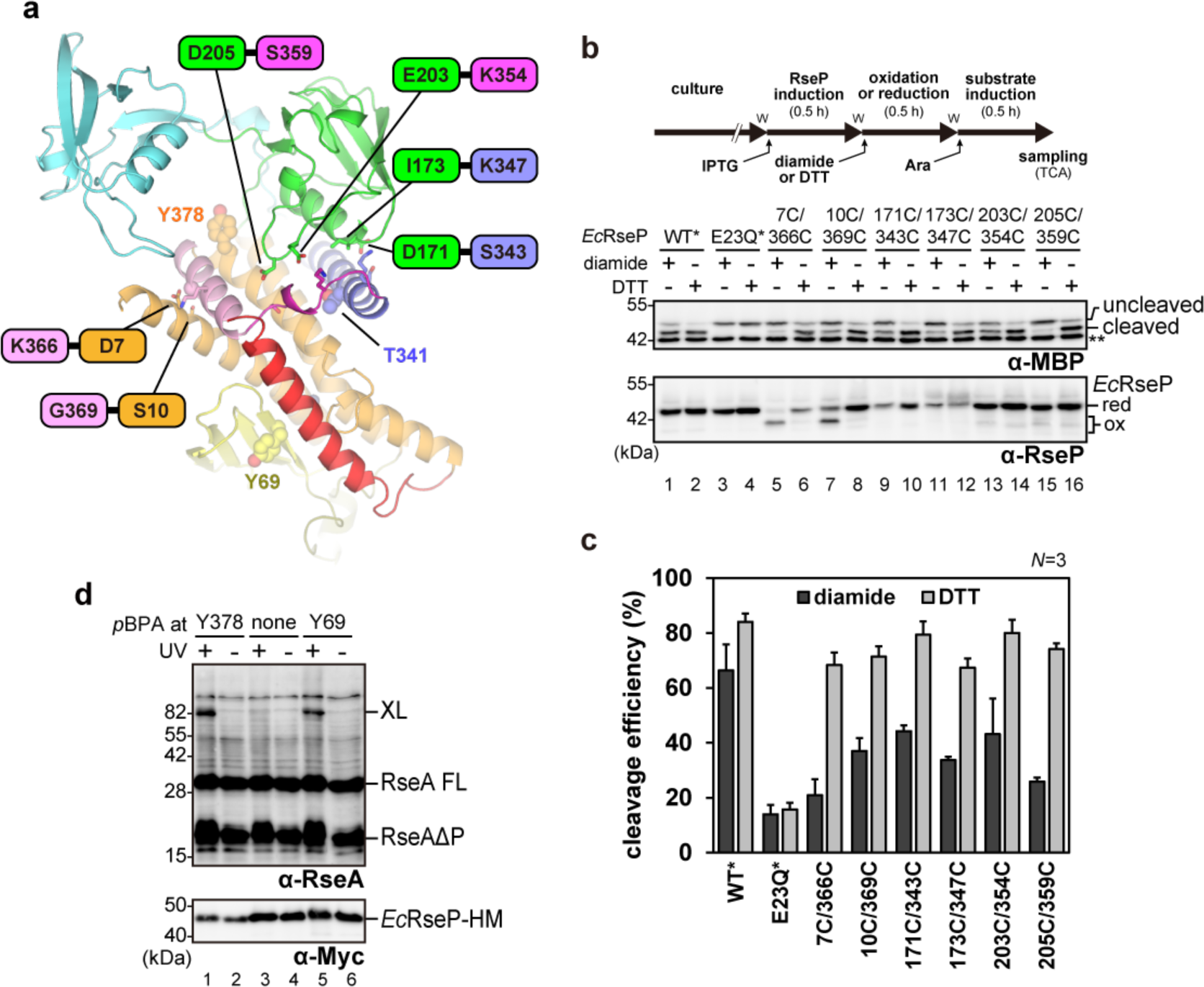
Cross-linking experiments to examine the structural change during substrate cleavage. **(a)** Introduction of intramolecular cross-links. Six pairs of residues where Cys mutations were introduced are indicated by a label and their side chains are shown as stick models (G369 is indicated by a sphere at the Cα atom). Residues where the introduced *p*BPA is cross-linked with RseA are shown with sphere models. **(b, c)** *In vivo* proteolytic activity for *Ec*RseP after domain immobilization by disulfide crosslinking. AD2544 (Δ*rseA* Δ*rseP*) cells harboring an IPTG-inducible plasmid for *Ec*RseP (pYH835) or its variants and an arabinose-inducible plasmid for the model substrate HA-MBP-RseA(LY1)148 (pTM949) were cultured at 30°C in L medium. The cultures were sequentially supplemented with 5 mM IPTG (for RseP expression) and then with 5 mM diamide (for oxidation) or 10 mM DTT (for reduction). Time-course diagram were shown at the top where ’w’ indicates a cell-wash step. After washing the cells, substrate expression and cleavage was induced for 0.5 h with 0.02% L-arabinose. The relative cleavage ratio was determined from immunoblots as in Fig **2e**. Bar plots and error bars represent the means ± S.D. from three biological replicates. WT* and E23Q* indicate Cys-less derivatives of WT and E23Q RseP, respectively. Double asterisk indicates endogenous MBP protein **(d)** *In vivo* photo-crosslinking between *Ec*RseP(Y378*p*BPA) and RseA. KA418 (Δ*ompA* Δ*ompC* Δ*rseP*, *rseA*^+^) /pEVOL-pBpF cells harboring a plasmid for RseP(E23Q)-His6-Myc (pKA52, none) or its variants having an amber mutation at the position of Y378 or Y69 were grown at 30°C in M9-based medium supplemented with 0.5 mM *p*BPA for 4 h then exposed to UV-irradiation (UV+) or withheld (UV-) for 10 min. TCA-precipitated proteins were analyzed by immunoblotting. XL indicates the cross-linked products between RseP-HM and endogenous RseA. RseA FL and RseAΔP indicate the full-length and the DegS-cleaved form of RseA, respectively. RseP(Y69*p*BPA) with *p*BPA in the MRE β-loop was used as a positive control for crosslinking with RseA. A representative result from three biological replicates is shown.

## Discussion

In this study, we successfully determined the crystal structures of *Ec*RseP and *Kk*RseP as the first experimental structures for S2Ps with extracytoplasmic PDZ domains (Group I). The structural features not only coincide with the previously proposed models, but also provide new insights into the substrate accommodation mechanism. Specifically, the substrate discrimination and accommodation by *Ec*RseP are presumed to be regulated by three processes: size-exclusion, gating, and unwinding (Fig. 6).

**Fig. 6.**
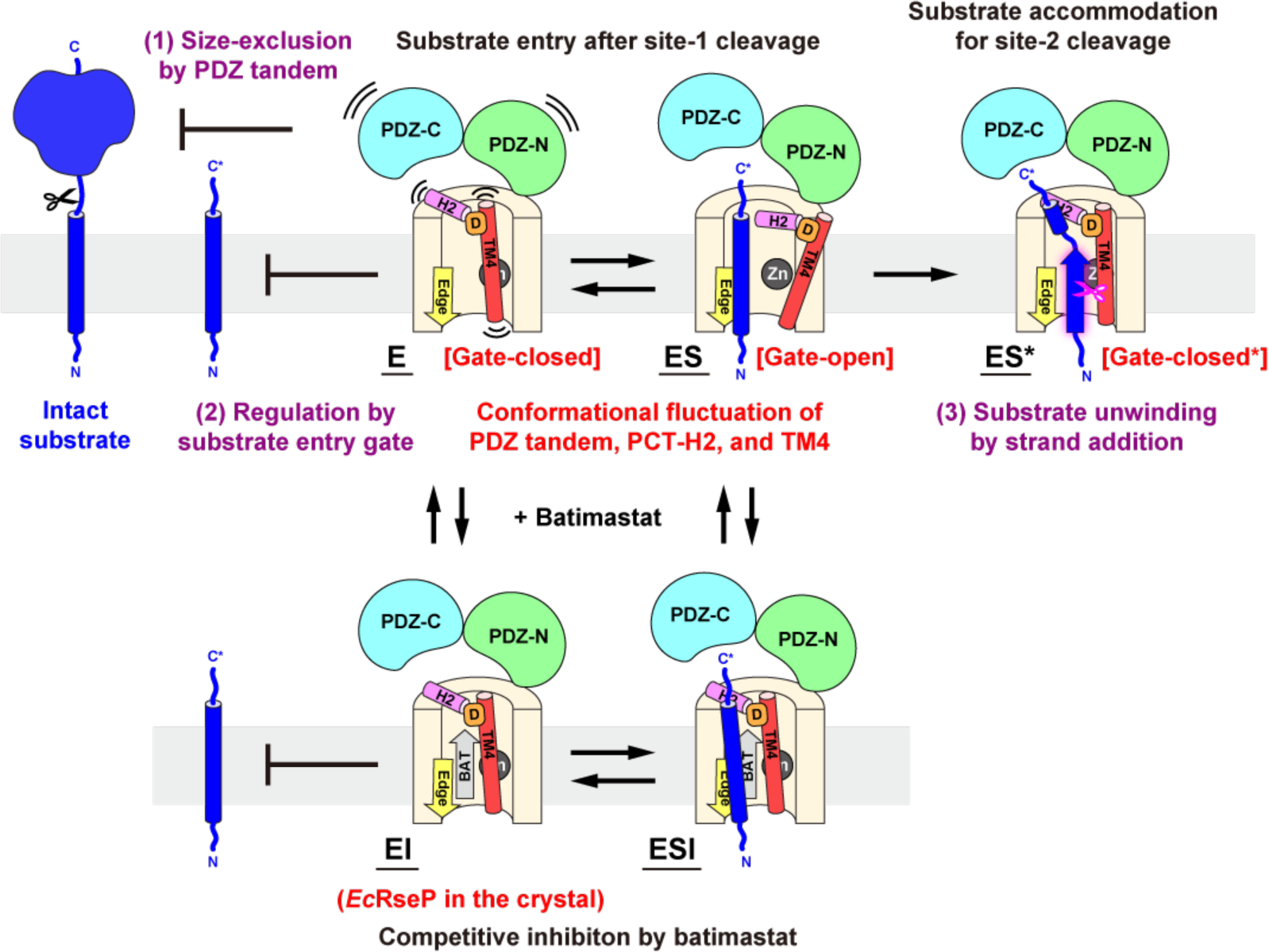
Proposed mechanism for substrate accommodation and cleavage in *Ec*RseP. The substrate accommodation by *Ec*RseP is thought to be regulated by three proesses. (1) Size-exclusion process: The PDZ tandem serves as a size-exclusion filter to restrict the entry of the intact substrates with a bulky extracytoplasmic domain. After the site-1 cleavage, the size-reduced substrate becomes accessible to the TM domain of *Ec*RseP. (2) Gating process: The PDZ tandem, PCT-H2, and TM4 serve as a gate to regulate the substrate entry into the active center. It is presumed that *Ec*RseP is in equilibrium between the gate-closed and gate-open conformations. In the gate-closed conformation (labeled the E state), the PDZ tandem is closed by interacting with the PCT region, and PCT-H2 is accommodated in the TM1-TM3 cleft above the active center. TM4 also covers the active center. Thus, the active center is inaccessible to the substrate TM segment. Although the gate-open conformation is not yet observed experimentally, the present study indicates that the PDZ tandem is separated from the PCT region upon substrate binding while the PCT-H2 moves away from the active center (labeled the ES state). In this gate-opening movement, TM4 is also thought to be displaced along with PCT-H2 while maintaining an electrostatic interaction via the conserved residue D446 (indicated in orange). Despite the conformational changes, the size-exclusion function of the PDZ tandem is assumed to be maintained as the PDZ-pocket is still inaccessible to substrates with the bulky extracytoplasmic domain. (3) Unwinding process: As a final step, the TM segment of the substrate is extended by the edge strand of the MREβ sheet to promote the cleavage (labeled the ES* state). We anticipate that the gate adopts a relatively closed conformation (labeled “Gate-closed*”) again to shield the substrate TM segment from the hydrophobic milieu and to form a hydrophilic compartment around the active site for efficient hydrolysis. Based on this substrate accommodation model, the crystal structure of *Ec*RseP with batimastat corresponds to the EI state. The substrate entry is suppressed not only by the binding of the inhibitor batimastat (gray arrow labeled “BAT”), but also by the gate closure. The crystal structure of batimastat-bound *Kk*RseP is thought to be non-physiological, but may partly reflect the ESI state, in which TM4 of the crystal packing neighbor mimics the substrate TM segment.

Previous structural and mutational analyses have proposed that the PDZ tandem serves as a size-exclusion filter to regulate the sequential cleavage of the substrates (*20*). It was also proposed that PCT-H1 is involved in the size-exclusion as an adaptor (*21*). Furthermore, most point mutations that deregulate sequential cleavage have been located in the PDZ-N domain (*29*). Based on the trypsin susceptibility, those deregulated mutations are presumed to cause large structural changes or unfolding of PDZ-N (*20*). The present crystallographic analysis of *Ec*RseP has shown that the PDZ-C domain protrudes in front of PCT-H2 to lie above the active center and thus appears to sterically hinder entry for substrates with a bulky periplasmic domain (Fig. 1d). Additionally, the PDZ-N domain makes direct contacts with the PCT-H1 in *Ec*RseP. Thus, the unfolding of the PDZ-N domain should destabilize the interaction between PDZ-N and PCT-H1, which may also perturb the geometry of size-exclusion filter to deregulate its function. The Cys-scanning mutagenesis in this study further demonstrated that the PCT-loop and -H2 also regulate the sequential cleavage (Fig. 4b). The EM analysis also suggested that the mutation to the PCT-loop region (L358C) causes deregulation through fluctuations in the orientation of the PDZ-C (Fig. S20). These observations indicate that the PDZ-N domain and the PCT region serve as a scaffold to place the PDZ-C domain in a position to perform the size-exclusion function.

Moreover, the present study indicates a domain rearrangement occurs during the substrate accommodation and cleavage. Based on the modification and cross-linking analyses, we infer that the PDZ tandem, PCT-H2, and TM4 in *Ec*RseP serve as a gate for substrate entry. We anticipate that *Ec*RseP on the cell membrane is in equilibrium between the “gate-open” and “gate-closed” conformations. The AMS modification analysis indicates that PCT-H2 is mobile on the *E. coli* cell membrane. After the site-1 cleavage in the extracytoplasmic region, the substrate passes through the size-exclusion filter to access the TM domain of *Ec*RseP. The substrate TM segment is thought to enter the active center when PCT-H2 and TM4 move away together, linked by the electrostatic interaction between the PCT-H2 backbone and conserved residue D446. Based on the intramolecular cross-linking analysis, the PDZ tandem is also thought to reorient relative to the PCT region during the substrate accommodation whereas the PDZ pocket is still inaccessible enough to maintain the size-exclusion function. The gating mechanisms are also presumed to exist not only in the S2P family, but also in the other intramembrane proteases such as bacterial rhomboid GlpG (*30, 31*) and γ-secretase (*32*), a member of the presenilin/SPP family. In addition, it was proposed that *E. coli* rhomboid GlpG possesses an “interrogation site” for substrates in addition to a “scission site” to discriminate the substrates from non-substrates (*33*). As the cleavage efficiency in *Ec*RseP differs depending on the substrate, the proposed gating mechanism may also contribute to the substrate interrogation.

After the gate opening for the substrate accommodation, the substrate cleavage segments should bind with the edge strand via strand addition. In addition, we anticipated that the gate adopts a relatively closed conformation again to shield the substrate TM segment from the hydrophobic milieu and to form a hydrophilic compartment around the active site for hydrolysis. Mutational analysis in this study suggests that the conserved N394 residue contributes to the stabilization of the substrate in a conformation efficient for cleavage. Our previous co-immunoprecipitation assay also showed that a cysteine mutation to N394 lowered the affinity for the TM segment of RseA (*34*), and the corresponding residue (N129) in the *Bacillus subtilis* homologue SpoIVFB has been reported to be also important for substrate cleavage (*35*). Because SpoIVFB is a Group III S2P, this backbone interaction may be common across diverse S2Ps. Furthermore, clamp-like interactions with unwound substrates appear to exist across the intramembrane proteases. The cryo-EM analysis of γ-secretase classified as an aspartic protease showed that substrate binding induces the formation of a β-sheet in the TM region of presenilin (*32, 36*) (Fig. S23a, b). The β-sheet binds with the substrate fragment via strand addition. In γ-secretase, the PAL motif is located on the opposite side of the β-sheet like a clamp for the substrate. For the serine protease Rhomboid family, crystal structures of *E. coli* GlpG have been determined in complex with a model substrate (*30*) (Fig. S23c, d). In those complexes, two loops connecting the TM helices sandwich the substrate peptide in an extended conformation, forming an integrated β-sheet. Despite the difference in the catalytic mechanism, unwinding the TM helix into a strand conformation and stabilizing the bound substrate with a clamp structure seem to be common features in the intramembrane proteolysis of helical TM spans. Structure determination of a substrate-bound and -unbound forms of RseP or the Group I S2Ps will deepen our understanding of the substrate cleavage mechanism conserved across the intramembrane proteases and aid the development of strategies for regulating proteolytic activity in the membrane to prevent off-target or promiscuous cleavage.

## Acknowledgements

We are grateful to the beamline staff of SPring-8 BL32XU (Hyogo, Japan) and Photon Factory (Tsukuba, Japan) for providing data collection facilities and support. We thank National BioResource Project (NBRP)—*E. coli* at National Institute of Genetics, Japan for the KEIO strain. We also thank Samuel Thompson for his critical reading and editing of the manuscript, prof. Junichi Takagi for valuable advice in the early stage of this study, profs. Hideo Takahashi, and Akinori Kidera for valuable discussions, prof. Tomohiro Nishizawa and associate prof. Kazuki Takeda for their advice on LCP crystallization, Sanae Tabata and Takeru Hirose for their contribution to antibody-assisted EM analysis, and Yutaro Yamagata and Saisei Honna for their contribution to *in vitro* substrate cleavage analysis.

## Funding

This research is partially supported by the Japan Society for the Promotion of Science (JSPS) KAKENHI under Grant Numbers JP19687004, JP22370039, JP26291016, and JP19H03170 (to T.N.), under JP21J15841 (to T.Y.), under JP19K06562 (to Y.H.), under JP18H02404 (to Y.A.), and under JP21K19236 (to S.A.), by the Ministry of Education, Culture, Sports, Science and Technology (MEXT) under JP19H05774 (to S.A.), by the Platform Project for Supporting in Drug Discovery and Life Science Research (Platform for Drug Discovery, Informatics, and Structural Life Science) from the Japan Agency for Medical Research and Development (AMED) under Grant Numbers JP16am0101020 (to T.N.), by the Platform Project for Supporting Drug Discovery and Life Science Research (Basis for Supporting Innovative Drug Discovery and Life Science Research (BINDS)) from AMED under Grant Number: JP21am0101078 (to Y.K.), by the Sumitomo Foundation, Basic Science Research Projects, by the Astellas Foundation for Research on Metabolic Disorders, by the grant for 2018 Research Development Fund of Yokohama City University, and by the Institute for Frontier Life and Medical Sciences, Kyoto University for INFRONT Office of Director’s Research Grants Program (2020) and (2021) (to Y.H.).

This work was performed in part under the Cooperative Research Program (Joint Usage/Research Center program) of the Institute for Frontier Life and Medical Sciences, Kyoto University, under the Collaborative Research Program of the Institute for Protein Research, Osaka University, CR-21-06, and under the Cooperative Research Project Program of Life Science Center for Survival Dynamics, Tsukuba Advanced Research Alliance (TARA Center), University of Tsukuba.

## Author contributions

T.N., Y.H., and Y.A. conceived the project. T.N. supervised the structural analysis. R.O. contributed to preparation of the full-length RseP. M.Tak. and O.N. identified the RseP ortholog suitable for structural analysis. K.M. and S.K. performed the crystallographic analysis of the partial fragment. Y.I. and K.T. performed the crystallographic analysis of the full-length RsePs. K.H. and H.M. collected and analyzed the diffraction data. R.A., M.H., K.I., and T.Ka. performed the EM analysis. M.K.K. and Y.K. contributed to preparation of the antibody fragment. M.Taj. and S.A. performed the MS analysis. Y.A. and Y.H. supervised the biochemical analysis. Biochemical analyses were performed by T.M., T.Ko., and Y.H. (*in vivo*) and by T.Y. (*in vitro*). All the authors contributed to paper preparation. T.N., Y.A, and Y.H. wrote the original draft of the manuscript, and T.N. compiled the paper.

## Data and materials availability

The atomic coordinates were deposited in the Protein Data Bank with the accession codes as follows: full-length *Ec*RseP (7W6X), full-length *Kk*RseP in the *P*1 crystal (7W6Y), full-length *Kk*RseP in the *P*21 crystal (7W6Z), *Kk*PDZ-C domain (7W70), and *Ec*PDZ-C domain complexed with the 12C7 Fab (7W71).

## Materials and Methods

### Construction of expression plasmids for structural analysis

A pUC118-based plasmid was constructed to produce *Ec*RseP for structural analysis. The resulting plasmid, designated as pNY1452, carried a gene encoding the wild-type full-length *Ec*RseP fused with a C-terminal tag containing the tobacco etch virus (TEV) consensus sequence, a His8 tag, a Myc epitope, and a PA tag (*37*): -GT-ENLYFQG-G-HHHHHHHH-I-EQKLISEEDL-GVAMPGAEDDVV. For negative-stain EM, an expression plasmid for the L358C mutant, designated as pNY1550, was constructed from pNY1452 by the inverse PCR method.

In parallel, we also searched for RseP orthologues suitable for structural analysis. Genes of 14 orthologues were amplified using genomic DNA obtained from the Japan Collection of Microrganisms (RIKEN, Microbe Division) and sub-cloned into NdeI and XhoI sites of a modified pET-21b vector. Using the constructed plasmids, the orthologues were produced with a C-terminal tag containing the TEV consensus sequence and a His8 tag: - LESSG-ENLYFQG-QFTS-HHHHHHHH. To examine the production level and dispersity, the orthologues produced in small-scale culture were subjected to detergent screening with fluorescence-detection size-exclusion chromatography (FSEC) after the C-terminal His8 tag was labeled using a peptide-based multivalent nitrilotriacetic acid (NTA) fluorescent probe, P3NTA. From the screen, an orthologue Kkor_1905 from *Kangiella koreensis* str. DSM 16069 (*Kk*RseP; UniProtKB: C7R5Z1), a gram-negative bacterium isolated from sea water of the tidal flat (*38*), was selected as a promising candidate for structural analysis. The expression plasmid carrying the *Kk*RseP gene was designated as pNY1432. The highest yield of monodisperse *Kk*RseP sample was obtained with an optimized preparation using β-dodecylmaltoside (DDM) supplemented with cholesteryl hemisuccinate (CHS).

The pGEX-2T (Cytiva) vector was used for the expression plasmids for soluble fragments. pNO1494, pNO2413, and pNY1480 contained genes encoding the *Ec*RseP PDZ tandem (EcPDZ tandem; P125-I309), *Ec*RseP PDZ-C (*Ec*PDZ-C; G219-I309), and *Kk*RseP PDZ-C (*Kk*PDZ-C; G221-Q307), respectively. In pNO2413 and pNY1480, a His8 tag and the TEV consensus sequence were incorporated between the N-terminal GST (glutathione S transferase) and the PDZ domain sequences to facilitate the tag removal with TEV protease. The strains and plasmids used in the structural analysis including the procedures described below are summarized in Supplementary Table 1 and 2.

### Production and purification of the *Ec*RseP PDZ tandem for antibody generation

For generating monoclonal antibodies against *Ec*RseP, the *Ec*PDZ tandem fragment was overproduced in *E. coli* BL21(DE3). *E. coli* cells transformed with pNO1494 were grown at 37°C to an OD600 of 0.7 in an LB medium supplemented with 50 μg/mL ampicillin, followed by induction of overexpression with 1 mM IPTG and incubation at 37°C for additional 4 hours. Cells were harvested by centrifugation and lysed by sonication in phosphate buffered saline (PBS): 137 mM NaCl, 2.7 mM KCl, 10 mM Na_2_HPO_4_, and 1.8 mM KH_2_PO_4_. The soluble fraction of the cell lysate was mixed with Glutathione-Sepharose 4B resin (Cytiva) and incubated at 4°C for 1 hour. After washing out the unbound fraction with PBS, the *Ec*PDZ tandem was cleaved off from the GST portion through on-column digestion with thrombin (Cytiva) at 20°C for 12 hours. The released *Ec*PDZ tandem was further purified using anion-exchange chromatography HiTrap Q HP (Cytiva). As a final step of purification, *Ec*PDZ tandem was applied to a Superdex 200 10/300 GL (Cytiva) column equilibrated with PBS.

### Generation of monoclonal antibody for *Ec*RseP

GANP transgenic mice (*39*) were immunized with the purified *Ec*PDZ tandem. Injections were repeated 4 times with intervals of two weeks. Spleen cells were obtained from the immune mouse showing the highest titer and were fused with the P3U1 myeloma cell line. The fused cells were selected in complete RPMI1640-10% fetal calf serum (FCS) with HAT supplement and subjected to limiting dilution culture, followed by culture in HT medium. The reactivity and titer of antibodies were examined by enzyme-linked immunosorbent assay (ELISA) using plates coated with the *Ec*PDZ tandem protein. An epitope domain for each clone was identified by ELISA using plates coated with PDZ-N or -C domain fragments. 12C7 was established as a monoclonal antibody recognizing the PDZ-C domain.

Milligram quantities of the 12C7 IgG was purified from the ascites through Protein A affinity chromatography. The purified IgG was digested at 37°C for 12 hours using Immobilized Papain (Thermo Scientific) with a buffer containing 12 mM Na_2_HPO_4_, 8 mM KH_2_PO_4_, 10 mM EDTA, and 100 mM Cys-HCl. After removing the Immobilized Papain resin, the digestion mixture was dialyzed against 20 mM Na-phosphate (pH 7.0) and applied to Protein G Sepharose resin (Cytiva). After washing out the unbound fraction with 20 mM Na-phosphate (pH7.0), the 12C7 Fab was eluted from the resin by adding 100 mM Gly-HCl (pH 2.4). The elution fractions were immediately neutralized by adding a quarter volume of 1 M Tris-HCl (pH9.0). The 12C7 Fab was further purified using a Superdex 200 10/300 GL SEC column (Cytiva) equilibrated with 10 mM Tris-HCl (pH 7.4), and 150 mM NaCl.

For primary structure determination, total RNA was prepared from the hybridoma cells, 12C7, using an RNeasy PLUS Mini Kit (Qiagen Inc., Hilden, Germany). The initial cDNA strand was synthesized using the SuperScript IV Reverse Transcriptase (Thermo Fisher Scientific, Inc., Waltham, MA) via a priming oligo-dT, according to the manufacturer’s instructions. PCR amplification was performed with oligonucleotide mixtures of the degenerate primer and the constant region primer for the heavy (γ) and light (κ) chains. PCR reactions were performed with HotStar *Taq* polymerase (Qiagen Inc.). The PCR products were purified, sub-cloned into the pCR4-TOPO vector (Thermo Fisher Scientific, Inc.), and subjected to Sanger nucleotide sequencing.

### Production, purification, and crystallographic analysis of the PDZ-C domains

The *Ec*PDZ-C and *Kk*PDZ-C domains were also produced as an N-terminal GST-fusion protein in *E. coli* BL21(DE3) cells. The purification procedure was essentially the same as that for the *Ec*PDZ tandem described above, except that TEV protease was used to cleave the N-terminal GST portion from the target PDZ-C domains. For the *Ec*PDZ-C domain, the purified protein was mixed with the 12C7 Fab and applied to a Superdex 200 10/300 GL SEC column (Cytiva) equilibrated with 10 mM Tris-Cl (pH 7.4), and 150 mM NaCl to fractionate the complex.

Initial crystallization conditions were searched using the Index^TM^ (Hampton Research) screening kit. 0.2 μL each of protein solution and crystallization buffer were dispensed into 96-well plates using a Crystal Gryphon (Art Robbins Instruments) and equilibrated against 60 μL of crystallization buffer in the reservoir by the sitting-drop vapor-diffusion method. The *Ec*PDZ-C domain complexed with the 12C7 Fab was crystallized in a solution containing 25% (wt./vol.) PEG 10,000, 0.2 M ammonium sulfate, and 0.1 M Bis-Tris-HCl (pH 5.5). The *Kk*PDZ-C domain was crystallized in a solution containing 1.0 M ammonium sulfate and 0.1 M HEPES-Na (pH7.5).

All crystals were quickly soaked in the cryoprotectant, which was prepared by mixing the crystallization buffer and ethylene glycol in a ratio of 4:1 (vol./vol.), and frozen in liquid nitrogen. X-ray diffraction data were collected using PILATUS3 S 6M (Dectris) at Photon Factory (PF) BL-17A (Tsukuba, Japan). Data were processed and scaled with XDS (*40*) and aimless (*41*). Diffraction intensities were converted to structure factors using the CCP4 suite where 5% of the unique reflections were randomly selected as a test set for the calculation of free *R*-factor (Winn *et al.*, 2011). Data collection statistics are summarized in Table S4.

For both crystals, initial phases were determined by the molecular replacement method using Molrep (*42*) in CCP4. For the *Ec*PDZ-C domain complexed with 12C7 Fab, the Fv and constant regions were assigned separately using the crystal structure of the mouse IgG Fab that was raised against the PDZ tandem of an RseP homologue from *Aquifex aeolicus* (*20*) (PDB code: 3WKM). Subsequently, the *Ec*PDZ-C domain was searched using the atomic coordinates of the individual *Ec*PDZ-C domain (*29*) (PDB code: 2ZPM). As a result, two complexes were assigned in the asymmetric unit. For the *Kk*PDZ-C domain, the structure of *Ec*PDZ-C domain (PDB code: 2ZPM) were used as a search model, and two monomers were assigned in the asymmetric unit. The initial models were manually modified and fit into the electron density map using the program COOT (*43*). The updated models were refined with phenix.refine (*44*) iteratively while monitoring the stereochemistry with MolProbity (*45*).

### Production and purification of full-length *Ec*RseP and *Kk*RseP

*Ec*RseP was overproduced in *E. coli* C43(DE3) (Lucigen). *E. coli* C43(DE3) cells transformed with the expression plasmid pNY1452 were grown at 30°C to an OD600 of 0.7 - 0.8 in LB medium (10 g of bactotryptone, 5 g of yeast extract and 10 g of NaCl per liter) supplemented with 50 μg/mL ampicillin, followed by induction of overexpression with 0.1 mM IPTG and incubation at 30°C for additional 4 hours. Cells were harvested by centrifugation and lysed by sonication in 10 mM Tris-HCl (pH 7.4) with 150 mM NaCl. The cell lysates were centrifuged at 40,000 ×g for 45 min at 4°C. Subsequently, the supernatant was separated by ultracentrifugation at 200,000 ×g for 90 min at 4°C. The membrane fraction collected as a precipitant was suspended in 10 mM Tris-HCl (pH 7.4) with 150 mM NaCl and was ultracentrifuged again under the same conditions. Finally, the precipitated membrane fraction was suspended in 10 mM Tris-Cl (pH 7.4) with 150 mM NaCl and the total protein was quantified using the bicinchoninic acid (BCA) assay. The resuspended membrane fraction was diluted to adjust the protein concentration to 10 mg/mL using bovine serum albumin as a standard.

*Ec*RseP was solubilized by adding equal volume of a solubilization buffer containing 40 mM Tris-HCl (pH 8.5), 150 mM NaCl, and 2% sucrose monododecanoate (SM) to the suspension of the membrane fraction. After incubation at 4°C for 1 hour, the mixture was ultra-centrifuged at 210,000 ×g for 90 min. at 4°C. The supernatant was applied to NZ-1 antibody conjugated Sepharose resin (anti PA-tag) and the unbound fraction was washed out with a buffer containing 10 mM Tris-HCl (pH 8.5), 150 mM NaCl, and 0.05% SM. *Ec*RseP was eluted from the resin with a buffer containing 10 mM Tris-HCl (pH 8.5), 150 mM NaCl, 0.05% SM, and 0.1 mg/mL PA14 peptide (EGGVAMPGAEDDVV). The C-terminal tag was cleaved off by adding TEV protease to the elution fraction, followed by incubation at 20°C overnight. The reaction mixture was then applied to a Superdex 200 10/300 GL size-exclusion chromatography (SEC) column (Cytiva). The peak fraction containing putative monomeric *Ec*RseP was collected and again applied to the same SEC column to remove oligomeric *Ec*RseP and aggregated TEV protease. After the second round of SEC, monodisperse *Ec*RseP sample was obtained with high purity (∼95%).

To produce *Kk*RseP, the plasmid pNY1432 was transformed into *E. coli* C43(DE3) harboring pRARE2. For the native *Kk*RseP, *E. coli* cells were grown at 30°C to an OD600 of 0.6 - 0.7 in an LB medium supplemented with 50 μg/mL ampicillin and 34 μg/mL chloramphenicol, followed by induction of overexpression with 0.1 mM IPTG and incubation at 16°C for additional 18 hours. To produce the selenomethionine (SeMet)-substituted *Kk*RseP, *E. coli* cells were first cultured in an LB medium supplemented with 50 μg/mL ampicillin and 34 μg/mL chloramphenicol grown at 30°C overnight. The cells were further inoculated into M9 medium supplemented with 50 μg/mL ampicillin and 34 μg/mL chloramphenicol, and grown at 30°C. At an OD600 of 0.3, a mixture of amino acids was added to the medium where the final concentrations were as follows: 100 mg/mL lysine, 100 mg/mL phenylalanine, 100 mg/mL threonine, 50 mg/mL valine, 50 mg/mL leucine, 50 mg/mL isoleucine, 60 mg/mL SeMet. At an OD600 of 0.7 - 0.8, 0.1 mM IPTG was added to induce the overproduction, followed by incubation at 16°C overnight. Harvested cells were lysed by sonication in 10 mM Tris-HCl (pH 7.4) with 150 mM NaCl, followed by centrifugation 20,000 ×g for 30 min. at 4°C. Subsequently, the supernatant was further separated by ultra-centrifuge at 200,000 ×g for 60 min at 4°C. The collected membrane fraction was finally suspended in 20 mM Tris-HCl (pH 8.0) 300 mM NaCl, 15% glycerol, and 0.1 mM PMSF, and the total protein concentration was adjusted to 10 mg/mL.

*Kk*RseP was solubilized by adding equal volume of a solubilization buffer containing 40 mM Tri-HCl (pH 8.0), 300 mM NaCl, 20 mM imidazole, 10% glycerol, 2% β-DDM, and 0.4% CHS to the suspension of the membrane fraction. After incubation at 4°C for 1 hour, the mixture was ultra-centrifuged at 210,000 ×g for 30 min. at 4°C. The supernatant was applied to Ni-NTA agarose resin and the unbound fraction was washed out with a buffer containing 20 mM Tris-HCl (pH 8.0), 300 mM NaCl, 10% glycerol, 0.03% β-DDM, 0.006% CHS and 50 mM imidazole. *Ec*RseP was eluted with a buffer containing 20 mM Tris-HCl (pH 8.0), 300 mM NaCl, 10% glycerol, 0.03% β-DDM, 0.006% CHS and 300 mM imidazole. The C-terminal tag was cleaved off by adding TEV protease to the elution fraction, followed by incubation at 20°C overnight. The reaction mixture was dialyzed against a buffer containing 20 mM Tris-HCl (pH 8.0), 300 mM NaCl, 10% glycerol, 0.03% β-DDM, 0.006% CHS and 10 mM imidazole, and was applied to the Ni-NTA agarose resin and the the tag-cleaved *Kk*RseP was collected in the unbound fraction. *Kk*RseP was further purified with a Superdex 200 10/300 GL SEC column (Cytiva).

### Crystallographic analysis of full-length RsePs

Batimastat (Toronto Research Chemicals) was added to the SEC elution fractions of *Ec*RseP and *Kk*RseP, respectively, at a final concentration of 300 μM. Subsequently, *Ec*RseP and *Kk*RseP were concentrated up to 7-12 mg/mL by ultrafiltration, and incorporated into lipidic cubic phase (LCP) by mixing the protein solution and monoolein (NU-CHECK-PREP, Inc.) with a volume ratio of 5:8 using two syringes attached with a coupler. Crystallization conditions were searched and optimized by microbatch crystallization method using Laminex™ sandwich plates and MemMeso™ screening Kit (Molecular Dimensions). 50 or 100 nL aliquots of the protein-monoolein mixture were dispensed onto 96-well glass plates and overlaid with 800 nL of crystallization solution using a mosquito LCP (TTP LabTech) or Crystal Gryphon LCP (Art Robbins Instruments).

The crystals of *Ec*RseP used for data collection were obtained in a crystallization solution containing 30% (vol./vol.) polyethylene glycol (PEG) 500 dimethyl ether, 100 mM NaCl, 100 mM MgCl2, and 100 mM HEPES-Na (pH7.0), while those of *Kk*RseP were generated from crystallization solutions containing 28-30% (vol./vol.) PEG 500 monomethyl ether, 100 mM NaCl, 100 mM CaCl2, and 100 mM HEPES-Na (pH7.0). Crystals were harvested using MicroMounts™ (MiTeGen) or LithoLoops (Molecular Dimensions), and flash-frozen in liquid nitrogen.

Diffraction data from the crystals obtained from the LCP crystallization were collected at SPring-8 beamline BL32XU (*46*) using an EIGER X9M detector (Dectris). Microfocused X-rays with a beam size of 15 µm × 10 µm at wavelengths of 1.0000 Å (*Ec*RseP native data), 1.2800 Å (Zn-anomalous diffraction data), and 0.9700 Å (Se-anomalous diffraction data), respectively, were used for both raster scan and data collection. A dataset with a total oscillation range of 10° and 0.1° oscillations per frame was collected from each crystal under an absorbed dose of 10 MGy. The partial datasets collected with the automated data-collection system ZOO (*47*) were merged, integrated, and scaled using the KAMO system (*48*), which integrates BLEND (*49*), XDS, and XSCALE (*40, 50*).

Initial phases of the SeMet-substituted *Kk*RseP crystal were determined by a combination of molecular replacement and single-wavelength anomalous diffraction methods (MR-SAD). After molecular replacement using the structure of *Kk*PDZ-C domain as a search model, the Se-SAD phasing was performed using phenix.autosol (*51*). The structure of *Ec*RseP was solved by MR using the partial model of *Kk*RseP in addition to the individual *Ec*PDZ-N (PDB code: 2ZPL) and *Ec*PDZ-C (PDB code: 2ZPM) models (*29*). The PCT domain and TM4, where the conformations were significantly different from those of *Kk*RseP, were modelled manually. Model modification, structure refinement, and validation were performed according to the same procedures as described above for the PDZ-C domains. Statistics for data collection and refinement are summarized in Table S5 and S6, respectively.

Structural superposition and RMSD calculation were performed by a pair-wise alignment protocol using LSQKAB (*52*). Figures for protein structures were prepared with PyMOL (The PyMOL Molecular Graphics System, Version 2.3 Schrödinger, LLC.).

### Negative-stain EM

For the negative-stain EM analysis, the wild-type *Ec*RseP and the L358C mutant were both produced according to the same method as that for the crystallization sample, but purified with a modified method. After the wash step for immunoaffinity chromatography using NZ-1 Sepharose, the detergent used for solubilization was exchanged by washing the resin to a buffer containing 10 mM Tris-HCl (pH 8.5), 150 mM NaCl, and 0.005% glycol-diosgenin (GDN). Subsequently, the GDN-solubilized *Ec*RseP was eluted using a buffer containing 10 mM Tris-HCl (pH 8.5), 150 mM NaCl, 0.005% GDN, and 0.1 mg/mL PA14 peptide. Tag cleavage by TEV protease was performed similarly to that in the purification of the SM-solubilized *Ec*RseP. SEC was repeatedly performed using a Superdex 200 10/300 GL column until a monodisperse protein with high purity was obtained. The purified *Ec*RseP was mixed with the 12C7 Fab and again applied to the SEC column using the same buffer to fractionate the complex.

For negative-stain EM, all purified samples were diluted to 5.0 μg/mL. 5 μL of protein was applied to glow-discharged, 600 mesh, carbon-coated grids. The grids were negatively stained with 2% uranyl acetate, blotted with filter paper, and air dried. A Talos Arctica EM (Thermo) was operated at 200 kV to acquire micrographs using a Falcon3EC camera (Thermo) in linear mode. Micrographs were recorded at 1.588Å/pix and 1.4Å/pix for the wild-type and L358C mutant, respectively, with defocus of -0.5 to -2.0 μm. A movie of 20 frames was taken for each micrograph, and motion correction was performed using MotionCor2 (*53*). The contrast transfer function was estimated by Gctf (*54*) or CTFFIND-4.1 (*55*). All subsequent processing was carried out using RELION3.1 (*56*). Particles were selected using the Relion Autopicker and three rounds of 2D classification were performed to select particles for 3D reconstruction. The number of the particles used for each reconstruction of the Fab-complexed *Ec*RseP wild-type and L358C mutant are indicated in Fig. S20, respectively. Fitting of the atomic coordinates of Fab-PDZ complex into the 3D reconstruction model and figure preparation were performed with Chimera (*57*).

### Mass spectrometry

*Ec*RseP produced from pNY1452 (MW: 53703.3) and purified as described above was used for MS measurements. Purified *Ec*RseP in a buffer containing 10 mM Tris-Cl (pH 8.5), 150 mM NaCl, and 0.032 % (w/v) SM was applied to a centrifugal device (Micro Bio-Spin 6, Bio-Rad, Hercules) to exchange the buffer to a MS buffer containing 200 mM ammonium acetate (pH 7.4), and 0.5 % (w/v) n-octyltetraoxyethylene (C8E4). Prior to buffer exchange, the device was washed with 200 mM ammonium acetate containing 1.0 mM EDTA-Na to remove residual zinc ions completely, and then equilibrated with the MS buffer. Mass spectral data were acquired using a SYNAPT G2 HDMS instrument (Waters, Milford) equipped with a nano ESI source. The samples were deposited in platinum-coated borosilicate capillaries with a 5 µm i.d. purchased from HUMANIX (Hiroshima, Japan).

### Media for biochemical analysis

L medium (10 g/L Bacto Tryptone, 5 g/L yeast extract and 5 g/L NaCl; pH adjusted to 7.2 by using NaOH) and M9 medium (without CaCl2; (*58*)) supplemented with 2 μg/mL thiamine and 0.4% glucose were used to culture *E. coli* cells. Ampicillin (50 μg/mL), chloramphenicol (20 μg/mL), and/or spectinomycin (50 μg/mL) were added for selecting transformants and for growing plasmid-harboring cells.

### Construction of strains and plasmids for biochemical analysis

*E. coli* strain YH2902 was constructed as follows. The Δ*rseA*::*kan* region from JW2556 (*59*) was introduced into AD16 by P1 transduction, and then the *kan* cassette of the resulting strain was deleted using pCP20, as described previously (*60*), to obtain AD2543. Next, the Δ*acrA*::*kan* region from JW0452 (*59*) was P1 transduced into AD2543, and the *kan* cassette was similarly deleted to obtain YH2898. Finally, the Δ*rseP*::*kan* from KK211 was P1 transduced into YH2898 to obtain YH2902. AD2544 [formerly misdescribed as AD2328 in (*24*)], the parental strain of KA306, was constructed by P1 transducing Δ*rseP*::*kan* into AD2543.

Plasmids encoding a mutant form of *Ec*RseP or *Kk*RseP were constructed by standard site-directed mutagenesis using appropriate combinations of mutagenic primers and a template plasmid as described below. pYH131 [*EcrseP-his8* with the nSD (native Shine Dalgarno) sequence, encoding an *Ec*RseP derivative having a C-terminal His8 tag (EFIEGR-HHHHHHHH))] was constructed by cloning a PCR product amplified from plasmid pKK49 using primers P1/P2 into the KpnI site of pUC118. pYH820 (*EcrseP-his8* with an “improved” SD (iSD) sequence (AAGGAGGA) for stable protein expression) was constructed as follows; first, the PCR product amplified from pYH131 using primers P3/P4 was digested with EcoRI and ligated with the “vector” fragment of EcoRI-digested pYH131 to obtain pYH819. Then, the SacI/HindIII fragment of pYH819 was cloned into pTWV228, yielding pYH820. To construct pYH825 (*EcrseP* with iSD, encoding EcRseP with no C-terminal tag) and pYH826 [*EcrseP(E23Q)* with iSD], the PCR product amplified from pKK47 or pYK2 using primers P3/P4 was digested with EcoRI/HindIII and inserted into pTWV228. To construct pYH835 [*EcrseP(Cys-less)* with iSD], the PCR product amplified from pTM101 using primers P3/P5 was digested with EcoRI/HindIII and inserted into pTWV228. To construct pYH829 (*KkRseP-TEV-His8* with iSD), the PCR product amplified from n1432 using primers P6/P7 was digested with EcoRI/BamHI and inserted into pTWV228. To construct pYH833 (*KkRseP* with iSD), the PCR product amplified from pYH829 using primers P1/P8 was digested with EcoRI/HindIII and inserted into pTWV228. pYH838 (*KkRseP* with iSD, encoding a *Kk*RseP derivative having an internal PA14 tag: Lys54-EGGVAMPGAEDDVV-His55) was constructed by using the In-Fusion HD Cloning Kit (Takara Bio) as follows. A DNA fragment for the PA14 tag was prepared by annealing oligonucleotides P9/P10, and then connected with the linearized vector backbone amplified from pYH833 using primers P11/P12. pYH817 (*ha-mbp-rseA(KkTM)148*) was constructed by replacing the SalI/PstI fragment for the TM sequence of HA-MBP-RseA(LY1)148 on pYH124 (*19*) by a fragment for the TM sequence (P^100^FKQ<u>LAGYAIAASVALVVLFNV</u>G^122^, predicted TM region was underlined) of *Kk*RseA (UniProtKB: C7RA49, *Kangiella koreensis* str. DSM 16069: Kkor_0748) that had been prepared by annealing oligonucleotides P13/P14. pTM949 (pBAD33-*ha-mbp-rseA(LY1)148)* was constructed by cloning the SacI/HindIII fragment of pYH20 into pBAD33. The strains, plasmids and oligonucleotides used in this study are summarized in Table S1-S3.

### *In vivo* proteolytic activity assay of *Ec*RseP and *Kk*RseP and their derivatives

The *in vivo* proteolytic activities of *Ec*RseP and *Kk*RseP were analyzed essentially as described previously (*24, 61*). Precultured cells were inoculated into M9 medium supplemented with 20 µg/mL of each of the 20 amino acids, 2 µg/mL thiamine, 0.4% glucose, 1 mM IPTG and 1 mM cAMP and grown at 30°C for 3 h. The proteins were precipitated by trichloroacetic acid (TCA) treatment, washed with acetone, suspended in a SDS sample buffer with 2-mercaptoethanol (2ME) and analyzed by Laemmli SDS-PAGE and immunoblotting using an Immobilon membrane filter (MilliporeSigma) and appropriate antibodies. Proteins were visualized by Lumino image analyzer LAS-4000 mini (Cytiva) using ECL or ECL Prime Western Blotting Detection Reagents (Cytiva). Rabbit polyclonal anti-HA (HA-probe (Y-11), Santa Cruz Biotechnology), anti-MBP (*62*), anti-*Ec*RseP (*19*) and anti-SecB (*21*) antibodies, and rat monoclonal anti-PA antibodies (*63*) were used for immunoblotting. Also, for detection of His-tagged proteins, anti-His antibodies from the Penta-His HRP Conjugate Kit (Qiagen) were used. In the short induction method, precultured cells were inoculated into L medium with 0.4% glucose and grown at 30°C for 2.5 h. Collected cells were suspended in 100 μL of L medium and pre-incubated at 30°C for 10 min in an Eppendorf Thermomixer comfort (600 rpm). After addition of 1 mM IPTG, the cells were incubated at 30°C for 15 min with shaking. Finally, the proteins were precipitated by TCA treatment described above. Cleavage efficiencies of the substrates were calculated according to the following equation: Cleavage efficiency (%) = 100 × (cleaved)/[(cleaved) + (full length)], where (cleaved) and (full length) are intensities of the respective bands.

### In vitro substrate cleavage assay with purified EcRseP or KkRseP

Model substrate HA-RseA148 was synthesized using the PURE*frex®* 1.0 reconstituted cell-free protein synthesis kit (Gene Frontier Co., Japan, (*64, 65*)), using a template DNA (PCR amplified from pYH18) and standard reagents including 0.02% *n*-dodecyl-*β*-D-maltoside (DDM) and ^35^S-labeled methionine (^35^S-Met). *Ec*RseP or *Kk*RseP were purified according to the same procedures as those for the samples for crystallization. Synthesized ^35^S-labeled substrate was mixed with purified *Ec*RseP [final conc., 2.5 ng/μL (50 nM)], *Kk*RseP [final conc., 100 ng/μL (2.0 μM)] or each of the respective protein buffers (Mock) and incubated at 37°C in buffer containing 50 mM Tris-HCl(pH8.1), 2.5% glycerol, 5 μM zinc-acetate, 0.05% DDM, 100 mM NaCl, 10 mM DTT, and 5 mM of zinc chelator 1,10-phenanthroline (PT+) or 5% DMSO (PT-) with shaking in an Eppendorf Thermomixer comfort (500 rpm) for the indicated periods. Samples were then mixed with an equal volume of 2x SDS sample buffer plus 2ME and boiled for 5 min. The proteins were separated by SDS-PAGE using a 15% Bis-Tris gel and MES SDS running buffer (*66*) and visualized using a phosphor imager BAS5000 (Cytiva).

### *In vivo* batimastat sensitivity assay

Batimastat inhibition of *in vivo* proteolytic activity of RseP derivatives (batimastat sensitivity) was evaluated as described previously (*27*) with several modifications. YH2902 (Δ*acrA* background) cells harboring two plasmids encoding an *Ec*RseP derivative (pYH825-based) and a model substrate HA-MBP-RseA(LY1)148 (pYH124) were inoculated into L medium with 0.4% glucose and grown at 30°C for 2.5 h. Collected cells were suspended in L medium and divided into three portions. Each portion received 12.5, 3.125, or 0 μM (final conc.) batimastat (the stock solution of batimastat was dissolved in DMSO) and pre-incubated at 30°C for 10 min in an Eppendorf Thermomixer comfort (600 rpm). Samples were mixed with IPTG (final conc., 1 mM) to induce protein expression and incubated for 1 h with shaking. Proteins were TCA-precipitated and analyzed by SDS-PAGE and immunoblotting.

### Substituted Cysteine Accessibility Analysis of single-cysteine *Ec*RseP or *Kk*RseP mutants

A methoxypolyethylene glycol 5,000 maleimide (mal-PEG) accessibility assay was performed with a single Cys derivative of *Ec*RseP or *Kk*RseP using essentially the same method as described previously (*20, 61*). KK374 cells (*62*) carrying a plasmid encoding a tag-less *Ec*RseP Cys mutant or a *Kk*RseP Cys mutant with an internal PA14-tag were cultured in L medium containing 0.4% glucose, 1 mM IPTG and 1 mM cAMP at 30°C for 2.5 h and converted to spheroplasts by lysozyme/EDTA treatment as described previously (*29*). Spheroplast samples were treated with water or 2% Triton X-100 at 0°C for 3 min in the presence of 5 mM MgCl2, 1 mM phenylmethylsulfonyl fluoride (PMSF), and 1 mM Tris(2-carboxyethyl)phosphine (TCEP, MilliporeSigma). Samples were mixed with equal volume of 2x reaction buffer (60 mM Tris-HCl (pH 8.1) and 2 mM mal-PEG (MilliporeSigma)) and incubated at 4°C for the indicated periods. The Cys-modification reaction was stopped by adding 10% 2ME. The proteins were TCA-precipitated and analyzed by SDS-PAGE and immunoblotting.

Single Cys *Ec*RseP mutants were modified in two steps with 4-acetamide-4’-maleimidylstilbene-2,2’-disulfonic acid (AMS) and mal-PEG using essentially the same method as described previously (*21, 61*). Spheroplasts samples of KK374 cells expressing a single Cys derivative of RseP-His_6_-Myc (RseP-HM) were prepared as above, then treated with water or 1% Triton X-100 at 0°C for 30 min in the presence of 10 mM MgCl2, 1 mM PMSF and 1 mM TCEP. After pre-warming at 24°C for 5 min, samples were treated with 1 mM AMS (Thermo Fisher Scientific) in the presence or absence of 1% Triton X-100 at 24°C for 5 min. Following quenching of AMS by incubation with 62.5 mM DTT at 24°C for 18 min, the proteins were precipitated and washed with 5% TCA. Samples were solubilized in 100 mM Tris-HCl (pH 8.1) containing 1% SDS and 1 mM TCEP, and treated with 5 mM mal-PEG at 37°C for 1 h with vigorous shaking to modify free thiols. AMS/mal-PEG modified proteins were analyzed by SDS-PAGE and immunoblotting. The proportion of a RseP-HM Cys mutants modified with AMS was calculated according to the following equation: AMS modification (%) = 100 × (*a* − *b*)/*a*, in which *a* is the ratio of the mal-PEG-modified forms to total RseP-HM in the control sample that are prepared without AMS treatment, and *b* is the ratio of the mal-PEG-modified forms to total RseP-HM in the AMS-treated sample. For immunoblotting analysis of mal-PEG-labeled proteins, an Immobilon-P^SQ^ membrane filter (MilliporeSigma) and Transfer Buffer containing 24 mM Tris-base, 192 mM glycine, 10% methanol and 0.05% SDS were used.

### Complementation assay

*E. coli* KK31 [Δ*rseP*/pKK6 (PBAD-*rseP*)] cells harboring a plasmid encoding *Ec*RseP or its derivatives under the *lac* promoter were grown at 30°C in L medium containing 0.02% L-arabinose for 2 h. The cultures were serially diluted with saline and 3 μL of the diluted cultures were spotted on L agar plates containing 1 mM IPTG or 0.02% L-arabinose. The plates were incubated at 30°C for the indicated period.

### *β*-Galactosidase activity assay

The σ^E^ activity was assayed by monitoring *β*-Galactosidase (LacZ) activity expressed from a chromosomal σ^E^-dependent *lacZ* reporter gene (rpoHP3-*lacZ*). Cells were grown at 30°C for 5 h in L medium with 0.1 mM IPTG and 1 mM cAMP. The LacZ activity of growing cells was measured using essentially the same method as previously described (*21*).

### Trypsin susceptibility assay

The trypsin susceptibility assay for *Ec*RseP was performed using essentially the same method as previously described (*21*). Cells carrying a plasmid encoding a derivative of RseP-HM were grown at 30°C for 3 h in M9-based medium supplemented with 20 μg/mL each of the 20 amino acids, 1 mM IPTG, and 1 mM cAMP. Spheroplasts were prepared and treated with 2.5 μg/mL Trypsin on ice for the indicated time periods. Proteins were TCA-precipitated, suspended in a SDS sample buffer with 2ME, 1 mM PMSF, and 1 mM PefablocSC (MilliporeSigma), and then analyzed by SDS-PAGE and immunoblotting.

### Site-directed *in vivo* photo crosslinking

Site-directed *in vivo* photo crosslinking was carried out using essentially the same method as previously described (*21*). Cells harboring pEVOL-pBpF and a plasmid encoding an RseP(E23Q)-HM derivative were grown at 30°C for 4 h in M9 medium containing 0.5 mM *p*-benzoyl-L-phenylalanine (*p*BPA; Bachem AG). After adding 100 μg/mL spectinomycin to stop further protein synthesis, a portion of the culture was withdrawn and UV-irradiated for 10 min at 4°C using a B-100 AP UV lamp (365 nm; UVP, LLC.). Proteins were TCA-precipitated and dissolved in SDS sample buffer with 2ME.

### Disulfide crosslinking-mediated domain immobilization experiment

The *in vivo* proteolytic activity of *Ec*RseP after domain tethering by disulfide crosslinking was examined as follows. AD2544 cells harboring two independently-inducible plasmids, each encoding an RseP derivative with a pair of Cys mutations under the Plac promoter (pYH835-based) or the model substrate HA-MBP-RseA(LY1)148 (pTM949) under the PBAD promoter, were used. Precultured cells were inoculated into L medium with 0.4% glucose, grown at 30°C for 1.5 h, and divided into two portions. Collected cells were washed with and resuspended in 150 μL of L medium with 5 mM IPTG and incubated at 30°C for 30 min in an Eppendorf Thermomixer comfort (600 rpm) to induce expression of the RseP derivatives. After incubation, each of the two samples was washed with IPTG-free L medium, resuspended in L medium with 5 mM diamide (for oxidation) or 10 mM DTT (for reduction), and further incubated at 30°C for 30 min. The cells were washed again with L medium and resuspended in L medium containing 0.02% L-arabinose with (for the +DTT sample) or without (for the +diamide sample) 1 mM TCEP, and incubated at 30°C for 30 min to induce substrate expression. Finally, proteins were precipitated with TCA, resuspended in an SDS sample buffer without 2ME, and analyzed by SDS-PAGE and immunoblotting.

To evaluate the efficiency of intramolecular disulfide crosslinking for double-Cys RseP, all TCA-precipitated post-reaction samples were dissolved in 100 mM of Tris-HCl (pH 8.1) with 1% SDS and vigorously shaken for 30 min at room temperature. Then, the samples were divided into two portions, treated with 0 or 1 mM mal-PEG at 37°C for 1 h with vigorous shaking to modify free thiols, mixed with 2x SDS sample buffer containing 2ME, and finally analyzed by SDS-PAGE and immunoblotting.

### Multiple alignment analysis of bacterial RseP homologues

Selection and amino acid sequence alignment of RseP homologues were performed as follows. BLAST search (blastp) (https://blast.ncbi.nlm.nih.gov/Blast.cgi) was performed, using the amino acid sequence of *Kk*RseP (*Kangiella koreensis* str. DSM 16069: Kkor_1905 (UniProtKB: C7R5Z1)) as the query, against the Non-redundant UniProtKB/SwissProt sequence database. From the 66 S2P homologs found by this search, duplicates, non-bacterial homologs, and homologs with obviously different molecular sizes were manually excluded, leaving 39 bacterial homologues. Multiple sequence alignment was performed with these 39 species by the Clustal W ver.2.1 program (http://www.clustal.org/clustal2/) using genetic information processing software Genetyx (GENETYX Corporation, Japan).

