## Supplementary Figures & Tables for "Mechanistic insights into intramembrane proteolysis by *E. coli* site-2 protease homolog RseP"

This file includes:

Figs. S1 to S23

Tables S1 to S6

### Supplementary Figures

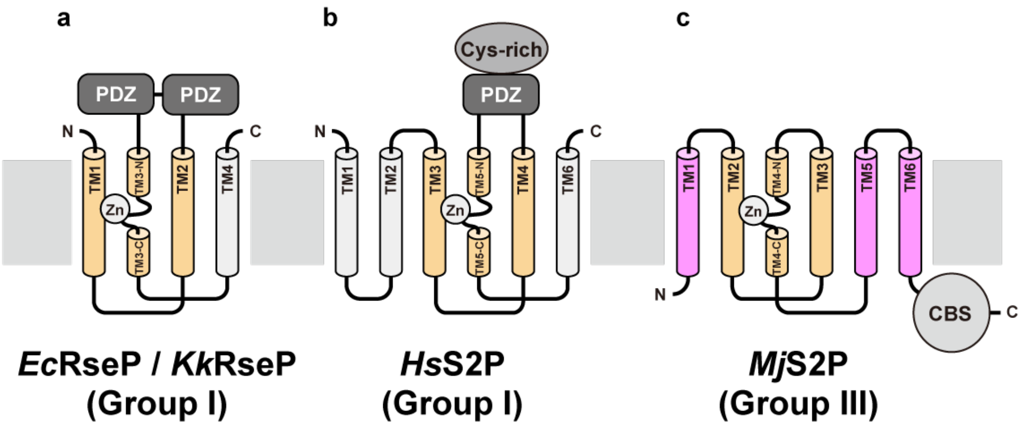

**Figure S1. Domain organization of the S2P family members**

The topology of TM domain and the location of putative soluble globular domains are schematically shown for **(a)** *E. coli* RseP (*Ec*RseP) and the *Kangiella koreensis* RseP orthologue (*Kk*RseP) of the Group I, **(b)** human S2P (*Hs*S2P) of the Group I, and **(c)** *Mj*S2P of the Group III, respectively. In each protein, the three TM helices colored orange constitute a core region conserved across the entire S2P family and contain the zinc-coordinating and catalytic residues. The remaining helices colored gray or light magenta are less or not conserved among the subfamilies. The members of the Group I subfamily possess a different number of PDZ domains in the extracytoplasmic region. In addition, a Cys-rich region is predicted to be inserted into the PDZ domain in *Hs*S2P. Members of the Group III subfamily (e.g. *Mj*S2P) possess a cystathionine-β-synthase (CBS) domain in the cytoplasmic C-terminal region, but they contain no PDZ domain. In *Mj*S2P, TM1, 5, and 6 (light magenta) were proposed to serve as the substrate entry gate.

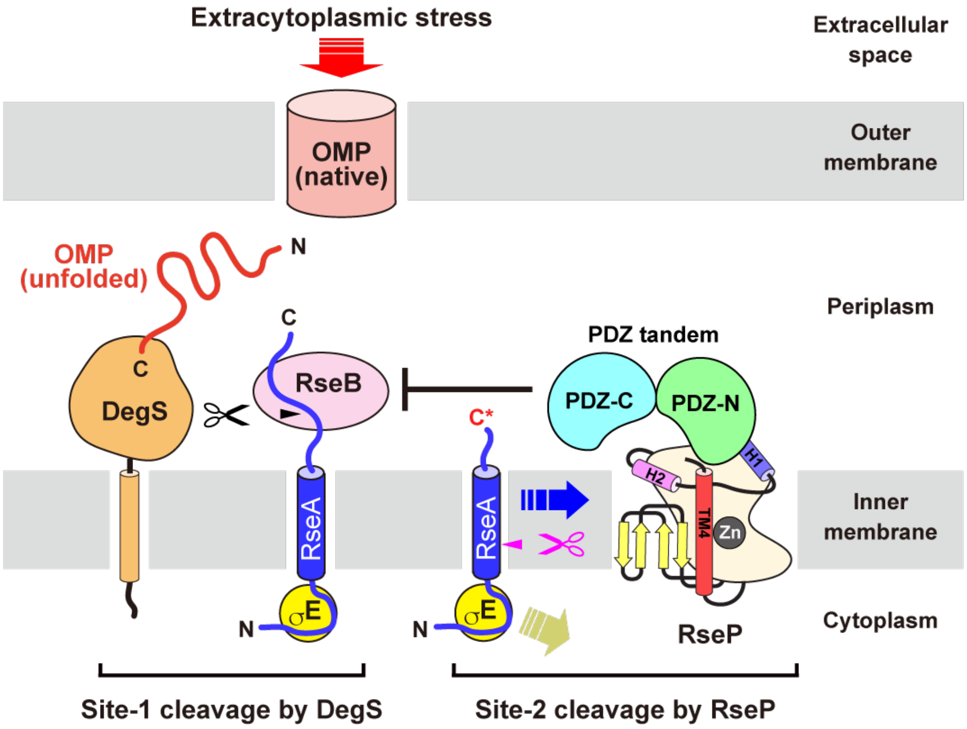

**Figure S2. Involvement of RseP in the extracytoplasmic stress response in *E. coli***

In *E. coli*, exposure to extracytoplasmic stress such as heat and alkali stresses induces the two-step sequential cleavage of the anti-sigma factor RseA on the inner membrane to activate the transcription factor $\sigma^E$. Extracytoplasmic stress causes accumulation of unfolded or denatured outer membrane proteins (OMPs) as well as lipopolysaccharide (LPS) in the periplasm. The membrane-anchored serine protease DegS, the *E. coli* counterpart of site-1 protease (S1P), is activated by interaction with the C-terminus of the unfolded OMP. Cleavage of the periplasmic region of RseA by the activated DegS (site-1 cleavage), together with LPS-induced dissociation of the periplasmic protein RseB from RseA, results in the removal of the negative regulation by RseB. Subsequently, the intramembrane zinc metalloprotease RseP accommodates the DegS-cleaved form of RseA and performs intramembrane proteolysis (site-2 cleavage). As a consequence, the N-terminal portion of RseA is released from the inner membrane and further degraded to activate $\sigma^E$. It was proposed that the PDZ tandem of RseP sterically hinders entry of the full-length RseA complexed with RseB. In this study, structure-based mutational and cross-linking analyses have been conducted to address the question of how RseP accommodates the substrates after the site-1 cleavage using structural elements such as the PDZ tandem, the PCT region including the two helices (H1 and H2), TM4, and MREβ sheet.

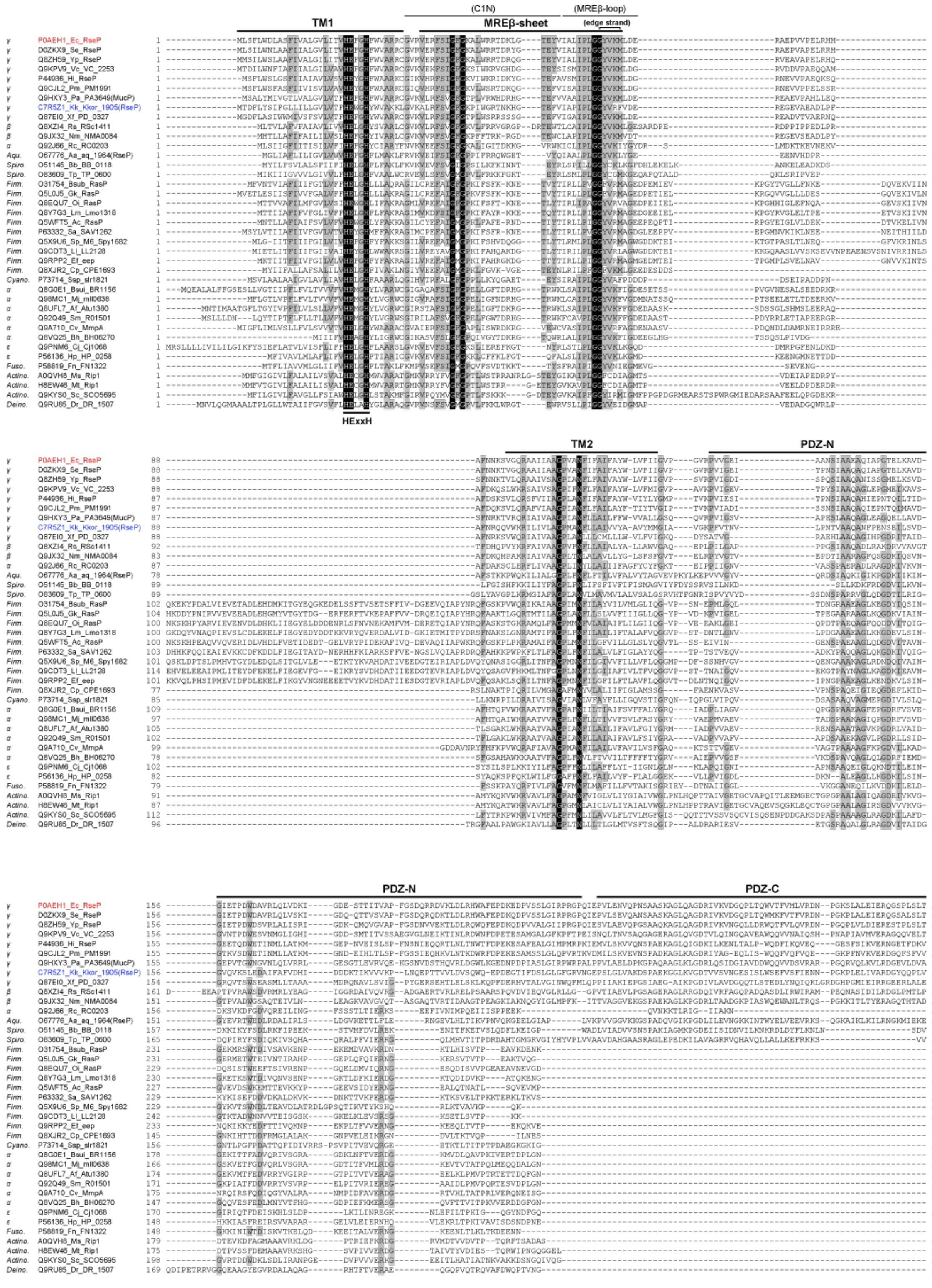

(C1N)
(MREβ-loop)
TM1
MREβ-sheet
(edge strand)
HExxH
TM2
PDZ-N
PDZ-N
PDZ-C

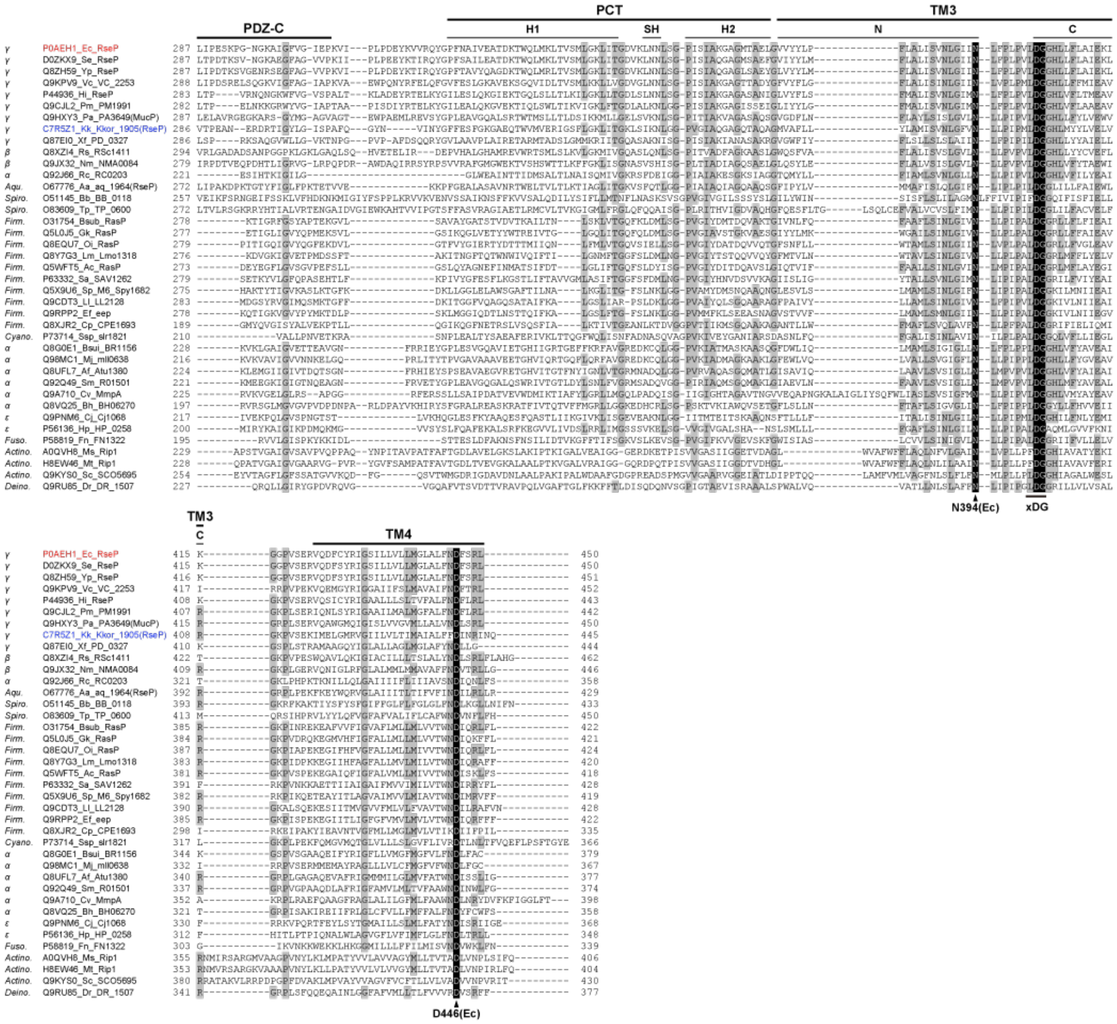

**Figure S3. Multiple alignment of bacterial RseP orthologues**

Amino acid sequences of bacterial RseP orthologues were aligned using Clustal W ver. 2.1 (http://www.clustal.org/clustal2/). The phylum (*Aqu*., Aquificae; *Spiro*., Spirochaetes; *Firm*., Firmicutes; *Cyano*., Cyanobacteria; *Fuso*., Fusobacteria; *Actino*., Actinobacteria; *Deino*., Deinococcus-Thermus) or class (*α*, *β*, *γ*, and *ε* for proteobacteria), the UniProtKB accession number, the abbreviation of species, and the gene name of each orthologue are shown on the left. Conserved and similar residues are boxed in black and gray, respectively. Red and blue indicate *Ec*RseP and *Kk*RseP, respectively. TM segments, structural elements and featured residues are based on those of *Ec*RseP. Previous studies suggested that *Ec*RseP possesses two intramembrane β-hairpins (labeled C1N and MREβ-loop) between TM1 and TM2. This study showed that the corresponding regions are integrated into a four-stranded β-sheet, labeled MREβ-sheet, in which the C-terminal residues, G68 to M73, form the edge strand as labeled.

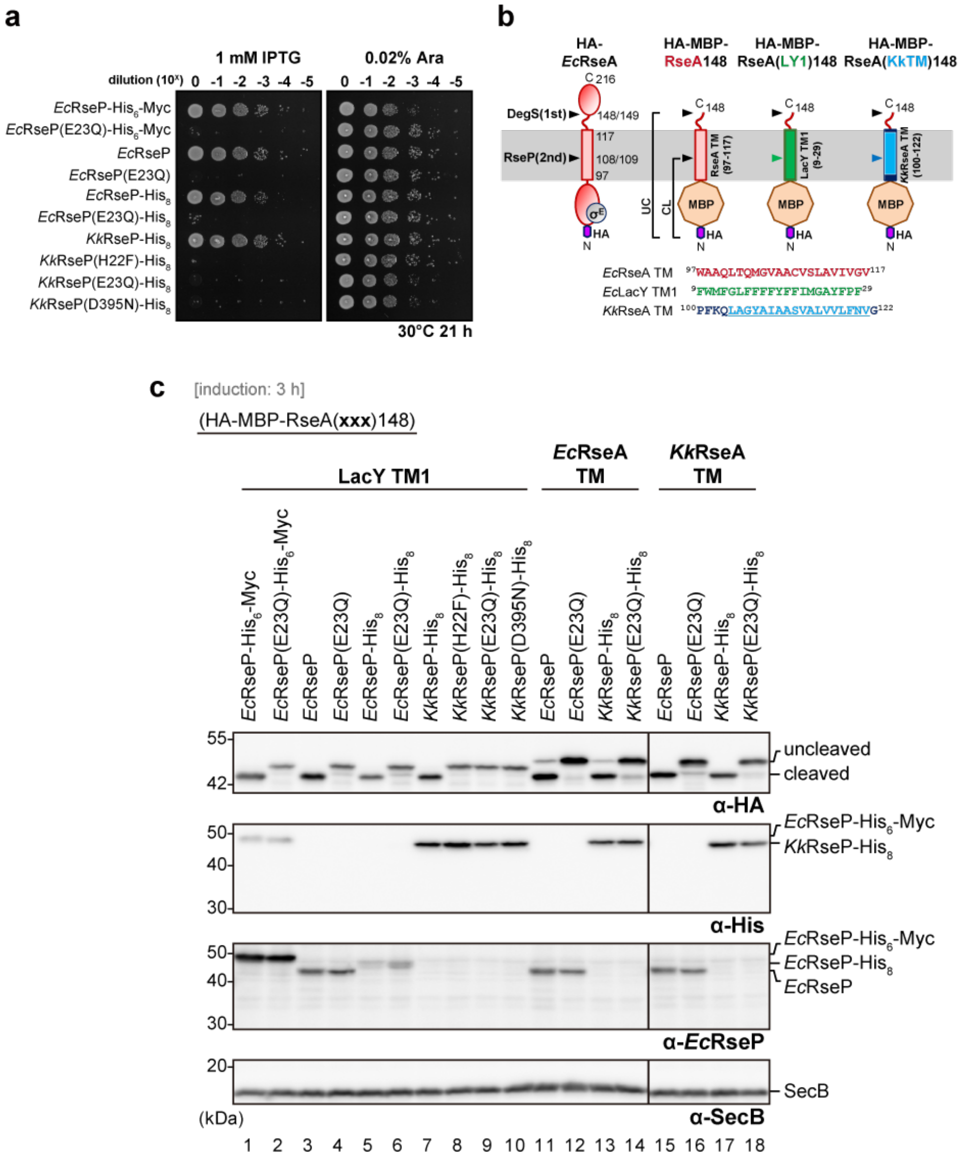

**Figure S4. Complementation and *in vivo* proteolytic activities of *Ec*RseP and *Kk*RseP with various C-terminal tag sequences**

**(a)** Complementation assay. Cultures of *E. coli* KK31 [Δ*rseP*/pKK6 (P$_{BAD}$-*rseP*)] cells harboring pKK11 (*Ec*RseP-His$_6$-Myc), pYH825 (*Ec*RseP with no detection tag), pYH820 (*Ec*RseP-His$_8$), pYH829 (*Kk*RseP-TEV-His$_8$) or a plasmid encoding tagged or untagged derivatives of *Ec*RseP or *Kk*RseP with the indicated mutations in their catalytic site motifs (*i.e.* HExxH and xDG) were serially diluted and spotted on L agar plates containing 1 mM IPTG (left) or 0.02% L-arabinose (right). The plates were incubated at 30°C for 21 h. A representative result of three biological replicates is shown. Note that whereas the pKK11 and its derivative encoding *Ec*RseP(E23Q)-His$_6$-Myc had the native SD sequence for *Ec*RseP, the others had a mutant form of the SD sequence (improved SD) for increased expression. **(b)** Schematic representation of the model substrates used in cleavage assays. Established model substrates have the RseA TM region or the first transmembrane region of LacY (LY1) (*19*), and a newly constructed derivative has a predicted TM region of *Kk*RseA. The amino acid sequences of the RseA TM region, LY1, and the predicted TM region of *Kk*RseA (underlined) are shown at the bottom. **(c)** *In vivo* substrate cleavage assay. *E. coli* KA306 cells harboring pYH124 (HA-MBP-RseA(LY1)148), pKA65 (HA-MBP-RseA148) or pYH817 (HA-MBP-RseA(KkTM)148) (labels, underlined) in addition to one of the indicated *Ec*RseP and *Kk*RseP-expressing plasmid were grown at 30°C in M9-based medium containing 1 mM IPTG and 1 mM cAMP for 3 h. Proteins were analyzed by Laemmli SDS-PAGE and immunoblotting. The results indicate that the C-terminally-attached tags had little observable effect on the intrinsic proteolytic activities of *Ec*RseP and *Kk*RseP, although these tags appear to slightly affect the stability of the proteins. Cytoplasmic protein SecB serves as a loading control. A representative result of three biological replicates is shown.

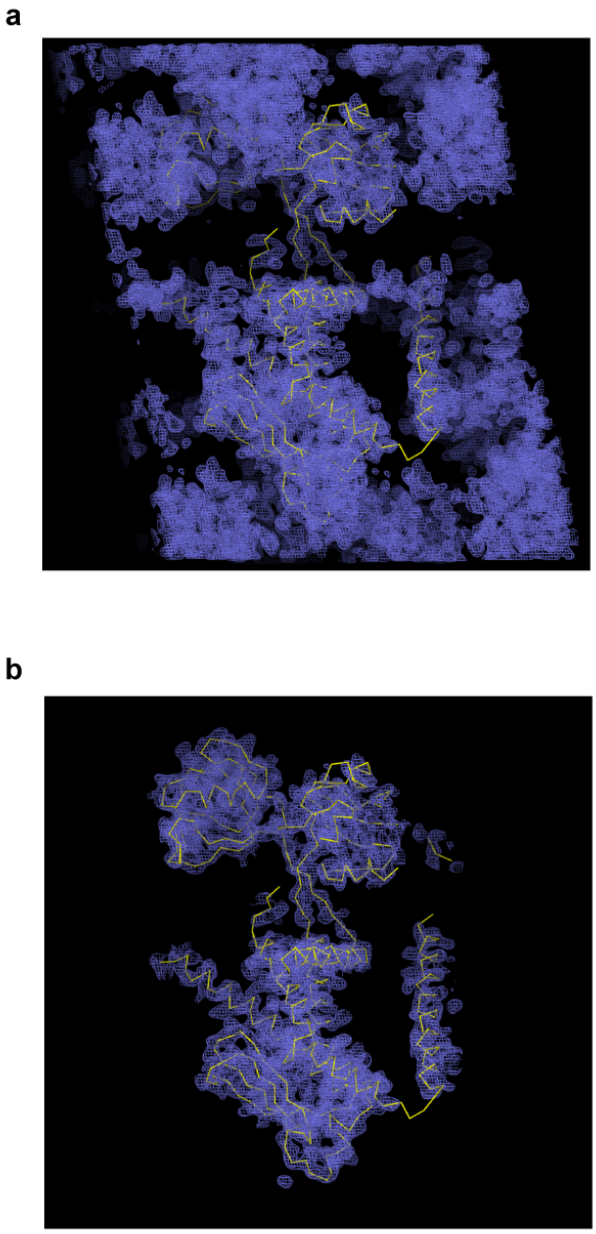

**Figure S5. Electron density map of *Kk*RseP**

**(a)** Electron density map calculated from the MR-SAD phasing and density modification. The electron density map of the *Kk*RseP crystal belonging to the space group *P*1 is contoured at 1.0 σ together with the Cα trace of the final model. **(b)** Electron densities contoured at 1.0 σ and within 2.0 Å of any atom in the final model.

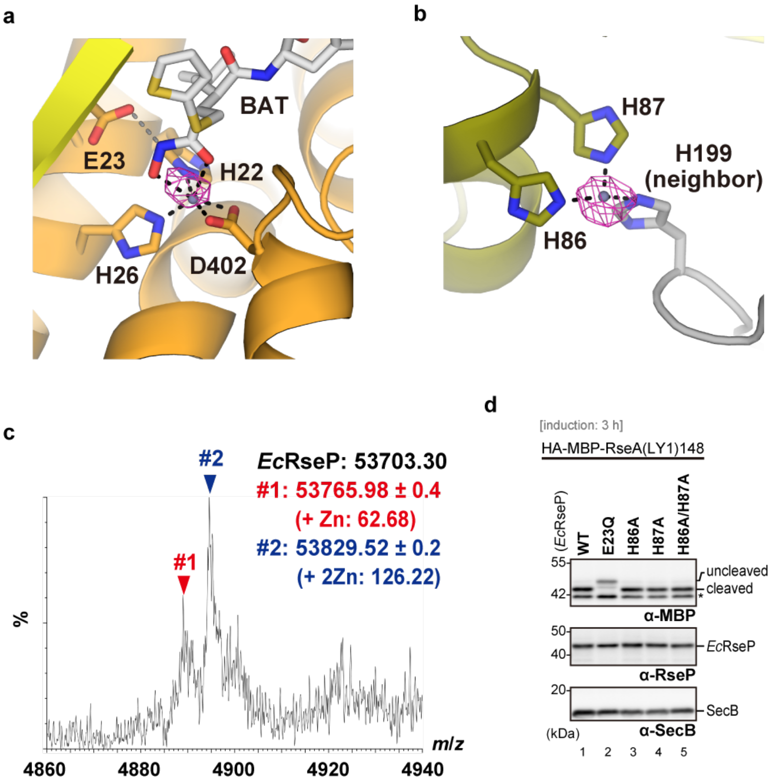

**Figure S6. Zinc-binding sites identified in the crystal structure of *Ec*RseP**

**(a)** Binding site of the catalytic zinc ion. Batimastat (white) and the side chains of zinc-coordinating residues (H22, H26, and D402) and E23 are shown as stick models. Anomalous difference Fourier map contoured at 4σ is shown as a magenta mesh. The map was calculated using structure factors from diffraction data collected near the zinc-absorption edge and phases calculated from the final model of *Ec*RseP. **(b)** Binding site of the secondary zinc ion in the cytoplasmic domain. The secondary zinc ion is coordinated by H86 and H87 on the same *Ec*RseP molecule and by H199 on the crystal lattice neighbor. Side chains are shown as stick models. The anomalous difference Fourier map is calculated and shown similarly as in **(a)**. **(c)** NanoESI mass spectrum of *Ec*RseP obtained under a “near native” condition (200 mM ammonium acetate containing 0.5 % (w/v) C8E4, pH 7.4). Ions at 11+ charge state are indicated. Major peaks #1 and #2 correspond to molecular masses of 53766.0 ± 0.4 and 53829.5 ± 0.2, respectively, confirming that two $Zn^{2+}$ are bound in the majority of the protein population. **(d)** *In vivo* substrate cleavage assay. *E. coli* KA306 cells harboring pYH124 (HA-MBP-RseA(LY1)148) plus pYH825 (*Ec*RseP) or its derivative encoding the indicated mutant were grown and analyzed as in Fig. S4c. An asterisk indicates endogenous MBP protein. A representative result of three biological replicates is shown.

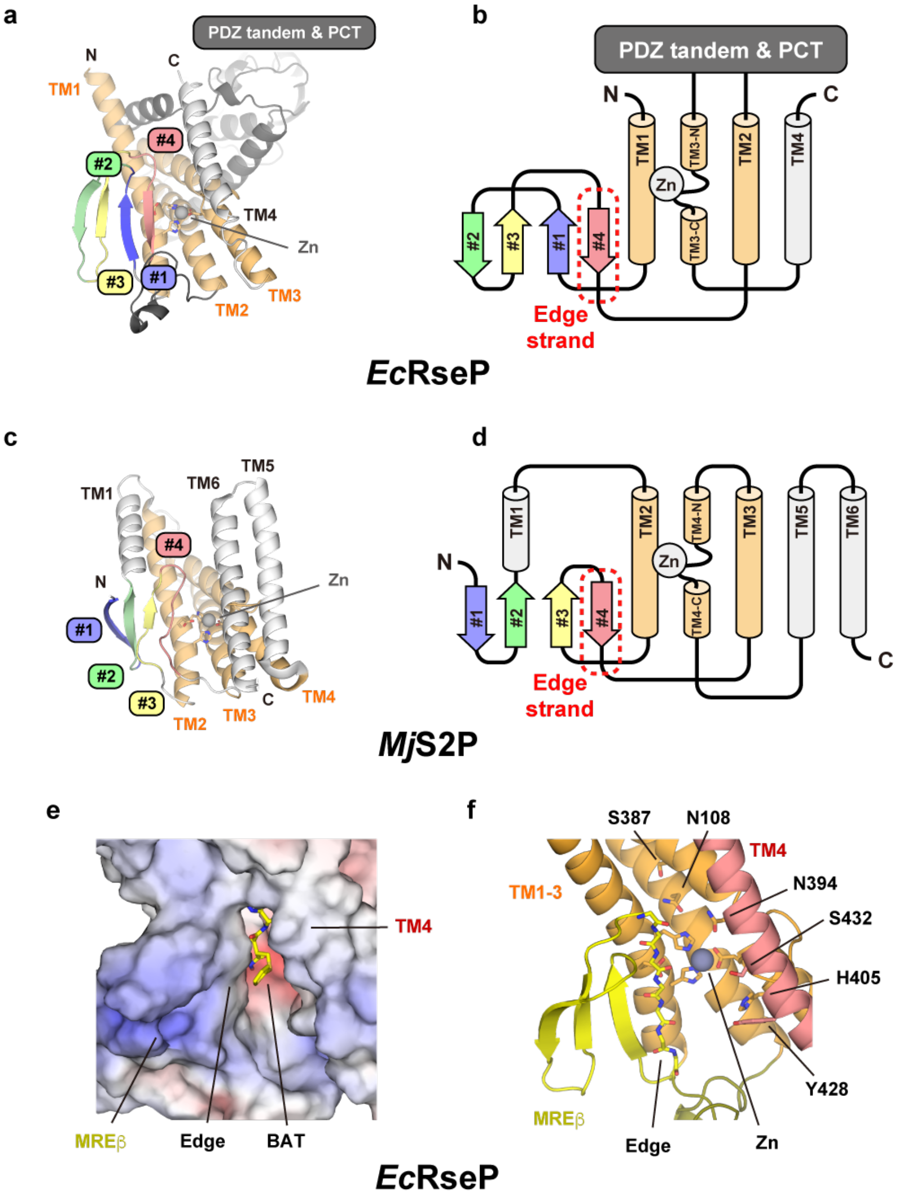

**Figure S7. Conservation of the MREβ sheet**

**(a)** Ribbon model of *Ec*RseP. TM1-3 constituting the conserved core are colored orange. The residues coordinating the zinc ion (gray sphere) or involved in catalysis are shown as stick models. β strands are colored uniquely. **(b)** Topology diagram of *Ec*RseP. All four strands are between TM1 and 2. Strand 4 corresponds to the edge strand expected to bind with the substrate. The domain organization and folding are consistent between *Ec*RseP and *Kk*RseP. **(c)** Ribbon model of the TM core domain of *Mj*S2P (PDB ID: 3B4R, chain A) (*26*). In *Mj*S2P, TM2-4 constitute a conserved core. **(d)** Topology diagram of *Mj*S2P. In contrast to *Ec*RseP, strands 1 and 2 are present upstream of TM1 while strands 3 and 4 are between TM2 and 3. However, the strand orientations are identical to those in *Ec*RseP. **(e)** Electrostatic surface potential of *Ec*RseP. The solvent excluded surface is colored according to the electrostatic potential (-10 to 10 kT/e) with negative potential in red and positive potential in blue. The two zinc ions and batimastat were omitted from the calculation of the potential. The bound batimastat is shown as a stick model. **(f)** Ribbon model of *Ec*RseP in the same view as that of the surface model in **(e)**. Batimastat is omitted to visualize the binding site. The side chains of the polar residues inside the batimastat-binding compartment or the residues constituting the active center are shown as stick models. The backbone of the edge strand on the MREβ-sheet is also shown as a stick model. As the MREβ-sheet possesses several basic residues on the cytoplasmic side, a hydrophilic cavity leading to the batimastat-binding compartment is formed within the TM domain as shown in **(e)**. Furthermore, the surface potential also indicates that TM4 separates the hydrophilic interior of the batimastat-binding compartment from the hydrophobic milieu of the lipid bilayer.

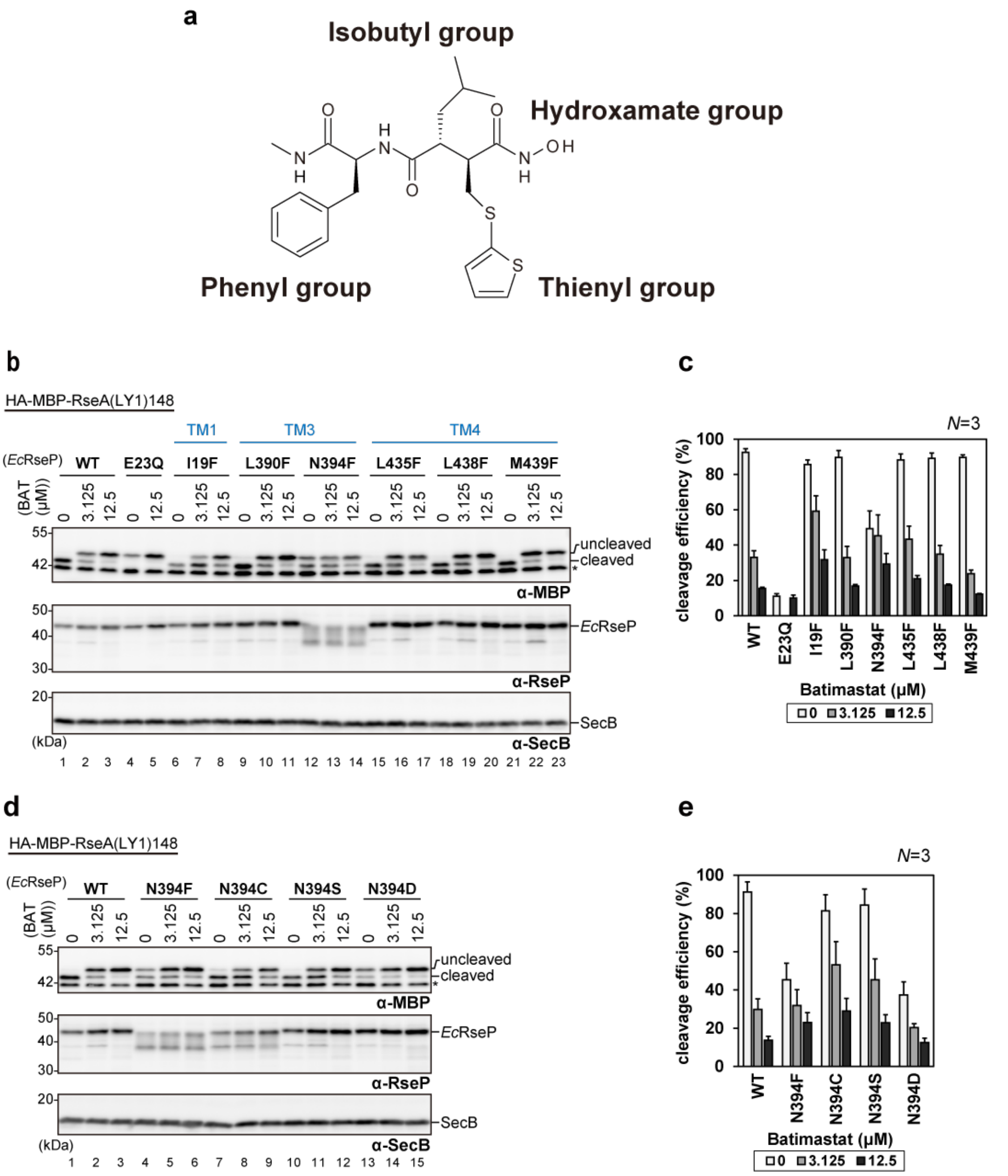

**Figure S8. *In vivo* batimastat sensitivity assay for *Ec*RseP mutants**

**(a)** Chemical structure of batimastat. **(b-d)** *E. coli* YH2902 (Δ*acrA* background (*27*)) cells harboring two plasmids, one encoding a model substrate HA-MBP-RseA(LY1)148 (pYH124) and the other encoding *Ec*RseP (pYH825) or its derivative, were treated with 0, 12.5 or 3.125 μM of batimastat at 30°C for 10 min (the mock sample (0 μM batimastat) received only the DMSO solvent). After incubation for 1 h in the presence of 1 mM IPTG, proteins were TCA-precipitated and analyzed by Laemmli SDS-PAGE and immunoblotting. **(b, d)** Gel images of the immunoblotting results. The region of mutation is labeled in blue. An asterisk indicates endogenous MBP protein. **(c, e)** Quantified cleavage ratio of each condition. Bar plots show means ± S.D. from three biological replicates.

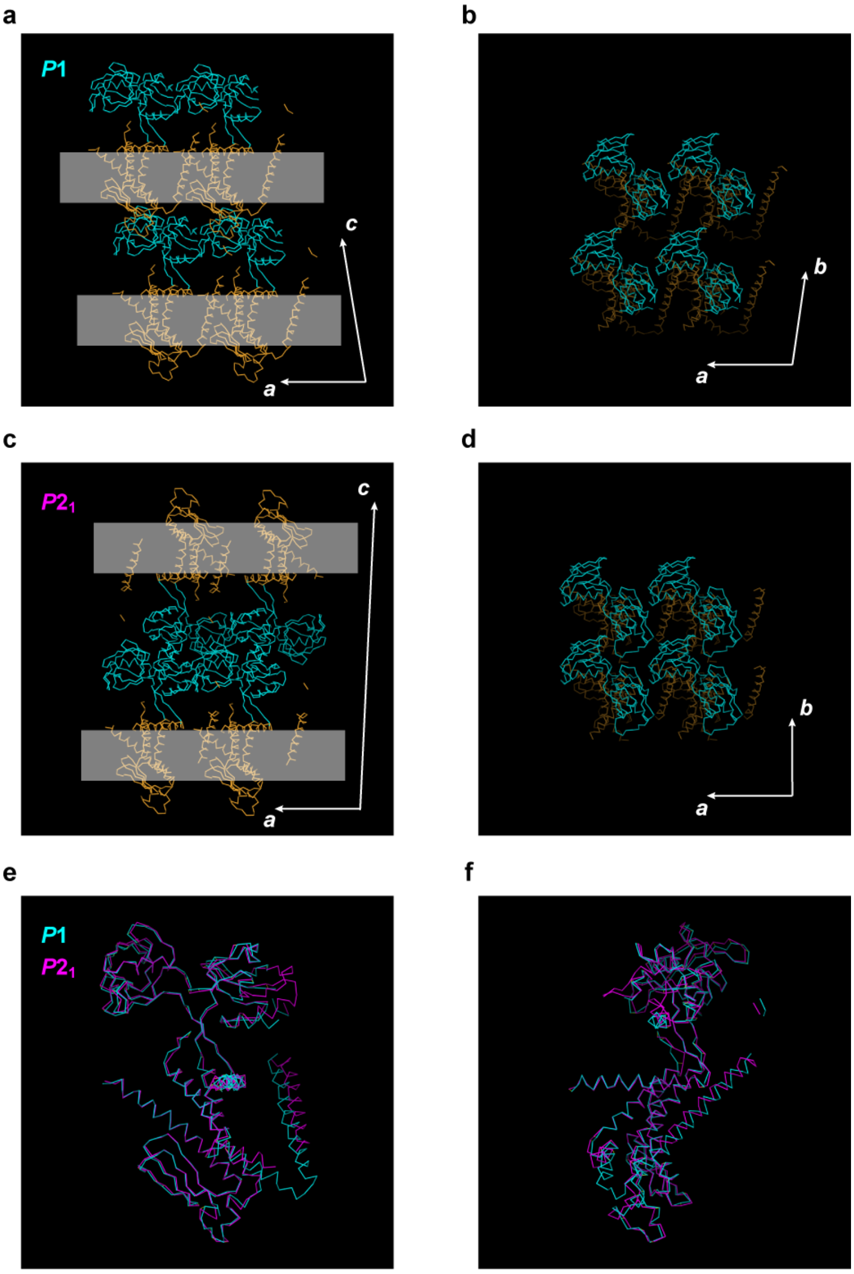

**Figure S9. Crystal packing of *Kk*RseP**

**(a, b)** *P*1 crystal. **(c, d)** $P2_1$ crystal. Each model is shown as Cα traces (orange). The PDZ tandem is colored cyan. The presumed lipid bilayer region is indicated by white transparent bands. The crystallographic axes are shown as white arrows. **(a)** In the *P*1 crystal form, 2D arrays of *Kk*RseP align along the *c*-axis in a head-to-tail orientation through interaction between the PDZ tandem and the cytoplasmic domain. **(c)** In the $P2_1$ crystal form, 2D arrays align along the *c*-axis in a head-to-head orientation through alternating interactions between the PDZ tandems or between cytoplasmic domains. **(b, d)** *Kk*RsePs crystal packing along the *a*-axis on the 2D array is similar between the two crystal forms. **(e, f)** Superposition of the two *Kk*RseP crystal structures (*P*1 in cyan, $P2_1$ in magenta). The main chain structure is almost identical between the two forms except for disorder in the PDZ-N and the linker between TM3 and 4. The orientation of TM4, which interacts with the neighboring molecule, is also slightly different between the two crystal forms.

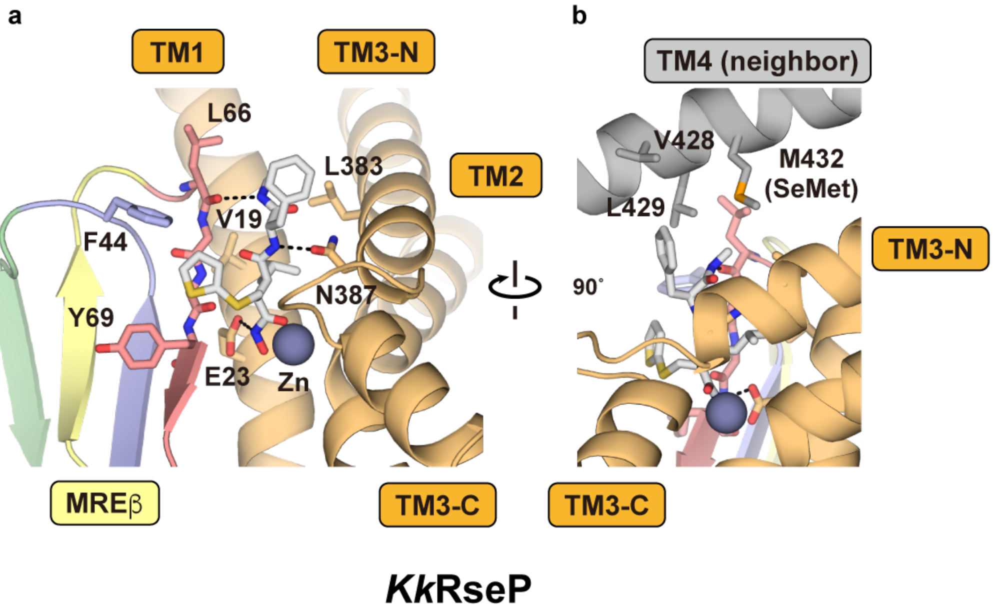

**Figure S10. Binding mode of batimastat to *Kk*RseP**

**(a)** Close-up view of the batimastat-binding site. Batimastat and the residues in direct contact with batimastat are shown as stick models. N387 corresponds to N394 in *Ec*RseP, which was demonstrated to contribute to efficient substrate cleavage in the present study. Each of the four strands constituting the MREβ sheet are shown in unique colors as in Fig. S5. The zinc ion in the active site is shown as a sphere. **(b)** Interactions between TM4 and the crystal packing neighbor. The model is shown similarly to those in **(a)**. TM4 residues located within 5 Å of batimastat are shown as stick models.

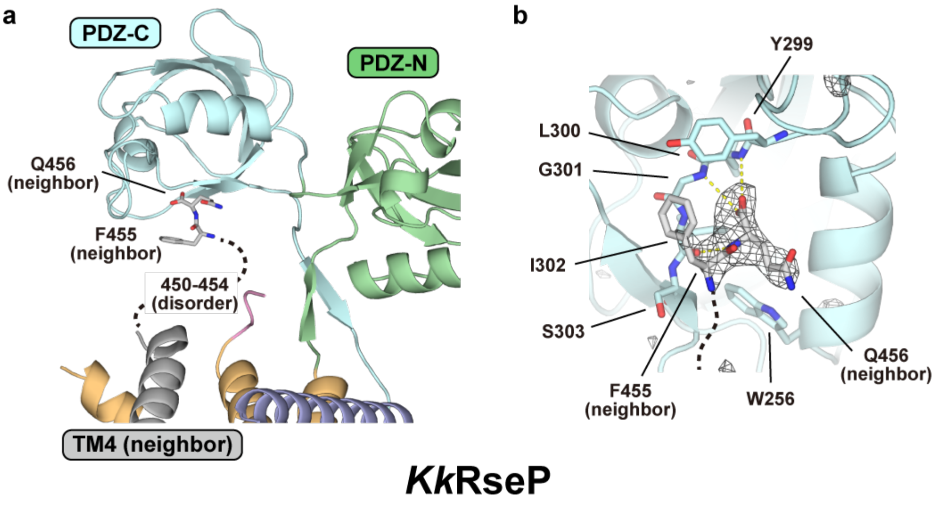

**Figure S11. Accommodation of the *Kk*RseP C-terminus into the PDZ-C domain**

**(a)** The full-length *Kk*RseP from the $P2_1$ crystal form. TM4 of the neighboring molecule, colored gray, is accommodated into the cleft between TM1 and TM3. In addition, F455 and Q456 at the C-terminus, shown as stick models, are observed in proximity to PDZ-C whereas residues 450-454 are disordered. **(b)** Close-up view of the C-terminus accommodation site on PDZ-C in *Kk*RseP from the $P2_1$ crystal form. F455 and Q456, together with the surrounding residues on PDZ-C, are shown as stick models. The $F_o$-$F_c$ map contoured at 3σ is shown with gray mesh. The residual electron density suggested that F455 and Q456 from the crystal packing neighbor are accommodated into PDZ-C. The residues Y299 to S303 correspond to the carboxylate-binding loop, which is generally present in the PDZ domains. The terminal carboxyl group of Q456 is presumed to form hydrogen bonds with the main chain N-H groups on this loop, as indicated by yellow dotted lines.

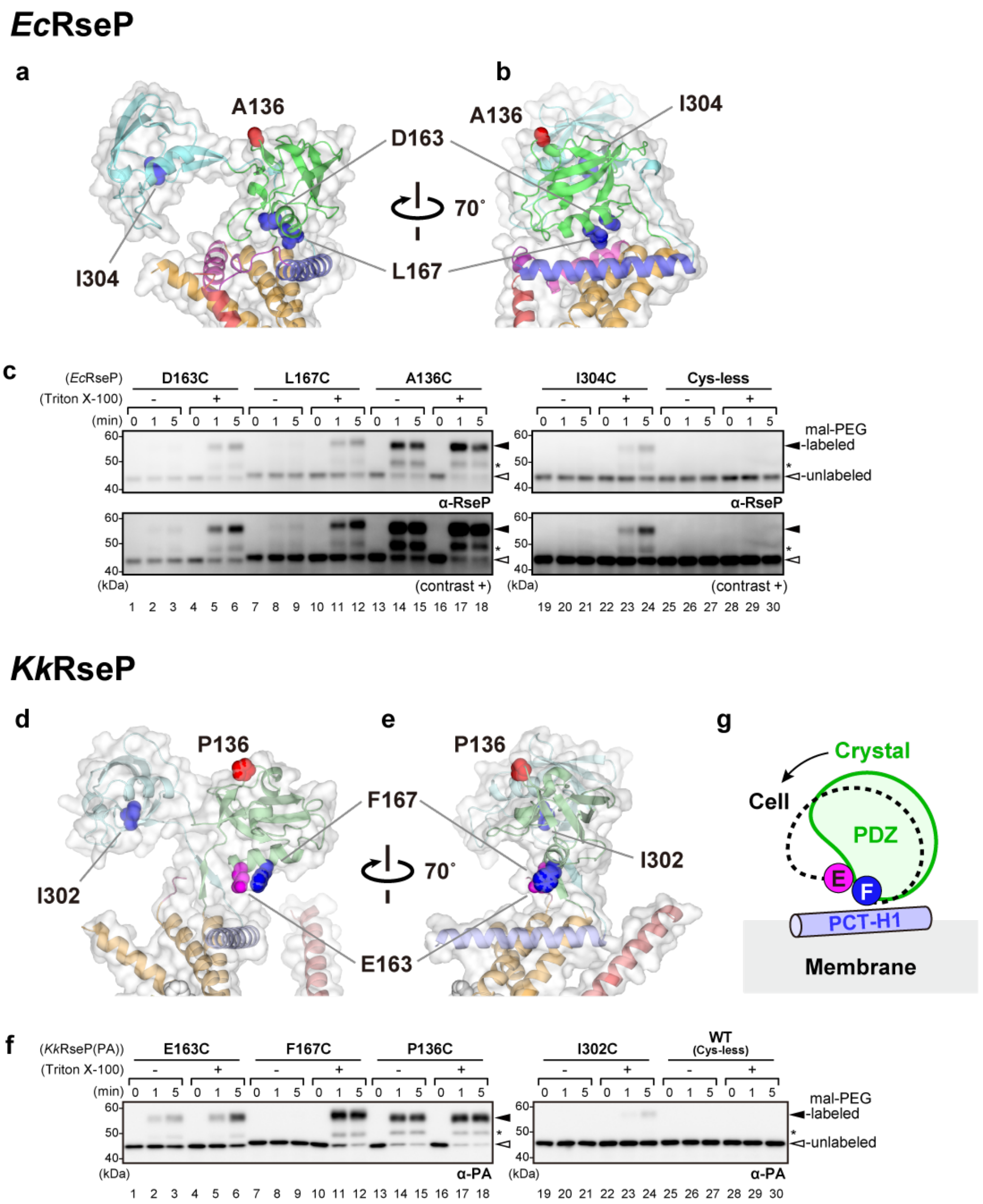

**Figure S12. mal-PEG accessibility assay for the PDZ tandem**

**(a, b)** Mapping mutation sites onto the crystal structure of *Ec*RseP. The crystal structure of *Ec*RseP is shown as a ribbon model with a transparent surface. The side chains of the mutated residues are shown as sphere models. Residues modified without detergent Triton X-100 (A136) are colored red. The three residues modified upon addition of the detergent (D163, L167, I304) are colored blue. L167 on PDZ-N makes direct contacts with PCT-H1 while D163 is also deep in the PDZ pocket. I304 belongs to the carboxylate-binding loop of PDZ-C. **(d, e)** Mapping mutation sites onto the crystal structure of *Kk*RseP. The structure is shown as in **(a, b)**. Cysteine residues were introduced into the sites of the four residues corresponding to the mutation sites in *E. coli*. The residue that was modified slightly without the detergent (E163) is colored magenta. **(c, f)** mal-PEG accessibility assay for the *Ec*RseP and *Kk*RseP Cys mutants. Spheroplasts of KK374 cells carrying a plasmid encoding a Cys-less EcRseP (pYH835), *Kk*RseP with internal PA14-tag (pYH838), or one of their single-cysteine derivatives were treated with 1 mM mal-PEG in the presence or absence of 2% Triton X-100 at 4°C for the indicated periods. TCA-precipitated proteins were analyzed by Laemmli SDS-PAGE and immunoblotting. 'contrast +' indicates a signal-enhanced image. Filled arrowheads indicate the mal-PEG-labeled forms of the RseP derivatives. An asterisk indicates RseP derivatives modified with a minor mal-PEG component with a smaller mass, as reported previously (*67*). A representative result of two biological replicates is shown. **(g)** Conformation of the *Kk*RseP PDZ tandem in the crystal and on the cell membrane. In the crystal structure of *Kk*RseP, the PDZ tandem adopts an open conformation, as indicated in green. However, the mal-PEG assay indicates that the *Kk*RseP PDZ tandem on the cell membrane adopts a relatively closed conformation, as indicated with a black dotted line, in which F167 (blue circle) is expected to be located in closer proximity to the PCT-H1 (light-blue column). The partial modification of E163 (magenta circle) in the absence of detergent indicates that the *Kk*RseP PDZ tandem is not as close to the PCT region as the *Ec*RseP PDZ tandem both in the crystal and on the cell membrane.

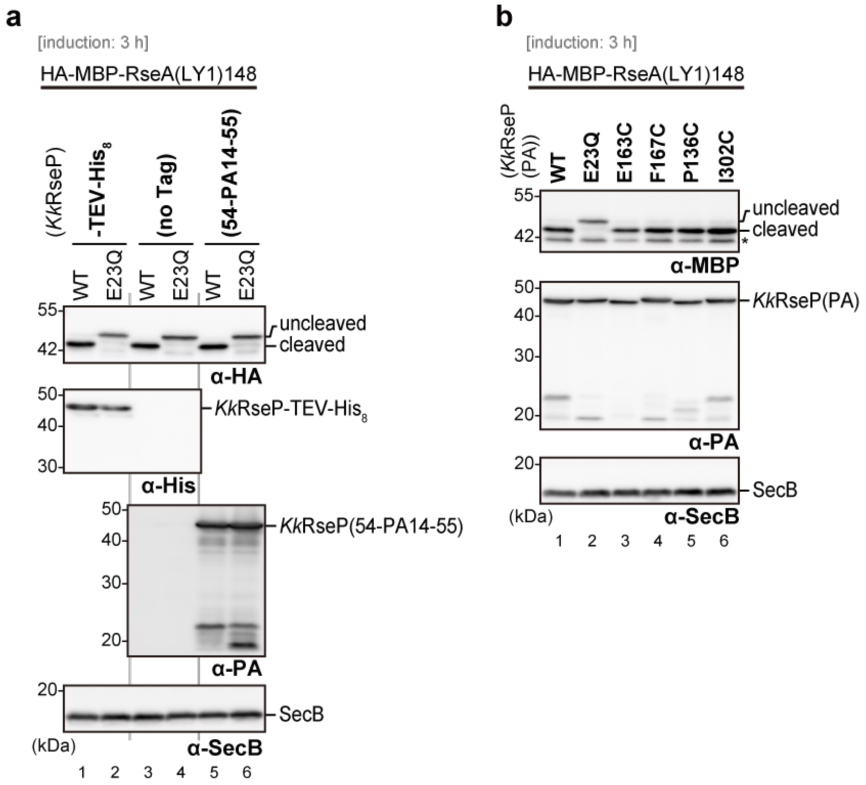

**Figure S13. *In vivo* substrate cleavage assay for *Kk*RseP derivatives**

*E. coli* KA306 cells harboring pYH124 (HA-MBP-RseA(LY1)148) plus pYH829 (*Kk*RseP-TEV-His$_8$), pYH833 (tag-less *Kk*RseP), pYH838 (*Kk*RseP(K54-PA14-H55)) or their derivatives were grown and analyzed as in Fig. S4c. **(a)** Proteolytic activity of the *Kk*RseP possessing an internal PA14 tag between Lys-54 and His-55. **(b)** Proteolytic activity of the Cys-introduced *Kk*RseP mutants with internal PA14 tag. An asterisk indicates endogenous MBP protein. A representative result of two **(a)** or three **(b)** biological replicates is shown, respectively.

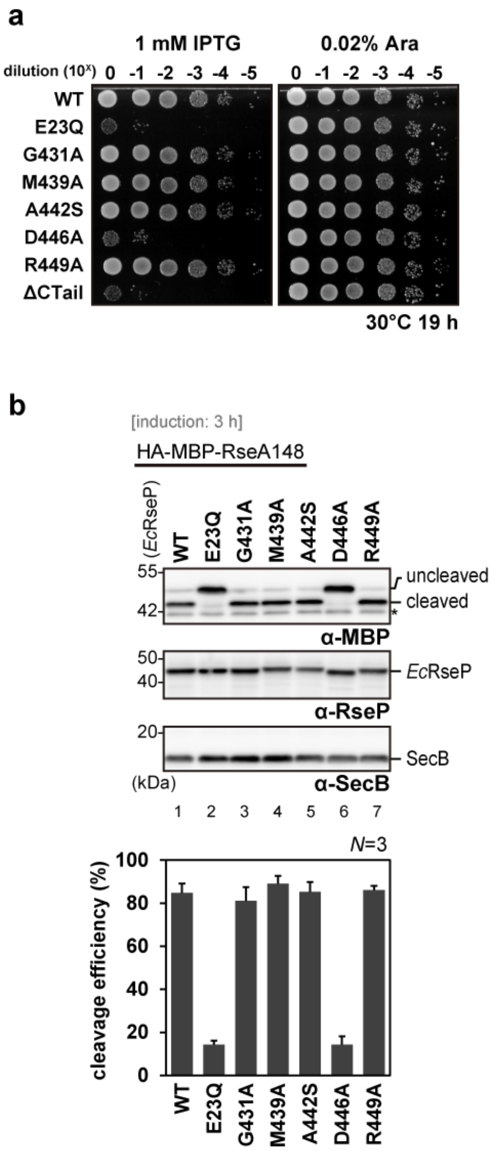

**Figure S14. Complementation and *in vivo* proteolytic activities of *Ec*RseP derivatives having a mutation to conserved residues in TM4**

**(a)** Complementation assay. Cultures of *E. coli* KK31 cells harboring pYH825 (*Ec*RseP) or its derivative were serially diluted and spotted on L agar plates containing 1 mM IPTG (left) or 0.02% L-arabinose (right). The plates were incubated at 30°C for 19 h. A representative result of three biological replicates is shown. **(b)** *E. coli* KA306 cells harboring pKA65 (HA-MBP-RseA148) plus pYH825 (*Ec*RseP) or its derivatives were grown and analyzed as in Fig. S4c. An asterisk indicates endogenous MBP protein. The bar plot shows the means ± S.D. from three biological replicates.

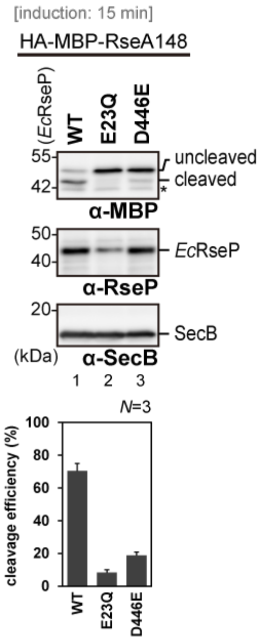

**Figure S15. *In vivo* substrate cleavage assay for the *Ec*RseP D446E mutant with short induction**

*E. coli* KA306 cells harboring pKA65 (HA-MBP-RseA148) plus pYH825 (*Ec*RseP) or its derivatives were used. After 2.5 h cultivation, cells were collected, pre-incubated at 30°C for 10 min, and induced for 15 min with 1 mM IPTG. Proteins were TCA-precipitated and analyzed by Laemmli SDS-PAGE and immunoblotting. The bar plot shows the means ± S.D. from three biological replicates. Cytoplasmic protein SecB serves as a loading control (α-SecB). An asterisk indicates endogenous MBP protein.

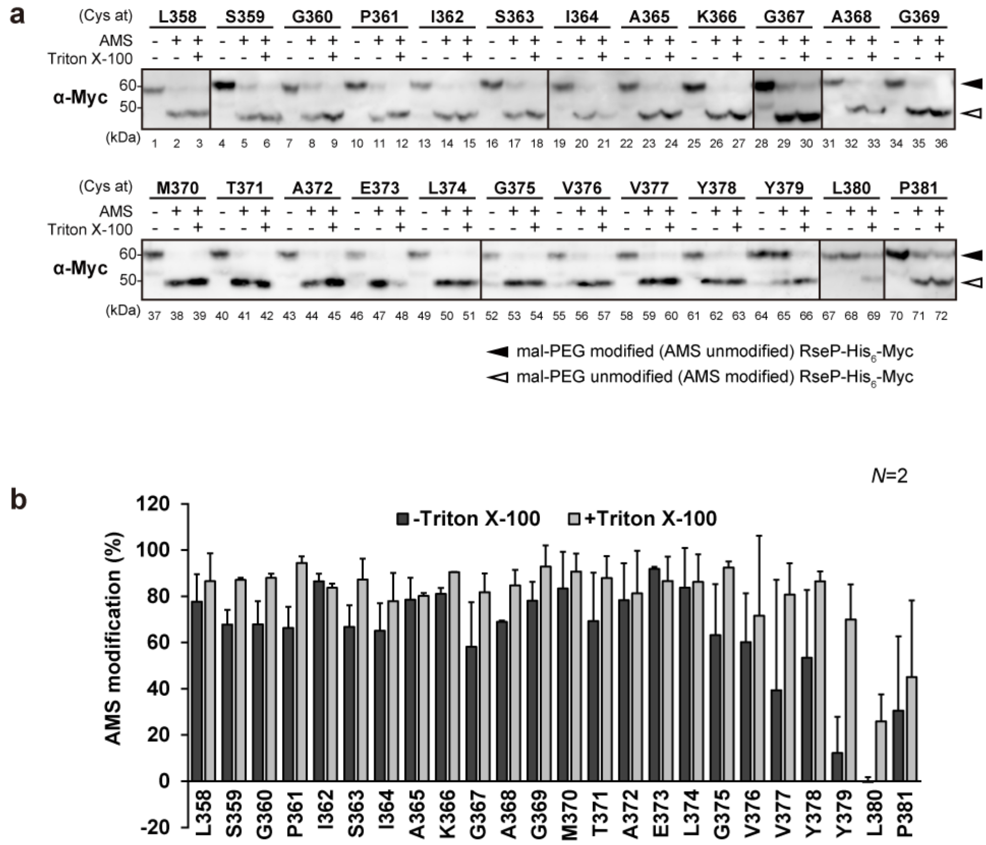

**Figure S16. Substituted cysteine accessibility analysis on the PCT-H2 region**

Spheroplasts prepared from KK374 cells harboring a plasmid encoding a derivative of *Ec*RseP-His$_6$-Myc possessing a single unique Cys residue at the indicated position (pTM101 derivatives) were treated with 1 mM AMS in the presence or absence of 1% Triton X-100 at 24°C for 5 min. After quenching AMS, proteins were precipitated with TCA, solubilized in 1% SDS, and treated with 5 mM mal-PEG at 37°C for 1 h. The samples were analyzed by Laemmli SDS-PAGE and anti-Myc immunoblotting. Filled and open arrowheads indicate mal-PEG-modified (AMS-unmodified) and mal-PEG-unmodified (AMS-modified) forms of RseP, respectively. **(a)** Gel images of the immunoblotting. **(b)** AMS modification (%) were calculated and shown on bar plots as means ± S.D. from two biological replicates.

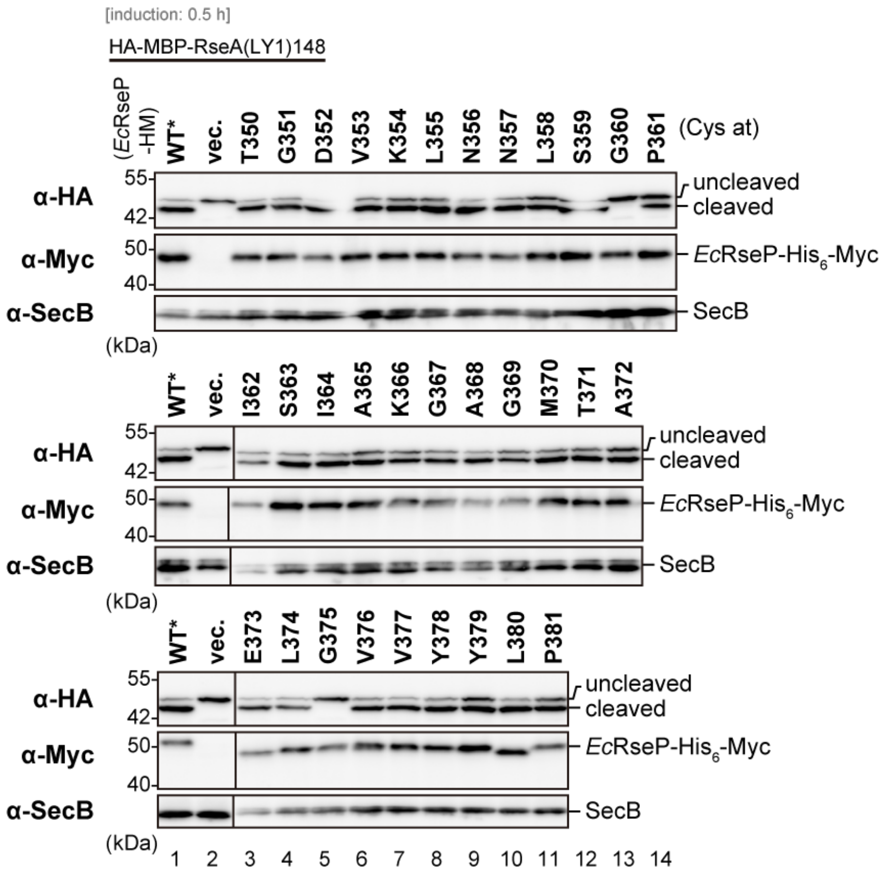

**Figure S17. Cysteine-scanning mutagenesis analysis on the PCT and adjacent regions**

Cleavage of the model substrate was examined with short-induction. KK211 cells harboring pYH20 (HA-MBP-RseA(LY1)148) plus pTM132 (*Ec*RseP(Cys-less)-His$_6$-Myc) or its derivatives were grown at 30°C in M9-based medium for 2.5 h. After addition of 1 mM IPTG and 5 mM cAMP, cells were further cultured for 0.5 h. Proteins were TCA-precipitated and analyzed by Laemmli SDS-PAGE and immunoblotting. Cleavage efficiency was shown graphically as the means ± S.D. from three biological replicates in Fig. 4. A representative result of three biological replicates is shown. WT* indicates the Cys-less derivative of WT RseP. Anti-HA and anti-SecB antibodies were pre-mixed and used to detect the model substrate and SecB simultaneously.

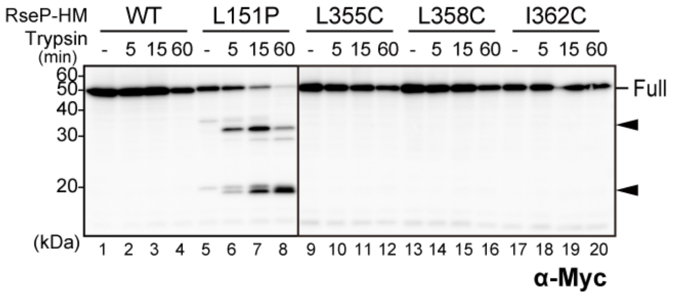

**Figure S18. Trypsin susceptibility of deregulated *Ec*RseP mutants**

Spheroplasts prepared from KK374 cells harboring pKK11 (*Ec*RseP-His$_6$-Myc) or its derivatives were incubated at 0°C with 2.5 μg/mL Trypsin for the indicated periods. '-' indicates a sample taken before Trypsin addition. TCA-precipitated proteins were analyzed by Laemmli SDS-PAGE and immunoblotting. *Full* indicates the intact RseP-HM protein. Filled arrowheads indicate tryptic fragments of RseP. A representative of two biological replicates is shown. L151P, a previously identified mutant with characteristic deregulation (*29*), shows increased Trypsin-susceptibility as the mutation causes large structural changes or unfolding of PDZ-N.

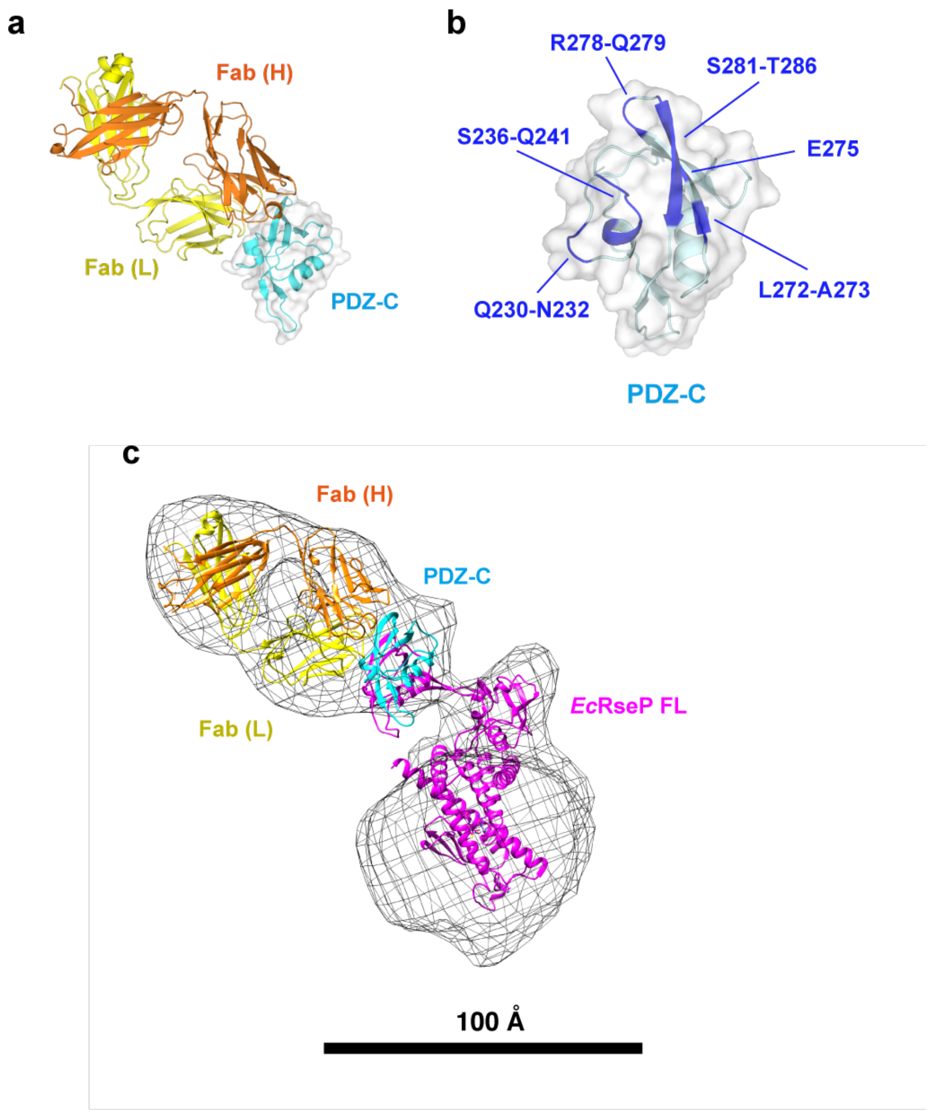

**Figure S19. Antibody-assisted structural analysis of *Ec*RseP**

**(a)** Crystal structure of *Ec*RseP PDZ-C in complex with the 12C7 Fab. The heavy and light chains of 12C7 Fab are shown as orange and yellow ribbon models, respectively. The PDZ-C domain is shown as a cyan ribbon model with a transparent surface. **(b)** Binding interface on PDZ-C. The PDZ-C domain from the Fab-complex structure is shown as a ribbon model in light cyan with a transparent surface. The residues in direct contact with the Fab (blue, labels) are scattered on the surface, indicating that 12C7 recognizes the ternary structure of PDZ-C. **(c)** 3D reconstruction model of the wild-type *Ec*RseP in complex with 12C7 Fab. The 3D model of class 4 (see Fig. S20) is shown as a mesh. In the 3D model, the TM domain is covered by a spherical detergent micelle, but the two PDZ domains are clearly identified as protrusions from the sphere. As the 12C7 Fab recognizes the ternary structure of PDZ-C rather than a linear epitope, the orientation of the 12C7 Fab is presumed to also be fixed relative to PDZ-C in the 3D model from the negative-stain EM. The positions of the 12C7 Fab (orange and yellow) and PDZ-C (cyan) were approximated based on the characteristic shape of Fab. The resulting orientation of the complex placed the PDZ-C domain with the density of the EM map. However, the PDZ-C domain appears to be in a slightly different position when the crystal structure of full-length *Ec*RseP (*Ec*RseP FL, magenta) is superposed onto the EM map such that the TM and PDZ-N domains are within the density on the EM map. The difference in the position of PDZ-C indicates that the PDZ-C domain in the EM sample is slightly separated from the TM domain as compared to that in the crystal structure. The packing interactions in the crystal structure or the negative staining process for the EM analysis probably caused the discrepancy. Despite the differences in orientation, the domain arrangement of *Ec*RseP both in the EM sample and in the crystal structure positions the PDZ domains sitting near the membrane and above the active site.

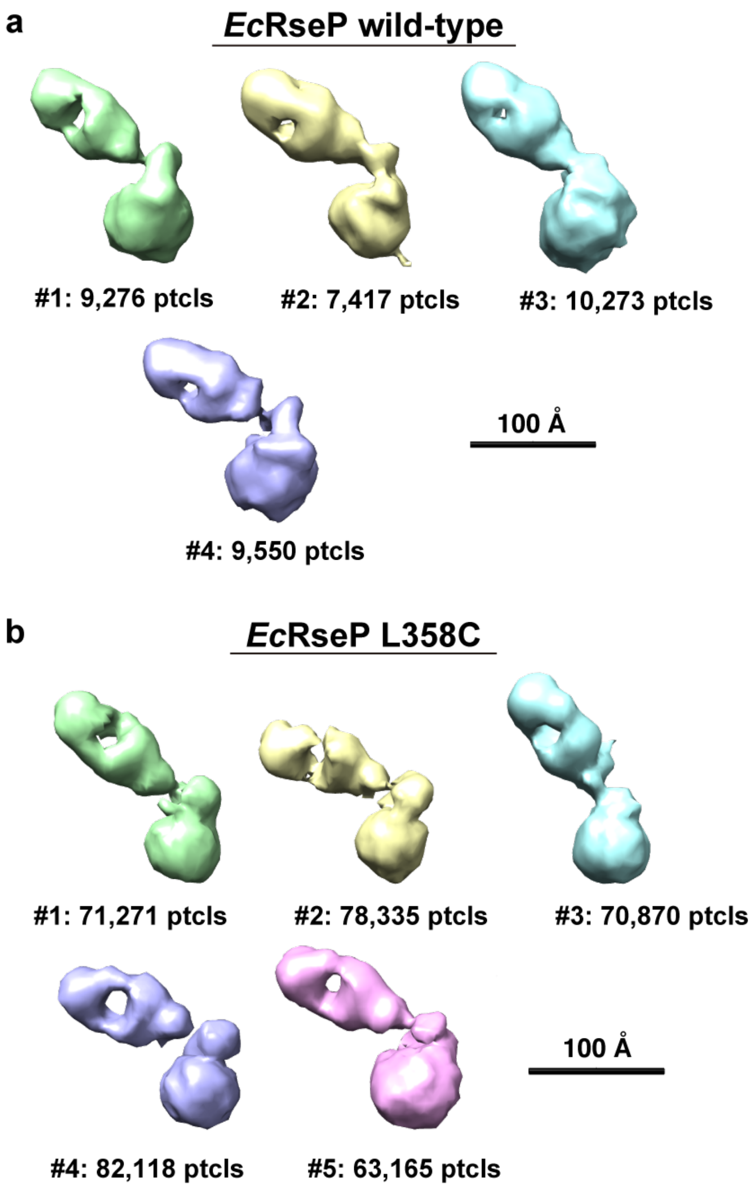

**Figure S20. 3D reconstruction models of *Ec*RseP complexed with 12C7 Fab**

**(a)** Fab-complex of the wild-type *Ec*RseP. Four different 3D models reconstructed from the 2D class averages are shown in different colors. Class 4 corresponds to the 3D model shown in Fig. S19c. **(b)** Fab-complex of the *Ec*RseP L358C mutant. Five different 3D models are shown as in **(a)**. The 3D models were aligned based on the position of the spherical micelle and the protrusion predicted to be PDZ-N. The numbers of particles used for each reconstruction are indicated below the model. The characteristic shape of the Fab aids in approximating the orientation. Compared to the orientation of Fab relative to the wild-type *Ec*RseP, the orientation of the Fab fluctuated to a larger extent relative to the L358 mutant. In addition, the Fab twists relative to *Ec*RseP based on the position of the central hole between the Fv and constant regions. All five models of the Fab-complex with the L358C mutant contain two protrusions from the spherical micelle that are interpreted to be the ternary structures of the individual PDZ-N and -C domains, but the orientation of PDZ-C fluctuated with the introduction of the L358C mutation in the PCT-loop region.

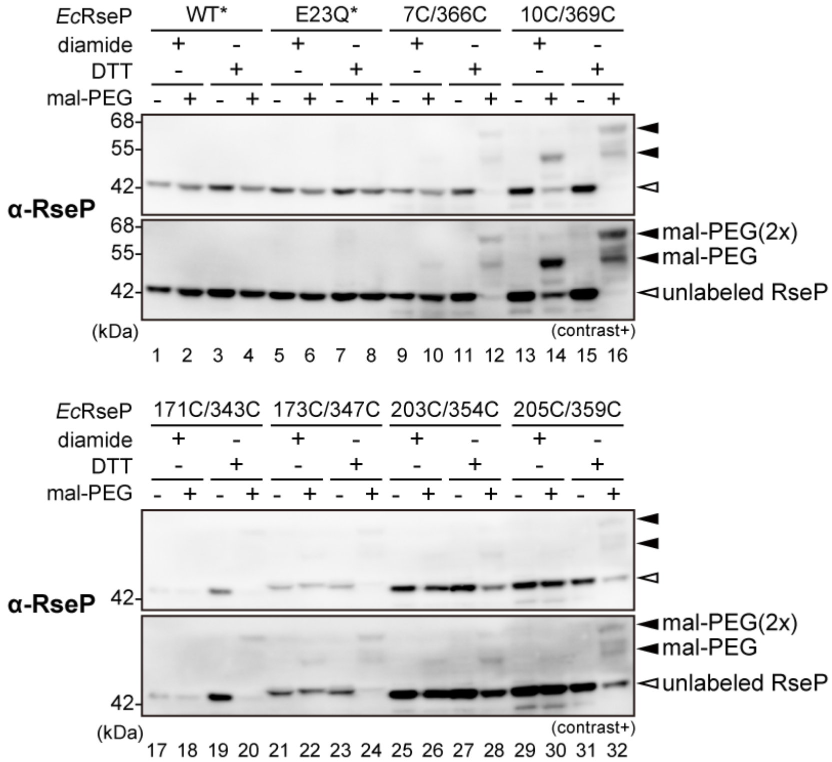

**Figure S21. Evaluation of disulfide-bond formation efficiency for double-Cys RseP mutants**

AD2544 cells harboring two plasmids, one encoding an RseP derivative with no or a pair of Cys residues (pYH835(Cys-less *Ec*RseP)-derivative) and the other encoding an model substrate HA-MBP-RseA(LY1)148 (pTM949), were used. Cells were cultured at 30°C for 30 min with 5 mM IPTG to induce expression of the RseP derivatives, washed, and cultured for an additional 30 min with 5 mM diamide or 10 mM DTT. Cells were then washed again and cultured for 30 min with 0.02% L-arabinose to induce a substrate expression and cleavage by RseP. TCA-precipitated proteins were solubilized in 1% SDS and subjected to modification with 0 or 1 mM mal-PEG at 37°C for 1 h to modify free thiols (The 0 mM mal-PEG samples received only the DMSO used as solvent for mal-PEG). Samples were then treated with 2ME and analyzed by Laemmli SDS-PAGE and immunoblotting. Open and filled arrowheads indicate the unmodified or mal-PEG-modified form (at one or two Cys residues) of RseP, respectively. No or little mal-PEG modification of the double-Cys RseP mutant proteins in the diamide-treated samples indicates that the Cys pair can form an interdomain disulfide bond either spontaneously or upon diamide-treatment. WT* and E23Q* indicate the Cys-less derivatives of WT and E23Q RseP, respectively. 'contrast +' indicates a signal-enhanced image. A representative result of two biological replicates is shown.

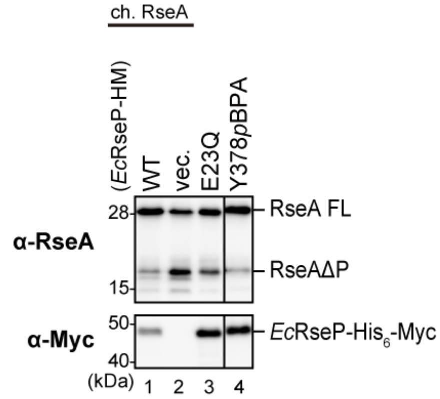

**Figure S22. Proteolytic function of *Ec*RseP(Y378*p*BPA)**

KA418 (Δ*rseP rseA*$^{+}$) cells harboring pEVOL-pBpF were transformed with pKK49 (*Ec*RseP-His$_6$-Myc, WT), pKA52 (E23Q) or a derivative of pKK49 with an amber mutation at Y378 in RseP. Cells were grown at 30°C for 4 h in M9-based medium supplemented with 0.5 mM *p*BPA. TCA-precipitated proteins were analyzed by Laemmli SDS-PAGE and immunoblotting. RseA FL and RseAΔP indicate the full-length and the DegS-cleaved form of RseA, respectively. Decrease in the accumulation level of RseAΔP in RseP(Y378*p*BPA)-expressing cells indicates that RseP(Y378*p*BPA) retained proteolytic activity. A representative result of three biological replicates is shown.

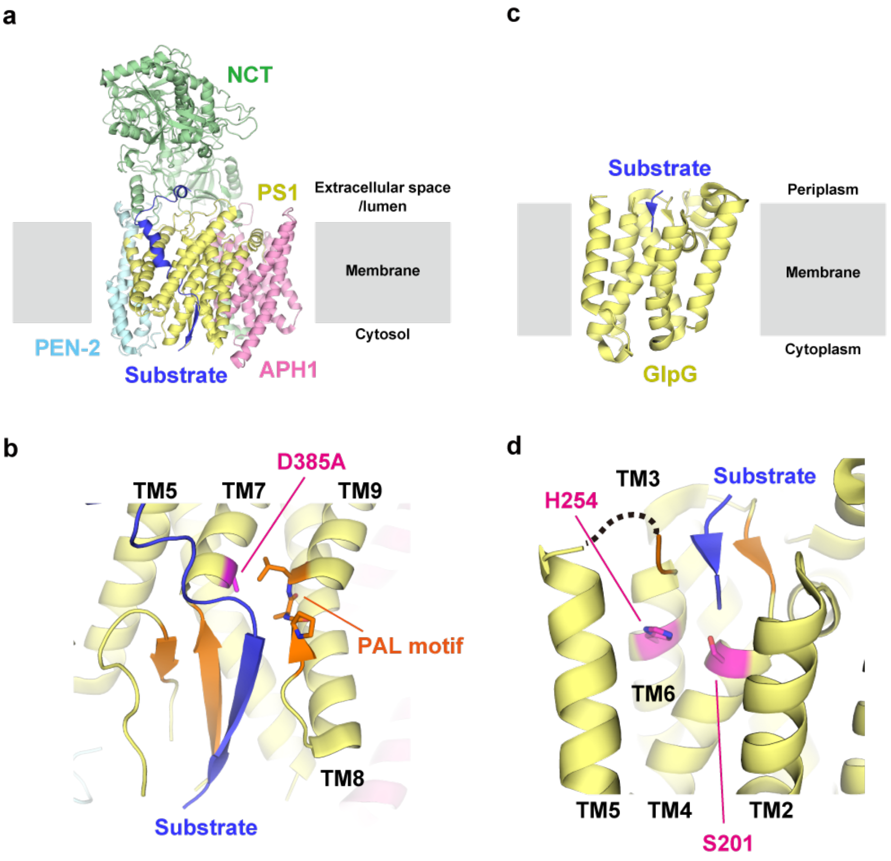

**Figure S23. Substrate unwinding modes in intramembrane proteases**

**(a, b)** Cryo-EM structure of human γ-secretase in complex with the Notch fragment (PDB ID: 6IDF) (*32*). **(a)** γ-Secretase is composed of four subunits: presenilin 1 (PS1), nicastrin (NCT), anterior pharynx-defective 1 (APH-1), and presenilin enhancer 2 (PEN-2). The Notch fragment indicated by 'Substrate' in blue is accommodated in the catalytic subunit PS1. **(b)** Close-up view around the active site. In this structure, the active site residue D385 was mutated to alanine as highlighted in magenta. The C-terminal region of Notch is unwound by forming a hybrid β-sheet with β-strands (orange) in the cytosolic loop between TM6 and 7 of PS1. The PAL motif between TM8 and 9, highlighted with stick models, appears to fix the Notch fragment as a clamp. **(c, d)** Crystal structure of *E. coli* GlpG in complex with the model substrate (PDB ID: 6PJ9) (*30*). **(c)** GlpG possesses six TM helices and accommodates the model substrate highlighted in blue in the cavity formed at the periplasmic side. **(d)** Close-up view around the active site. S201 and H254, highlighted in magenta, form the active center. The β-strands (orange) in the periplasmic loops between TM3 and 4 as well as between TM5 and 6 forms a parallel β-sheet with the substrate to fix it as a clamp.

### Supplementary Tables

**Table S1. Strains used in structural and biochemical analysis**

| Name | Genotype | Reference |
|---|---|---|
| BL21(DE3) | $F^-$, *ompT hsdS*$_B$($r_B^-$ $m_B^-$) *gal*(*λc*I 857 *ind*1 *Sam*7 *nin*5 *lacUV*5-T7*gene*1) *dcm*(DE3) | Merck |
| C43(DE3) | $F^-$, *ompT hsdS*$_B$($r_B^-$ $m_B^-$) *gal dcm* (DE3) | Lucigen |
| AD16 | Δ*pro-lac thi* / F' *lacI*$^q$ *Z*ΔM15 *Y*$^+$ *pro*$^+$ | (*68*) |
| KK211 | AD16, Δ*rseA*::*cat* Δ*rseP*::*kan* | (*17*) |
| AD2543 | AD16, Δ*rseA* | This study |
| AD2544 | AD16, Δ*rseA* Δ*rseP*::*kan* | This study |
| KA306 | AD16 Δ*rseA* Δ*rseP*::*kan* Δ*clpP*::*cat* | (*24*) |
| YH2898 | AD16, Δ*rseA* Δ*acrA* | This study |
| YH2902 | AD16, Δ*rseA* Δ*acrA* Δ*rseP*::*kan* | This study |
| KK31 | AD16, Δ*rseP*::*kan* Δ(*srl-recA*)*306*::Tn*10* /pKK6 ($P_{BAD}$-*rseP*) | (*14*) |
| BW25113 | $F^-$, *rrnB* Δ*lacZ4787* *hsdR514* Δ(*araBAD*)*567* Δ(*rhaBAD*)*568 rph-1* | (*60*) |
| JW2556 | BW25113, Δ*rseA*::*kan*, KEIO collection | (*59*) |
| JW0452 | BW25113, Δ*acrA*::*kan*, KEIO collection | (*59*) |
| MC4100 | *araD139* Δ(*argF-lac*)*U169 rpsL150 relA1 flbB5301 deoC1 ptsF25 rbsR* | (*69*) |
| CU141 | MC4100 / F'*lacI*$^q$ *lacZYA*$^+$ | (*70*) |
| KK374 | CU141, Δ*rseA*::*cat* Δ*rseP*::*kan* Δ*degS*::*tet* | (*62*) |
| KA418 | CU141, Δ*ompA* Δ*ompC* Δ*rseP*::*kan* | (*24*) |
| MC1061 | *araD* Δ(*ara-leu*)*7697* Δ(*codB-lacI*) *galK16 galE15 mcrA0 relA1 rpsL150 spoT1 mcrB9999 hsdR2* | (*71*) |
| CAG16037 | MC1061, Φλ[*rpoHP3-lacZ*] | (*72*) |
| AD2473 | CAG16037, Δ*ompA* Δ*ompC* Δ*rseP*::*kan* Δ*degS*::*tet* | (*21*) |

**Table S2. Plasmids used in structural and biochemical analysis**

| Name | Vector | Encoded proteins or descriptions | Reference |
|---|---|---|---|
| pET-21b | | pBR322-based vector; $P_{T7}$, $Amp^R$ | Merck |
| pGEX-2T | | pBR322-based vector encoding the schistosomal GST as a fusion partner; $P_{tac}$, $Amp^R$ | Cytiva |
| pBAD33 | | pACYC184-based vector; $P_{BAD}$, $Cm^R$ | (*73*) |
| pSTD689 | | pACYC184-based vector; $P_{lac}$, $Spc^R$ | (*74*) |
| pSTV29 | | pACYC184-based vector; $P_{lac}$, $Cm^R$ | Takara Bio |
| pTWV228 | | pBR322-based vector; $P_{lac}$, $Amp^R$ | Takara Bio |
| pUC118 | | pBR322-based vector; $P_{lac}$, $Amp^R$ | Takara Bio |
| pEVOL-pBpF | | p15A-derivative encoding mutant *M. jannaschii* aminoacyl-tRNA synthetase and suppressor tRNA for *p*BPA incorporation; $Cm^R$ | (*75*) |
| pCP20 | | FLP recombinase | (*76*) |
| pNY1452 | pUC118 | nSD, *Ec*RseP-TEV-$His_8$-Myc-PA | This study |
| pNY1432 | pET-21b | *Kk*RseP-TEV-$His_8$ | (*74*) |
| pNO1494 | pGEX-2T | GST-Thrombin-*Ec*PDZ tandem | This study |
| pNO2413 | pGEX-2T | GST-$His_8$-TEV-*Ec*PDZ-C | This study |
| pNY1480 | pGEX-2T | GST-$His_8$-TEV-*Kk*PDZ-C | This study |
| pSTD343 | pSTV29 | *lacI* | (*77*) |
| pKK6 | pBAD33 | *Ec*RseP | (*14*) |
| pKK11 | pTWV228 | nSD, *Ec*RseP-$His_6$-Myc (= HM) | (*14*) |
| pKK34 | pTWV228 | nSD, *Ec*RseP-HM, E23Q | (*14*) |
| pYH131 | pUC118 | nSD, *Ec*RseP-$His_8$ | This study |
| pYH819 | pUC118 | iSD, *Ec*RseP-$His_8$ | This study |
| pYH820 | pTWV228 | iSD, *Ec*RseP-$His_8$ | This study |
| pYH823 | pTWV228 | iSD, *Ec*RseP-$His_8$, E23Q | This study |
| pKK47 | pTWV228 | nSD, *Ec*RseP | (*66*) |
| pYK2 | pTWV228 | nSD, *Ec*RseP, E23Q | (*66*) |
| pYH825 | pTWV228 | iSD, *Ec*RseP | This study |
| pYH826 | pTWV228 | iSD, *Ec*RseP, E23Q | This study |
| pYH835 | pTWV228 | iSD, *Ec*RseP, Cys-less (= C33A/C427A) | This study |
| pTM910 | pTWV228 | iSD, *Ec*RseP, Cys-less, E23Q | This study |
| pYH846 | pTWV228 | iSD, *Ec*RseP, I19F | This study |
| pYH847 | pTWV228 | iSD, *Ec*RseP, L390F | This study |
| pYH848 | pTWV228 | iSD, *Ec*RseP, N394F | This study |
| pYH849 | pTWV228 | iSD, *Ec*RseP, L435F | This study |
| pYH850 | pTWV228 | iSD, *Ec*RseP, L438F | This study |
| pYH851 | pTWV228 | iSD, *Ec*RseP, M439F | This study |
| pYH855 | pTWV228 | iSD, *Ec*RseP, Cys-less, D163C | This study |
| pYH856 | pTWV228 | iSD, *Ec*RseP, Cys-less, L167C | This study |
| pYH857 | pTWV228 | iSD, *Ec*RseP, Cys-less, A136C | This study |
| pYH858 | pTWV228 | iSD, *Ec*RseP, Cys-less, I304C | This study |
| pYH870 | pTWV228 | iSD, *Ec*RseP, N394C | This study |
| pYH871 | pTWV228 | iSD, *Ec*RseP, N394S | This study |
| pYH872 | pTWV228 | iSD, *Ec*RseP, N394D | This study |
| pYH877 | pTWV228 | iSD, *Ec*RseP, H86A | This study |
| pYH878 | pTWV228 | iSD, *Ec*RseP, H87A | This study |
| pYH879 | pTWV228 | iSD, *Ec*RseP, H86A/H87A | This study |
| pKK49 | pUC118 | nSD, *Ec*RseP-HM | (*62*) |
| pKA52 | pUC118 | nSD, *Ec*RseP-HM, E23Q | (*24*) |
| pKA81 | pUC118 | nSD, *Ec*RseP-HM, E23Q, Y69amber | (*24*) |

| | | | |
|---|---|---|---|
| pTM352 | pUC118 | nSD, *Ec*RseP-HM, Y378amber | This study |
| pTM382 | pUC118 | nSD, *Ec*RseP-HM, E23Q, Y378amber | This study |
| pTM101 | pTWV228 | nSD, *Ec*RseP-HM, Cys-less | (*21*) |
| pYGF13 | pTWV228 | nSD, *Ec*RseP-HM, L151P | (*29*) |
| pTM129 | pTWV228 | nSD, *Ec*RseP-HM, Cys-less, L151P | This study |
| pTM303 | pTWV228 | nSD, *Ec*RseP-HM, Cys-less, T350C | This study |
| pTM304 | pTWV228 | nSD, *Ec*RseP-HM, Cys-less, G351C | This study |
| pTM305 | pTWV228 | nSD, *Ec*RseP-HM, Cys-less, D352C | This study |
| pTM306 | pTWV228 | nSD, *Ec*RseP-HM, Cys-less, V353C | This study |
| pTM307 | pTWV228 | nSD, *Ec*RseP-HM, Cys-less, K354C | This study |
| pTM308 | pTWV228 | nSD, *Ec*RseP-HM, Cys-less, L355C | This study |
| pTM309 | pTWV228 | nSD, *Ec*RseP-HM, Cys-less, N356C | This study |
| pTM310 | pTWV228 | nSD, *Ec*RseP-HM, Cys-less, N357C | This study |
| pTM162 | pTWV228 | nSD, *Ec*RseP-HM, Cys-less, L358C | This study |
| pTM163 | pTWV228 | nSD, *Ec*RseP-HM, Cys-less, S359C | This study |
| pTM164 | pTWV228 | nSD, *Ec*RseP-HM, Cys-less, G360C | This study |
| pTM165 | pTWV228 | nSD, *Ec*RseP-HM, Cys-less, P361C | This study |
| pTM166 | pTWV228 | nSD, *Ec*RseP-HM, Cys-less, I362C | This study |
| pTM167 | pTWV228 | nSD, *Ec*RseP-HM, Cys-less, S363C | This study |
| pTM168 | pTWV228 | nSD, *Ec*RseP-HM, Cys-less, I364C | This study |
| pTM169 | pTWV228 | nSD, *Ec*RseP-HM, Cys-less, A365C | This study |
| pTM170 | pTWV228 | nSD, *Ec*RseP-HM, Cys-less, K366C | This study |
| pTM171 | pTWV228 | nSD, *Ec*RseP-HM, Cys-less, G367C | This study |
| pTM172 | pTWV228 | nSD, *Ec*RseP-HM, Cys-less, A368C | This study |
| pTM173 | pTWV228 | nSD, *Ec*RseP-HM, Cys-less, G369C | This study |
| pTM174 | pTWV228 | nSD, *Ec*RseP-HM, Cys-less, M370C | This study |
| pTM175 | pTWV228 | nSD, *Ec*RseP-HM, Cys-less, T371C | This study |
| pTM176 | pTWV228 | nSD, *Ec*RseP-HM, Cys-less, A372C | This study |
| pTM177 | pTWV228 | nSD, *Ec*RseP-HM, Cys-less, E373C | This study |
| pTM178 | pTWV228 | nSD, *Ec*RseP-HM, Cys-less, L374C | This study |
| pTM179 | pTWV228 | nSD, *Ec*RseP-HM, Cys-less, G375C | This study |
| pTM180 | pTWV228 | nSD, *Ec*RseP-HM, Cys-less, V376C | This study |
| pTM181 | pTWV228 | nSD, *Ec*RseP-HM, Cys-less, V377C | This study |
| pTM182 | pTWV228 | nSD, *Ec*RseP-HM, Cys-less, Y378C | This study |
| pTM183 | pTWV228 | nSD, *Ec*RseP-HM, Cys-less, Y379C | This study |
| pTM184 | pTWV228 | nSD, *Ec*RseP-HM, Cys-less, L380C | This study |
| pTM185 | pTWV228 | nSD, *Ec*RseP-HM, Cys-less, P381C | This study |
| pYH9 | pSTD689 | nSD, *Ec*RseP-HM | (*19*) |
| pTM132 | pSTD689 | nSD, *Ec*RseP-HM, Cys-less | (*21*) |
| pTM311 | pSTD689 | nSD, *Ec*RseP-HM, Cys-less, T350C | This study |
| pTM312 | pSTD689 | nSD, *Ec*RseP-HM, Cys-less, G351C | This study |
| pTM313 | pSTD689 | nSD, *Ec*RseP-HM, Cys-less, D352C | This study |
| pTM314 | pSTD689 | nSD, *Ec*RseP-HM, Cys-less, V353C | This study |
| pTM315 | pSTD689 | nSD, *Ec*RseP-HM, Cys-less, K354C | This study |
| pTM316 | pSTD689 | nSD, *Ec*RseP-HM, Cys-less, L355C | This study |
| pTM317 | pSTD689 | nSD, *Ec*RseP-HM, Cys-less, N356C | This study |
| pTM318 | pSTD689 | nSD, *Ec*RseP-HM, Cys-less, N357C | This study |
| pTM186 | pSTD689 | nSD, *Ec*RseP-HM, Cys-less, L358C | This study |
| pTM187 | pSTD689 | nSD, *Ec*RseP-HM, Cys-less, S359C | This study |
| pTM188 | pSTD689 | nSD, *Ec*RseP-HM, Cys-less, G360C | This study |
| pTM189 | pSTD689 | nSD, *Ec*RseP-HM, Cys-less, P361C | This study |
| pTM190 | pSTD689 | nSD, *Ec*RseP-HM, Cys-less, I362C | This study |
| pTM191 | pSTD689 | nSD, *Ec*RseP-HM, Cys-less, S363C | This study |
| pTM192 | pSTD689 | nSD, *Ec*RseP-HM, Cys-less, I364C | This study |

| | | | |
|---|---|---|---|
| pTM193 | pSTD689 | nSD, *Ec*RseP-HM, Cys-less, A365C | This study |
| pTM194 | pSTD689 | nSD, *Ec*RseP-HM, Cys-less, K366C | This study |
| pTM195 | pSTD689 | nSD, *Ec*RseP-HM, Cys-less, G367C | This study |
| pTM196 | pSTD689 | nSD, *Ec*RseP-HM, Cys-less, A368C | This study |
| pTM197 | pSTD689 | nSD, *Ec*RseP-HM, Cys-less, G369C | This study |
| pTM198 | pSTD689 | nSD, *Ec*RseP-HM, Cys-less, M370C | This study |
| pTM199 | pSTD689 | nSD, *Ec*RseP-HM, Cys-less, T371C | This study |
| pTM200 | pSTD689 | nSD, *Ec*RseP-HM, Cys-less, A372C | This study |
| pTM201 | pSTD689 | nSD, *Ec*RseP-HM, Cys-less, E373C | This study |
| pTM202 | pSTD689 | nSD, *Ec*RseP-HM, Cys-less, L374C | This study |
| pTM203 | pSTD689 | nSD, *Ec*RseP-HM, Cys-less, G375C | This study |
| pTM204 | pSTD689 | nSD, *Ec*RseP-HM, Cys-less, V376C | This study |
| pTM205 | pSTD689 | nSD, *Ec*RseP-HM, Cys-less, V377C | This study |
| pTM206 | pSTD689 | nSD, *Ec*RseP-HM, Cys-less, Y378C | This study |
| pTM207 | pSTD689 | nSD, *Ec*RseP-HM, Cys-less, Y379C | This study |
| pTM208 | pSTD689 | nSD, *Ec*RseP-HM, Cys-less, L380C | This study |
| pTM209 | pSTD689 | nSD, *Ec*RseP-HM, Cys-less, P381C | This study |
| pTM913 | pTWV228 | nSD, *Ec*RseP-HM, Cys-less, D7C/K366C | This study |
| pTM916 | pTWV228 | nSD, *Ec*RseP-HM, Cys-less, S10C/G369C | This study |
| pTM919 | pTWV228 | nSD, *Ec*RseP-HM, Cys-less, D171C/S343C | This study |
| pTM922 | pTWV228 | nSD, *Ec*RseP-HM, Cys-less, I173C/K347C | This study |
| pTM925 | pTWV228 | nSD, *Ec*RseP-HM, Cys-less, E203C/K354C | This study |
| pTM928 | pTWV228 | nSD, *Ec*RseP-HM, Cys-less, D205C/S359C | This study |
| pKB4 | pTWV228 | iSD, *Ec*RseP, F426amber = ΔTM4 | This study |
| pKB5 | pTWV228 | iSD, *Ec*RseP, D446amber = ΔCTail | This study |
| pKB11 | pTWV228 | iSD, *Ec*RseP, G431A | This study |
| pKB12 | pTWV228 | iSD, *Ec*RseP, M439A | This study |
| pKB13 | pTWV228 | iSD, *Ec*RseP, A442S | This study |
| pKB14 | pTWV228 | iSD, *Ec*RseP, D446A | This study |
| pKB15 | pTWV228 | iSD, *Ec*RseP, R449A | This study |
| pKB28 | pTWV228 | iSD, *Ec*RseP, D446C | This study |
| pKB30 | pTWV228 | iSD, *Ec*RseP, D446F | This study |
| pKB31 | pTWV228 | iSD, *Ec*RseP, D446K | This study |
| pKB32 | pTWV228 | iSD, *Ec*RseP, D446N | This study |
| pKB38 | pTWV228 | iSD, *Ec*RseP, D446E | This study |
| pYH829 | pTWV228 | iSD, *Kk*RseP-TEV-$His_8$ | This study |
| pYH830 | pTWV228 | iSD, *Kk*RseP-TEV-$His_8$, H22F | This study |
| pYH831 | pTWV228 | iSD, *Kk*RseP-TEV-$His_8$, E23Q | This study |
| pYH832 | pTWV228 | iSD, *Kk*RseP-TEV-$His_8$, D395N | This study |
| pYH833 | pTWV228 | iSD, *Kk*RseP | This study |
| pYH834 | pTWV228 | iSD, *Kk*RseP, E23Q | This study |
| pYH838 | pTWV228 | iSD, *Kk*RseP(K54-PA14-H55) | This study |
| pYH839 | pTWV228 | iSD, *Kk*RseP(K54-PA14-H55), E23Q | This study |
| pYH866 | pTWV228 | iSD, *Kk*RseP(K54-PA14-H55), E163C | This study |
| pYH867 | pTWV228 | iSD, *Kk*RseP(K54-PA14-H55), F167C | This study |
| pYH868 | pTWV228 | iSD, *Kk*RseP(K54-PA14-H55), P136C | This study |
| pYH869 | pTWV228 | iSD, *Kk*RseP(K54-PA14-H55), I302C | This study |
| pYH18 | pTWV228 | HA-RseA148 | (*19*) |
| pYH20 | pTWV228 | HA-MBP-RseA(LY1)148 | (*19*) |
| pKA65 | pSTD689 | HA-MBP-RseA148 | (*24*) |
| pYH124 | pSTD689 | HA-MBP-RseA(LY1)148 | (*19*) |
| pYH817 | pSTD689 | HA-MBP-RseA(KkTM)148 | This study |

| | | | |
|---|---|---|---|
| pTM949 | pBAD33 | HA-MBP-RseA(LY1)148 | This study |

Amp$^{R}$, ampicillin-resistance marker; Spc$^{R}$, spectinomycin-resistance marker; Cm$^{R}$, chloramphenicol-resistance marker. nSD, native Shine Dalgarno; iSD, improved Shine Dalgarno.

**Table S3. Oligonucleotides used in biochemical analysis**

| Number | Name | Sequence |
|---|---|---|
| P1 | RV-N | TGTGGAATTGTGAGCGG |
| P2 | 120117RseP-His8_KpnI(-) | GGGGTACCTTAGTGATGGTGGTGATGGTGATGGTGACGGCCTTCG |
| P3 | 210128EcoSac-EcRseP(+) | GGAATTCGAGCTCGGTACAAGGAGGAAGAGCAAATGCTGAGTTTTCTCTGGGATTTGGC |
| P4 | M4C | TACGCCAGCTGGCGAAAGGG |
| P5 | 210310EcRseP-Hind(noTag)(-) | CTAGCCCAAGCTTCCTAACTAACTCTCATAACCGAGAGAAATCATTGAAAAG |
| P6 | 181212EcoSac_KkRseP(+) | GGAATTCGAGCTCGGTACAAGGAGGAAGAGCAAATGACTGATTTTTTATACAGTATTTTTGGC |
| P7 | 210121KkRseP-His_Bam(-) | CGCGGATCCTCAGTGATGGTGATGGTGATGGTGATGACTAG |
| P8 | 210407KkRseP(noTag)Hind(-) | CTAGCCCAAGCTTCATTGGTTTATTCGGTTTATGTCG |
| P9 | 210421Kk(K54-PA14-H55)_ins(+) | GGACGGACCAATAAGGAAGGTGGCGTTGCCATGCCAGGTGCCGAAGATGATGTGGTGCACGGCACTGAGTTT |
| P10 | 210421Kk(K54-PA14-H55)_ins(-) | AAACTCAGTGCCGTGCACCACATCATCTTCGGCACCTGGCATGGCAACGCCACCTTCCTTATTGGTCCGTCC |
| P11 | 210421InF_KkRseP-H55_vec(+) | CACGGCACTGAGTTTGTTATTGC |
| P12 | 210421InF_KkRseP-K54_vec(-) | CTTATTGGTCCGTCCCCAGATAGG |
| P13 | 210121Sal-KkTMv2-Pst(+) | TCGACCCATTCAAGCAATTAGCAGGCTATGCTATCGCTGCCTCAGTGGCCTTGGTGGTACTTTTCAACGTGGGACTGCA |
| P14 | 210121Sal-KkTMv2-Pst(-) | GTCCCACGTTGAAAAGTACCACCAAGGCCACTGAGGCAGCGATAGCATAGCCTGCTAATTGCTTGAATGGG |

**Table S4. Crystallographic analysis of PDZ-C fragments**

| Data sets | *Ec*PDZ-C:12C7 Fab | *Kk*PDZ-C |
|---|---|---|
| **Data Collection** | | |
| X-ray source | PF BL-17A | PF BL-17A |
| Wavelength (Å) | 0.9800 | 0.9800 |
| Space group | $P2_12_12_1$ | $P2_12_12_1$ |
| Cell dimensions | | |
| *a, b, c* (Å) | 58.10, 77.27, 283.63 | 40.67, 42.41, 97.95 |
| *α, β, γ* (°) | 90, 90, 90 | 90, 90, 90 |
| No. of moles or complexes / a.s.u. | 2 | 2 |
| Resolution limits (Å) | 49.50-3.20 (3.42-3.20) | 42.41-1.15 (1.17-1.15) |
| Unique reflection | 21,990 (3,909) | 60,798 (2,913) |
| Completeness (%) | 99.9 (100) | 99.6 (99.0) |
| Redundancy | 6.8 (7.1) | 6.4 (6.6) |
| $I/\sigma(I)$ | 7.5 (1.1) | 17.1 (2.2) |
| $R_{p.i.m.}$ | 0.105 (0.447) | 0.017 (0.350) |
| CC (1/2) | 0.992 (0.738) | 0.999 (0.794) |
| **Refinement** | | |
| Resolution limits (Å) | 49.50-3.20 (3.27-3.20) | 31.29-1.15 (1.16-1.15) |
| $R_{work}$[a] | 0.2432 (0.3944) | 0.1962 (0.3056) |
| $R_{free}$[b] | 0.2886 (0.4315) | 0.2057 (0.2759) |
| No. of non-H atoms | 7811 | 1582 |
| Average *B*-factor ($Å^2$) | 65.06 | 21.85 |
| RMSD from ideality | | |
| Bond length (Å) | 0.002 | 0.006 |
| Bond angle (°) | 0.46 | 0.82 |
| Ramachandran plot | | |
| Favored (%) | 93.04 | 97.65 |
| Outlier (%) | 0.30 | 0 |
| PDB code | 7W71 | 7W70 |

Values in parentheses are for highest-resolution shell.

[a] $R_{work}$ is the crystallographic *R*-factor calculated for the working set consisting of 95% of reflections used for the refinement.

[b] $R_{free}$ is the crystallographic *R*-factor calculated for the test set consisting of 5% of reflections excluded from the refinement.

**Table S5. Data collection statistics for the full-length RsePs**

| Data sets | *Ec*RseP (native) | *Ec*RseP (Zn) | *Kk*RseP (#1) | *Kk*RseP (#2) |
|---|---|---|---|---|
| X-ray source | SPring-8 BL32XU | SPring-8 BL32XU | SPring-8 BL32XU | SPring-8 BL32XU |
| Wavelength (Å) | 1.0000 | 1.2800 | 0.9700 | 0.9700 |
| Space group | *P*1 | *P*1 | *P*1 | $P2_1$ |
| Cell dimensions | | | | |
| *a, b, c* (Å) | 47.34, 56.12, 69.67 | 47.52, 56.40, 69.79 | 44.56, 49.78. 76.08 | 46.27, 40.80, 160.24 |
| *α, β, γ* (°) | 68.2, 74.6, 69.3 | 68.4, 74.6, 69.3 | 86.9, 79.2, 82.2 | 90, 91.6, 90 |
| No. of moles / a.s.u. | 1 | 1 | 1 | 1 |
| Resolution limits (Å) | 43.77-3.20 (3.31-3.20) | 47.14-3.71 (3.84-3.71) | 49.30-3.10 (3.21-3.10) | 40.80-3.15 (3.26-3.15) |
| Unique reflection | 10,129 (1,028) | 6,531 (637) | 11,493 (1,130) | 10,675 (1,055) |
| Completeness (%) | 99.7 (99.8) | 99.2 (99.5) | 99.8 (99.8) | 99.6 (99.8) |
| Redundancy | 8.4 (7.8) | 7.6 (7.9) | 135.8 (122.9) | 72.0 (66.3) |
| $I/\sigma(I)$ | 7.5 (1.1) | 7.7 (2.0) | 20.0 (1.4) | 13.8 (1.5) |
| $R_{p.i.m.}$ | 0.098 (1.347) | 0.100 (0.641) | 0.038 (1.601) | 0.081 (0.914) |
| CC (1/2) | 0.995 (0.417) | 0.984 (0.542) | 0.999 (0.821) | 0.999 (0.602) |

Values in parentheses are for highest-resolution shell.

**Table S6. Refinement statistics for the full-length RsePs**

| Data sets | *Ec*RseP (native) | *Kk*RseP (#1) | *Kk*RseP (#2) |
|---|---|---|---|
| Resolution limits (Å) | 43.76-3.20 (3.66-3.20) | 40.91-3.10 (3.24-3.10) | 40.55-3.15 (3.32-3.15) |
| $R_{work}$[a] | 0.2460 (0.2952) | 0.2564 (0.4159) | 0.2654 (0.3595) |
| $R_{free}$[b] | 0.3053 (0.3320) | 0.2987 (0.4787) | 0.2882 (0.4023) |
| No. of non-H atoms | 3478 | 3230 | 3252 |
| Protein | 3444 | 3197 | 3219 |
| $Zn^{2+}$ | 2 | 1 | 1 |
| Batimastat | 32 | 32 | 32 |
| Average *B*-factor (Å$^2$) | 94.52 | 92.30 | 91.49 |
| Protein | 94.60 | 92.50 | 91.35 |
| $Zn^{2+}$ | 83.44 | 64.45 | 110.74 |
| Batimastat | 87.24 | 73.57 | 105.10 |
| RMSD from ideality | | | |
| Bond length (Å) | 0.002 | 0.002 | 0.002 |
| Bond angle (°) | 0.48 | 0.50 | 0.44 |
| Ramachandran plot | | | |
| Favored (%) | 93.92 | 95.01 | 92.12 |
| Outlier (%) | 0 | 0 | 0.74 |
| PDB code | 7W6X | 7W6Y | 7W6Z |

Values in parentheses are for highest-resolution shell.

[a] $R_{work}$ is the crystallographic *R*-factor calculated for the working set consisting of 95% of reflections used for the refinement.

[b] $R_{free}$ is the crystallographic *R*-factor calculated for the test set consisting of 5% of reflections excluded from the refinement.
